# Exploring the functions of JAKMIP1 in neuronal IL-6/STAT3 signaling and its relevance to chromosome 15q-duplication syndrome

**DOI:** 10.1101/2025.10.21.683757

**Authors:** Josan G. Martin, Emily-Rose Martin, Natsuki Takamura, Charli E. Harlow, Rosemary A. Bamford, Zoë K.K. Mia, Rebecca G. Smith, Noel G. Morgan, Satomi Inaba-Inoue, Jonathan Mill, Deepak P. Srivastava, Helen R. Dawe, John K. Chilton, Mark A. Russell, Asami Oguro-Ando

## Abstract

Growing evidence supports neuroinflammation as a risk factor for neurodevelopmental and psychiatric disorders. As a classical pro-inflammatory cytokine, Interleukin 6 (IL-6) and its downstream JAK/STAT3 signaling have been associated with autism spectrum disorder (ASD)-related phenotypes. To better understand molecular factors that modify neuronal cytokine responses in ASD, we investigated potential roles for the *JAKMIP1* gene in regulating IL-6/STAT3 signaling. *JAKMIP1* encodes a JAK-interacting protein, and its dysregulation is linked to chromosome 15q-duplication syndrome (Dup15q; a form of syndromic ASD). We observe that JAKMIP1 deficiency impairs IL-6/STAT3 signaling and IL-6-induced neuritogenesis in SH-SY5Y cells; and discover that JAKMIP1 may regulate *STAT3* expression via its C-terminus, which exhibits nucleoplasmic localization and interacts with multiple regulators of transcription, splicing and RNA stability. Additionally, we find that IL-6/STAT3 signaling is altered in Dup15q hiPSCs-derived cortical neurons, which display heightened responsiveness to IL-6; though it remains unclear whether and how JAKMIP1 contributes to this. Overall, our findings identify JAKMIP1 as a modulator of neuronal IL-6/STAT3 signaling and support that ASD-linked genetic variants can alter the inflammatory landscape of ASD.

## Introduction

Beyond its classical role in the acute phase response (Tanaka, Narazaki and Kishimoto, 2014), Interleukin (IL)-6 (IL-6) plays a wide variety of roles in the nervous system. IL-6 has emerged as a regulator of neural cell differentiation (Islam *et al*., 2009), glial cell function (Recasens *et al*., 2021), neurite outgrowth (Ihara *et al*., 1996; Wu and Bradshaw, 1996) and synaptic plasticity (Wei *et al*., 2011; Esquivel-Rendón *et al*., 2019). Notably, neuroinflammation and more generally, imbalances in cytokines, has been associated with various neurodevelopmental or psychiatric disorders (Vargas *et al*., 2005; Warre-Cornish *et al*., 2020; Han *et al*., 2021) and neurodegenerative diseases (Guzman-Martinez *et al*., 2019). IL-6 signaling and its downstream effectors (e.g., IL-17A) have also recently been demonstrated to be causal for the manifestation of autism spectrum disorder (ASD)-like behavior abnormalities in rodent models of maternal immune activation (Parker-Athill and Tan, 2011; Zawadzka, Cieślik and Adamczyk, 2021). Hence, it is clear that cytokines act upon and modify the function of neural cells, and careful finetuning of their signaling during development governs changes to behavior and cognition.

To expand upon our understanding of the regulatory mechanisms that control neural responses to cytokines, and how these could be altered in neurological disorders such as ASD, we turned our attention to Janus Kinase and Microtubule-Interacting Protein 1 (JAKMIP1). In the brain, JAKMIP1 (also known as JAMIP1, MARLIN1, or GABA_B_RBP), is primarily expressed in neurons (Costa *et al*., 2007; Vidal *et al*., 2009) and is a protein that possesses distinct functional domains (Steindler *et al*., 2004; Costa *et al*., 2007; Vidal *et al*., 2012; Berg *et al*., 2015) (these are summarized in Fig. S1). *JAKMIP1* expression was initially reported to be altered in lymphoblastoid cell lines derived from cases of fragile X syndrome (FXS) and chromosome 15q-duplication syndrome (Dup15q), some of the most common types of syndromic ASD (de la Torre-Ubieta *et al*., 2016). Additionally, *JAKMIP1* expression was found to be downregulated in cellular and mouse models of FXS and Dup15q (Nishimura *et al*., 2007). Other studies indicate that JAKMIP1 is important for normal development, as *Jakmip1*^−/−^ mice exhibit altered glutamatergic neurotransmission and ASD-associated behaviors such as changes in social preference and repetitive jumping (Berg *et al*., 2015), and *Jakmip1* silencing impairs the radial migration of cortical neurons (Vidal *et al*., 2012). Furthermore, several genetic and epigenetic studies have linked likely pathogenic mutations and differential methylation sites in the *JAKMIP1* gene to cases of ASD (Hedges *et al*., 2012; Yehia *et al*., 2020), other genetic syndromes that display ASD-associated phenotypes (Yang *et al*., 2016; Helbig *et al*., 2020), schizophrenia (Mill *et al*., 2008), alcohol consumption (Cervera-Juanes *et al*., 2017) and Alzheimer’s disease (Li, Sun and Wang, 2020).

Yet, the exact physiological roles and molecular functions of JAKMIP1, and how its dysregulation can contribute to neurological disorders, remains unclear. Importantly, the distinct domains of JAKMIP1 outlined in Fig. S1 places it in a unique position to link receptor transport, microtubule-based functions and local synaptic protein translation together in neurons – processes integral to proper neuronal function. However, it is unknown whether JAKMIP1 interactions with the Janus Kinase (JAK) proteins (specifically JAK1 and TYK2) (Steindler *et al*., 2004) allow it to modulate neuronal cytokine signaling as well, especially since these JAKMIP1-JAK1 interactions have not been confirmed in neural cells. JAKMIP1 dysregulation could have important implications for the neuroinflammation observed in ASD, as JAKMIP1 may be responsible for transferring cytokine signals to the neuronal microtubule cytoskeleton or synaptic architecture. Importantly, we have recently discovered that increased expression of *Cytoplasmic FMRP-Interacting Protein 1* (*CYFIP1*), a gene implicated in Dup15q and other neuropsychiatric disorders, can alter IL-6/STAT3 signaling and IL-6-induced neurite outgrowth as well (Martin *et al*., 2026). Thus, we sought to further our understanding of the molecular functions of JAKMIP1 and test whether altered expression of JAKMIP1, as observed in models of syndromic ASD such as Dup15q (Nishimura *et al*., 2007), could lead to changes in how neuronal cells respond to cytokines. This would add to our understanding of neural factors that modify inflammatory signaling during development and how their dysregulation can contribute to the susceptibility or pathogenesis of neurodevelopmental and psychiatric disorders.

Here, we explore potential roles for JAKMIP1 in neuronal cytokine signaling. We provide evidence to support a novel role for JAKMIP1 in neuronal IL-6 signaling – showing that *JAKMIP1*^−/−^ human neuroblastoma SH-SY5Y cells exhibit reduced levels of the downstream transcription factor *Signal Transducer and Activator of Transcription 3* (*STAT3*) expression, impaired IL-6-induced STAT3 transcriptional activity and attenuated IL-6-induced neuritogenesis. We further explore the impact of JAKMIP1 deficiency by transcriptomic profiling and identify additional cytokine signaling-related genes that are differentially expressed in *JAKMIP1*^−/−^ cells. Additionally, we use tandem immunoprecipitation mass spectrometry (IP-MS) to characterize the JAKMIP1 interactome and find potential mechanisms by which JAKMIP1 may control *STAT3* expression. However, when testing if altered JAKMIP1 expression could modify IL-6/STAT3 signaling in a Dup15q human induced pluripotent stem cell (hiPSC) line and hiPSC-derived cortical neurons (iNeurons), we found that *JAKMIP1* expression did not always correlate with *CYFIP1* or *STAT3* expression. Further, we also discover that Dup15q hiPSCs and iNeurons appear to demonstrate enhanced IL-6-induced STAT3 responses, in contrast to our SH-SY5Y cell experiments. Taken together, we hypothesize that the complex genetic background of Dup15q creates a change in the landscape of cytokine signaling that prevents JAKMIP1 (and its interactome) from exerting its normal influence over *STAT3* expression and activity. Our findings provide new insights into the impacts that the complex genetic landscape of ASD can have on how neurons respond to cytokine stimuli.

## Results

### JAKMIP1 deficiency modifies neuronal responses to IL-6

To investigate if JAKMIP1 may be involved in neuronal cytokine responses, we examined whether JAKMIP1 can form a component of cytokine receptor complexes. Its interacting partners, JAK1 and TYK2, are known to associate with multiple cytokine receptors – including those that bind to the IL-6 family of cytokines (Fig. 1A) (Sims, 2020). Through immunocytochemical staining, we observe that JAKMIP1 co-localizes with the IL-6Rα and GP130 subunits of the IL-6 Receptor (IL-6R) complex as well as associated JAK1 (Fig. 1B), indicating that JAKMIP1 may form a novel component of the IL-6R complex in neuronal cells. This co-localization has also been validated in primary chick dorsal root ganglion neurons (Fig. S2).

**Figure 1.**
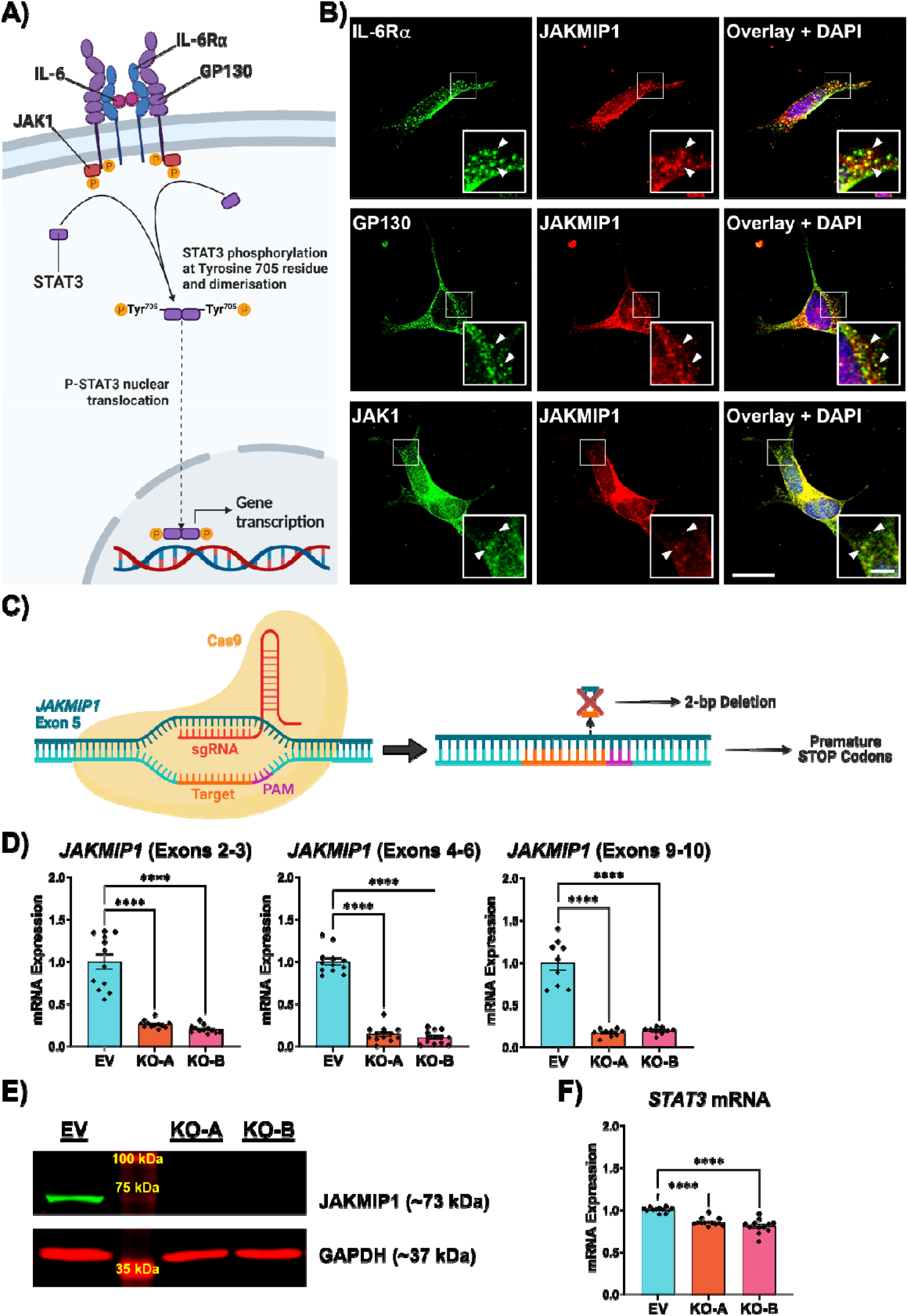
JAKMIP1 co-localizes with IL-6R complexes and JAKMIP1-deficient SH-SY5Y cells display reduced STAT3 expression. **A)** Key events of canonical IL-6 signaling via the IL-6 Receptor (IL-6R) complex (composed of IL-6Rα, GP130 and JAK1) and downstream STAT3. **B)** Fluorescent micrographs of SH-SY5Y cells co-stained for JAKMIP1 (red) and components of the IL-6R complex (IL-6Rα, GP130 or JAK1; green). Nuclei were counterstained with 4’,6-Diamidino-2-Phenylindole (DAPI; blue). Scale Bar = 20 μm. Inset scale bar = 5 μm. **C)** Schematic demonstrating the use of CRISPR-Cas9 gene editing technology to generate a 2 base-pair (bp) deletion in exon 5 of the *JAKMIP1* gene, leading to premature stop codons. **D)** Relative mRNA expression of *JAKMIP1* across multiple exon junctions (upstream, overlying, and downstream of the CRISPR-Cas9-mediated deletion) measured by qRT-PCR. One way ANOVA with Tukey’s HSD test for multiple comparisons. **E)** Loss of JAKMIP1 protein expression in both KO-A and KO-B cell lines confirmed via Western blotting. **F)** Relative mRNA expression of *STAT3* in EV, KO-A and KO-B cell lines measured by qRT-PCR. Statistics as in D). Values presented as mean ± SEM of N = 3 independent experiments. ****P < 0.0001.

To test whether JAKMIP1 can modify neuronal IL-6 responses, we generated two *JAKMIP1*^−/−^ SH-SY5Y cell lines by CRISPR-Cas9 gene-editing (referred to as KO-A and KO-B), (Fig. 1, C-E; and Fig. S3) and assayed *STAT3* expression, STAT3 phosphorylation dynamics and STAT3 transcriptional activity following IL-6 stimulation. An “empty vector” (EV) cell line that was transfected with the same plasmid construct for the CRISPR-Cas9 gene-editing process without a guide RNA was used as the control cell line for all experiments using the KO-A and KO-B lines.

We find that *JAKMIP1*^−/−^ SH-SY5Y cells display reduced levels of *STAT3* mRNA (Fig. 1F), suggesting that JAKMIP1 could regulate *STAT3* expression at the transcriptional level. To investigate whether *JAKMIP1*^−/−^ SH-SY5Y cells therefore respond differently to STAT3-agonistic cytokines, Western blotting was performed to measure IL-6-induced phosphorylation at the Tyr^705^ residue (representative of STAT3 activation). We observed ∼20% reduced STAT3 protein expression in *JAKMIP1*^−/−^ SH-SY5Y cells (Fig. 2, A and B), which was accompanied by reduced total levels of STAT3 (Tyr^705^) phosphorylation (Fig. 2C). Notably, levels of STAT3 (Tyr^705^) phosphorylation and its fold induction following IL-6 treatment are not significantly different between the EV and *JAKMIP1*^−/−^ SH-SY5Y cells when normalized to total levels of STAT3 expression (Fig. 2D). This indicates that JAKMIP1 deficiency does not alter (Tyr^705^) phosphorylation kinetics but mainly affects *STAT3* expression.

**Figure 2.**
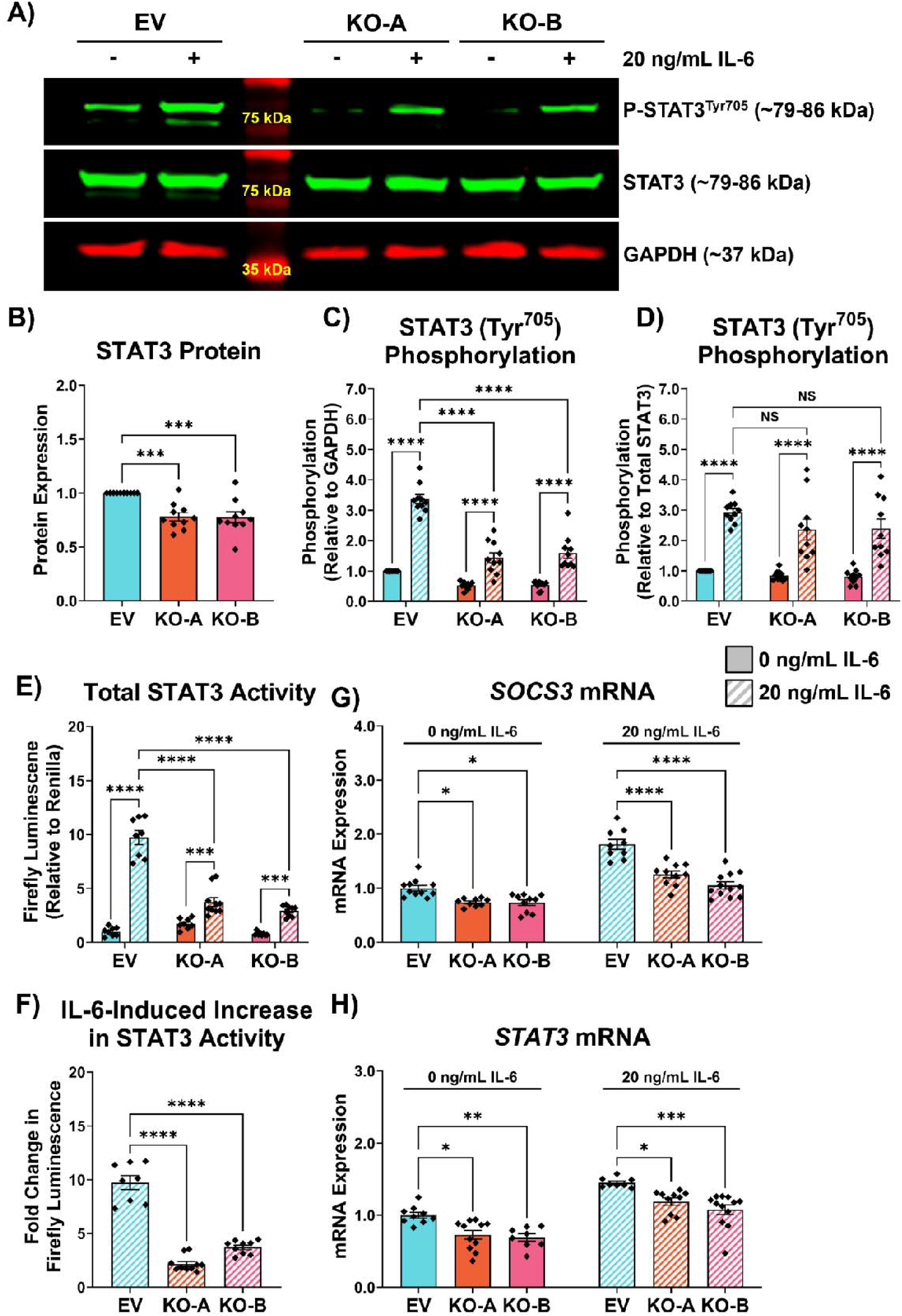
JAKMIP1-deficient SH-SY5Y cells display reduced IL-6-induced STAT3 (Tyr^705^) phosphorylation and attenuated STAT3 transcriptional activity. **A)** EV, KO-A and KO-B cell lines were treated with 20 ng/mL IL-6 or complete medium for 30 minutes prior to lysis for protein extraction. Western blotting was performed to measure STAT3 expression and STAT3 (Tyr^705^) phosphorylation. **B)** Densitometric analysis of total STAT3 protein expression relative to the GAPDH loading control. One way ANOVA with Tukey’s HSD test for multiple comparisons. **C)** Densitometric analysis of STAT3 phosphorylation at the Tyr^705^ residue (P-STAT3^Tyr705^) relative to the GAPDH loading control. Two-way ANOVA with Tukey’s HSD test for multiple comparisons. **D)** Densitometric analysis of P-STAT3^Tyr705^ relative to total STAT3 protein. Statistics as in C). **E)** Total transcriptional activity at a STAT3-regulated promoter in EV, KO-A and KO-B SH-SY5Y cell lines measured by Firefly luminescence in a DLR™ assay. SH-SY5Y cells were stimulated with 20 ng/mL IL-6 or complete medium for 18 hours post-transfection with Cignal Reporter plasmids. Firefly luminescence intensity is expressed as a ratio to Renilla luminescence intensity to adjust for transfection efficiency. Statistics as in C). **F)** Fold change in transcriptional activity from E) following IL-6 treatment in EV, KO-A and KO-B SH-SY5Y cell lines. **G)** Relative mRNA expression of *SOCS3* in EV, KO-A and KO-B cell lines exposed to complete medium with or without 20 ng/mL IL-6 for 4 hours, measured by qRT-PCR. Statistics as in C). **H)** Relative mRNA expression of *STAT3* in EV, KO-A and KO-B cell lines exposed to complete medium with or without 20 ng/mL IL-6 for 4 hours, measured by qRT-PCR. Statistics as in C). Values presented as mean ± SEM of N = 3-4 independent experiments. NS – not significant, *P < 0.05, **P < 0.01; ***P < 0.001, ****P < 0.0001.

To evaluate how the reduction in STAT3 (Tyr^705^) phosphorylation, and therefore total levels of activated STAT3, affect transcriptional activity, we utilized a dual-luciferase® reporter (DLR™) assay. This assay places the expression of the Firefly luciferase under the control of a STAT3 response element. Complementing our Western blotting data showing reduced levels of activated STAT3 following JAKMIP1 deficiency, STAT3 transcriptional activity (i.e., Firefly luminescence) was reduced in both *JAKMIP1*^−/−^ SH-SY5Y cell lines following IL-6 treatment relative to the EV cells (Fig. 2E). This attenuated transcriptional response becomes clearer when comparing the fold change in Firefly luminescence between IL-6-treated and non-IL-6-treated cells (Fig. 2F). To validate the DLR™ assay findings, we measured the expression of STAT3-responsive genes, *SOCS3* and *STAT3* (Narimatsu *et al*., 2001; Sims, 2020)) in *JAKMIP1*^−/−^ SH-SY5Y cells stimulated with IL-6. These genes were separately confirmed to be induced downstream of IL-6 signaling in SH-SY5Y cells (Fig. S4). In agreement with the DLR™ assay, IL-6-induced expression of *SOCS3* and *STAT3* mRNA were reduced in *JAKMIP1*^−/−^ SH-SY5Y cells (Fig. 2, G and H). This provides further evidence that JAKMIP1 may play a novel role in modulating transcriptional responses to cytokines such as IL-6.

Previous studies have reported that IL-6/STAT3 signaling promotes neurite outgrowth and axon extension (Wu and Bradshaw, 1996; Schäfer *et al*., 1999; Yang and Tang, 2017); and that STAT3 is responsible for the neurite outgrowth-promoting effects of other cytokines and neurotrophic factors (Bella *et al*., 2006; Chen *et al*., 2016). Given that JAKMIP1 can modulate IL-6-induced STAT3 signaling, we hypothesized that JAKMIP1 could control STAT3-mediated neurite outgrowth. In agreement with a previous finding, we observe that *JAKMIP1*^−/−^ SH-SY5Y cells show impairments, albeit subtle, in neurite outgrowth relative to the EV control (Fig. S5). A rescue experiment performed in parallel by selective overexpression of the N- and C-terminal domains of JAKMIP1 suggests that the microtubule-binding functions of JAKMIP1 N-terminus may be primarily responsible for its role in neurite outgrowth regulation (Fig. S6). However, when the SH-SY5Y cells were differentiated in the presence of IL-6, JAKMIP1 deficiency exerted a more pronounced effect. In EV control cells, IL-6 enhances neurite outgrowth (Fig. 3, A-C). On the other hand, this increase in neurite lengths following IL-6 stimulation is attenuated in the *JAKMIP1*^−/−^ cells (Fig. 3, A-C), in a similar manner to the attenuation of Firefly luminescence observed in the DLR™ assay. This suggests that JAKMIP1 may also contribute to neurite outgrowth via modulating IL-6/JAK1/STAT3 signaling and it is the reduced STAT3 activity found in both *JAKMIP1*^−/−^ SH-SY5Y cell lines that impaired the IL-6-stimulated neuritogenesis.

**Figure 3.**
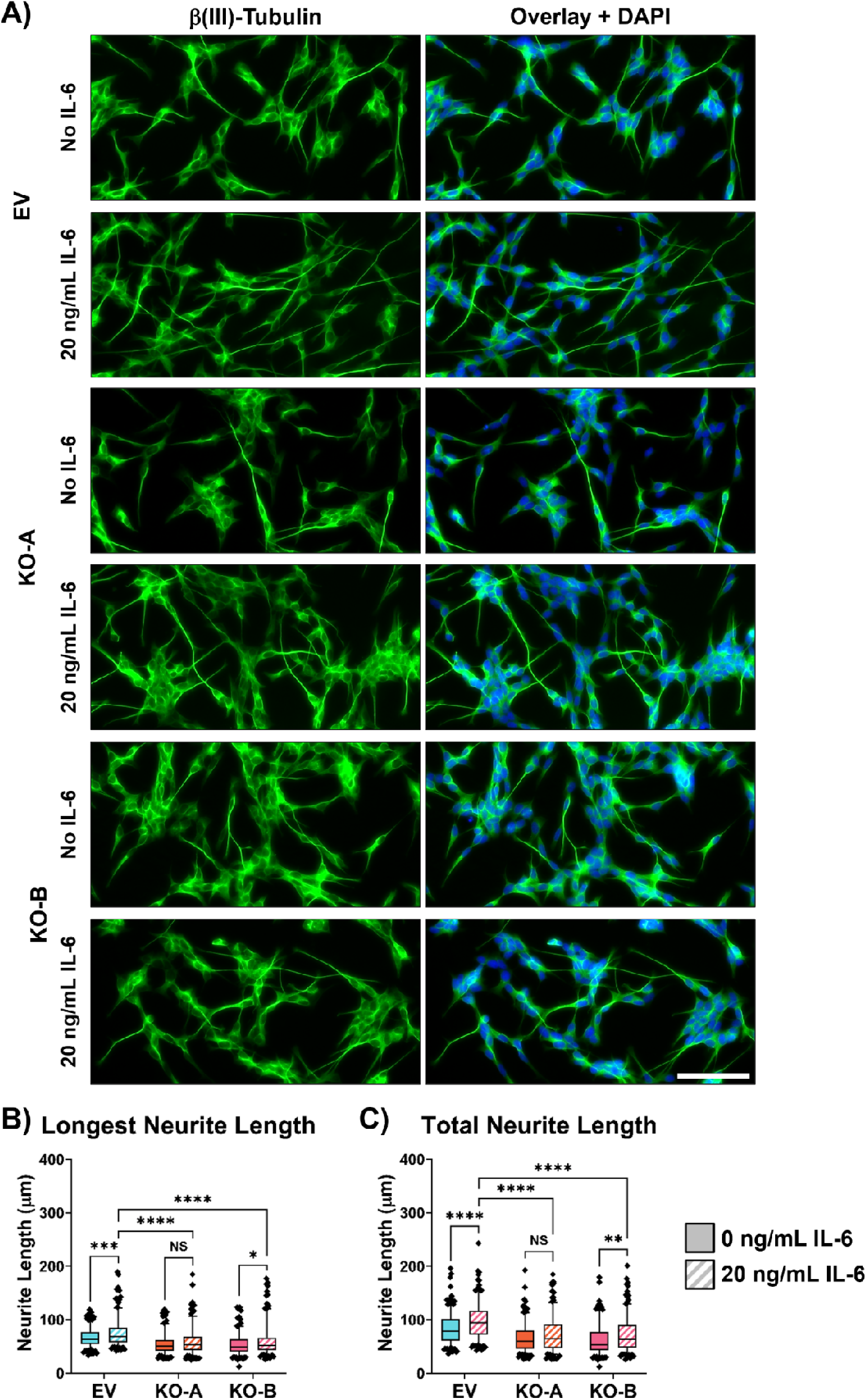
*JAKMIP1*^−/−^ cells display impaired IL-6-induced neurite outgrowth. **A)** EV, KO-A and KO-B SH-SY5Y cells were seeded onto poly-D-Lysine-coated glass coverslips differentiated with or without the presence of 20 ng/mL IL-6. SH-SY5Y cells were then fixed and stained for β(III)-Tubulin (TUBB3; green). Scale bar = 100 μm. **B)** Measurement of the length of the longest neurite (LNL) produced by individual SH-SY5Y cells in part A). Two-way ANOVA with Tukey’s HSD test for multiple comparisons. **C)** Measurement of the sum of all neurites (TNL) produced by individual SH-SY5Y cells in part A). Statistics as in B). Boxplots display distribution of neurite lengths, with whiskers extending to 5^th^ and 95^th^ percentiles of the data, N = 3 independent experiments. NS – not significant, *P < 0.05, **P < 0.01, ***P < 0.001, ****P < 0.0001.

### Transcriptomic profiling of *JAKMIP1*^−/−^ SH-SY5Y cells expands the repertoire of potential JAKMIP1-regulated functions in neuronal cells and reveals a role for JAKMIP1 in regulating the expression of ASD risk genes

Considering that JAKMIP1 appears to modulate *STAT3* transcription, we hypothesized that JAKMIP1 deficiency could alter the transcription of other genes during neuronal development, either directly or indirectly (as STAT3 itself is a transcription factor). Furthermore, as high degrees of crosstalk are known to exist among cytokine signaling pathways, it is possible that components of other cytokine signaling cascades are differentially expressed in *JAKMIP1*^−/−^ SH-SY5Y cells.

To explore this, we used RNA sequencing (RNA-seq) technology to profile differentially expressed genes (DEGs) in longer-term differentiated EV control and *JAKMIP1*^−/−^ SH-SY5Y cells that display a more mature, neuron-like phenotype. Principal component analyses (PCA) of the RNA-seq data effectively segregated the EV samples from both KO-A and KO-B samples (Fig. 4A), indicating that a large proportion of the changes to the transcriptome of *JAKMIP1*^−/−^ SH-SY5Y cells can be attributed to loss of JAKMIP1 expression. However, we observed a high degree of dissimilarity in the DEGs identified in transcriptome of KO-A and KO-B samples relative to the EV control (Fig. S7, A and B). Thus, to obtain a list of DEGs that are most likely to be specific to the effects of JAKMIP1 deficiency in differentiated SH-SY5Y cells, we compared the two separate differential gene expression analyses (EV against KO-A, and EV against KO-B) to identify a subset of statistically significant DEGs (adjusted P-value < 0.05) consistent between both *JAKMIP1*^−/−^ cell lines (termed both “overlapping” and “significant” DEGs) (Fig. 4B). This comparison revealed 962 overlapping significant DEGs common to both *JAKMIP1*^−/−^ SH-SY5Y cell lines (Fig. 4B), which mostly match in their direction of differential expression compared to the EV control (Fig. 4C). 14 of these genes did not show the same direction of effect, *AC090164.5*, *AIF1L*, *ATF3*, *CBLN4*, *COP1*, *DRGX*, *NPPC*, *OXTR*, *RAX*, *RGS4*, *RGS8*, *SEMA3E*, *SYT7* and *ZNF488*, being upregulated in one *JAKMIP1*^−/−^ cell line and downregulated in the other and hence were removed. This resulted in a final list of 948 overlapping and significant DEGs common to both *JAKMIP1*^−/−^ cell lines (Table S1).

**Figure 4.**
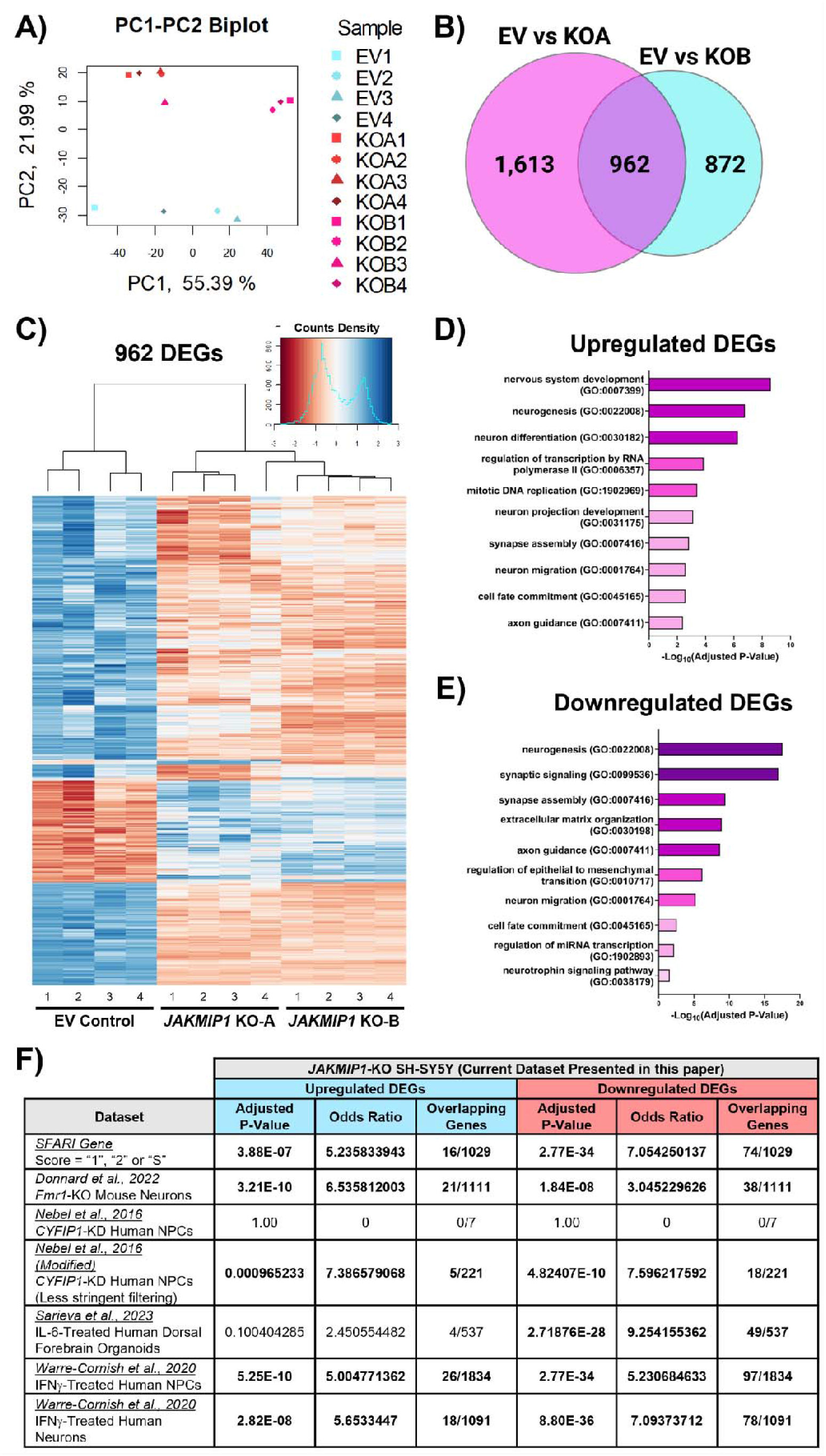
Identifying commonly differentially expressed genes in both *JAKMIP1*^−/−^ SH-SY5Y cell lines compared to the EV control, and comparing these to ASD-relevant datasets. **A)** Biplot of principal component (PC) 1 and PC2 following PC analysis to visualize the sources of variation in the transcriptomic data. The percentage of variance in the dataset accounted for by the corresponding PC is specified along the x- and y-axes. **B)** Venn diagram of the significant DEGs (adjusted P-value ≤ 0.05) across the different cell line comparisons. 962 genes are commonly altered in expression in both KO-A and KO-B compared to EV (i.e., the “overlapping” DEGs). **C)** Heatmap of the 962 overlapping and significant DEGs from part A). Clustering by Ward’s linkage was performed on regularized log counts of each DEG following conversion to z-scores. **D-E)** Results of GO analysis performed on the separated D) up- and E) downregulated DEGs with *g:Profiler* to identify enrichment of GO Biological Process 2021 terms. 10 terms of interest are reported with the adjusted P-value following Benjamini-Hochberg FDR correction for multiple testing. Terms are sorted by adjusted P-value; longer and darker bars reflect smaller P-values. **F)** Results of gene-set over-representation analyses (GSOA) to test for enrichment of ASD-associated genes, FMRP/CYFIP1-regulated genes and those altered by maternal immune activation in the list of DEGs obtained from *JAKMIP1*^−/−^ SH-SY5Y cells. Enrichment analyses were conducted with Fisher’s exact tests, and P-values were adjusted with Benjamini-Hochberg FDR correction to account for multiple testing. The “Overlapping Genes” columns refer to the genes present in a particular dataset (either an ASD-associated SFARI gene or a gene differentially expressed following altered expression of FXS- and Dup15q-linked genes *Fragile X Messenger Ribonucleoprotein* (*FMR1*) and *CYFIP1*; or genes responsive to IL-6 or IFN-γ treatment) that is also differentially expressed in the *JAKMIP1*^−/−^ SH-SY5Y cells. Comparisons with an FDR-adjusted P-value that meets the threshold for statistical significance have been bolded.

The 948 DEGs were then subjected to gene ontology (GO) pathway enrichment analyses to infer cellular processes potentially regulated by JAKMIP1. GO analysis indicates that the up- and downregulated DEGs converge on common biological processes important for neurodevelopment, such as “neurogenesis”, “synapse assembly”, “neuron migration”, “axon guidance” and “cell fate commitment” (Fig. 4, D and E; Table S2 and S3). However, there are also some pathways that are more specific to the up- and downregulated DEGs. For example, upregulated DEGs appear to be more enriched for functions in transcription or DNA replication (Fig. 4D). In contrast, the downregulated DEGs are more enriched for functions in extracellular matrix (ECM) organization, neurotrophin/growth factor/cytokine pathways, and an array of intracellular signaling cascades (Fig. 4E). Note that GO Biological Process 2021 enrichment analyses of the downregulated DEGs identified several cytokine/inflammation/immune function-related terms as well (Fig. S8). In particular, the RNA-seq data suggests that JAKMIP1 deficiency may alter neuronal responses to other cytokines that activate Transforming Growth Factor (TGF)-β/SMAD or Nuclear Factor Kappa B (NF-κB) signaling (Fig. S8). Altogether, these findings provide further evidence that JAKMIP1 may play novel roles in modulating neuronal cytokine signaling, among other potentially important functions such as neuronal morphogenesis, synaptic function and ECM organization. A selection of DEGs were chosen for follow-up independent validation by qRT-PCR (Fig. S9, A to F). qRT-PCR confirmed that components of the IL-6/JAK1/STAT3 pathway (*JAK1*, *STAT3* and *SOCS3*) are differentially expressed in differentiated *JAKMIP1*^−/−^ SH-SY5Y cells (Fig. S9B), as well as a range of neuronal cell adhesion molecules (Fig. S9C), axon guidance ligands and receptors (Fig. S9D), ECM components (Fig. S9E), as well as synaptic receptors and ion channels (Fig. S9F).

As previously mentioned, JAKMIP1 dysregulation has been linked to ASD and ASD-associated behavior phenotypes (Nishimura *et al*., 2007; Berg *et al*., 2015). Although we have demonstrated that JAKMIP1 regulates the expression of genes involved in neurodevelopment and neuronal function, it is unclear which JAKMIP1-regulated DEGs may be important in the context of ASD. Therefore, we performed gene-set over-representation analyses (GSOA) examining whether the list of DEGs in *JAKMIP1*^−/−^ SH-SY5Y cells were enriched for established ASD candidate risk genes (curated by *SFARI Gene* (Abrahams *et al*., 2013)) and those genes differentially expressed in neural models of *FMR1* and *CYFIP1* disruption (to model FXS and Dup15q) (Nebel *et al*., 2016; Donnard, Shu and Garber, 2022). Furthermore, we also tested whether JAKMIP1 may regulate the expression of genes altered in human neural models that mimic maternal immune activation (Warre-Cornish *et al*., 2020; Sarieva *et al*., 2023). This would provide additional evidence that JAKMIP1 can modulate the expression of genes that form part of the transcriptional response of neurons to cytokines, and whether they may be directly relevant to maternal immune activation-associated ASD.

The GSOA indicate that ASD risk genes are disproportionately affected by JAKMIP1 deficiency, and that overlapping gene expression signature exist between the DEGs from *JAKMIP1*^−/−^ SH-SY5Y cells and the various models included in the analysis. This suggests that a subset of genes sensitive to *FMR1* or *CYFIP1* disruption are also controlled by JAKMIP1 and similarly, there are genes whose expression is regulated by JAKMIP1 which are responsive to IL-6 or IFN-γ (Fig. 4F). Altogether, these analyses further cement a link between JAKMIP1 and ASD and imply that JAKMIP1-regulated genes and pathways may be relevant to maternal immune activation-associated ASD, where inflammatory signaling is particularly relevant to its etiology.

### The JAKMIP1 C-terminus exhibits nuclear localization and may facilitate a role in transcriptional regulation

Having demonstrated that JAKMIP1 may regulate the expression of *STAT3* mRNA, we posited that this may be due to undiscovered functions of JAKMIP1 in transcriptional regulation. JAKMIP1 has been reported to exist in the nucleus associated with euchromatin (Vidal *et al*., 2009), suggesting a potential nuclear mechanism for gene expression control. We first confirmed the presence of nuclear JAKMIP1 by endogenous JAKMIP1 staining in SH-SY5Y cells (Fig. 5A).

**Figure 5.**
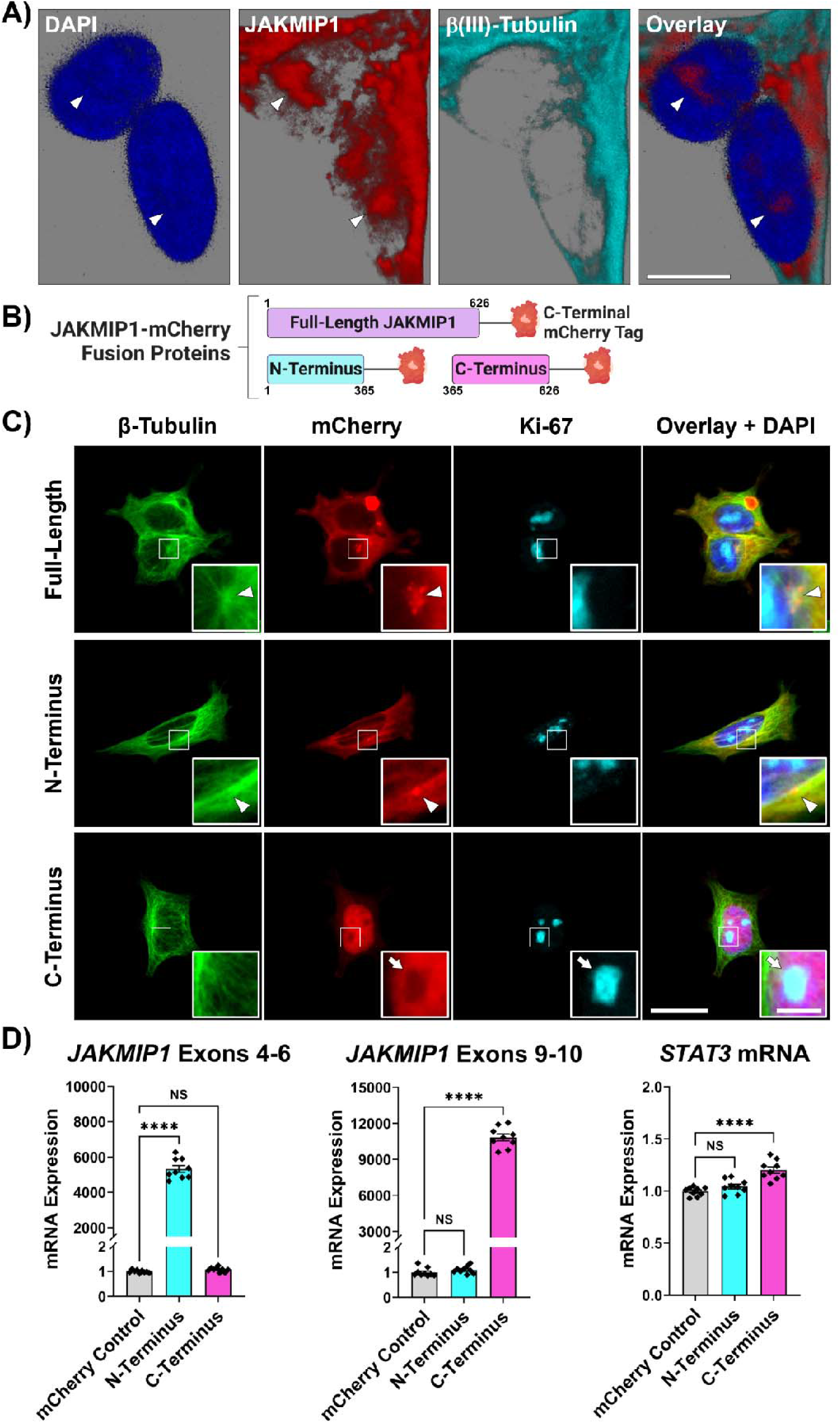
The JAKMIP1 C-terminus accumulates in the nucleoplasm and may control *STAT3* expression. **A)** Three-dimensional reconstruction of Z-stack imaging of SH-SY5Y cells stained for endogenous JAKMIP1 (red) and β(III)-Tubulin (TUBB3; cyan) using the LAS X software. Nuclei were counterstained with 4’,6-Diamidino-2-Phenylindole (DAPI; blue). Arrowheads indicate nuclear populations of JAKMIP1. Scale bar = 10 μm. **B)** Schematic of JAKMIP1-mCherry fusion proteins, including the full-length and truncated N- and C-termini domains. **C)** Fluorescence micrographs of HEK-293 cells transfected with plasmids encoding JAKMIP1 constructs in B), which were then fixed and stained for β-Tubulin (TUBB; green) and Ki-67 (cyan). Nuclei were counterstained with DAPI (blue). Scale bar = 20 μm. Arrowheads indicate juxtanuclear puncta (likely centrosomal localization) of the full-length and N-terminal JAKMIP1 constructs; arrow indicates non-nucleolar localization of the JAKMIP1 C-terminus construct. Inset scale bar = 5 μm. **D)** Relative mRNA expression of *JAKMIP1* (across multiple exon junctions) and *STAT3* in SH-SY5Y cells overexpressing the mCherry-tagged truncated JAKMIP1 constructs in B) relative to control SH-SY5Y cells overexpressing only mCherry, measured by qRT-PCR. One-way ANOVA with Tukey’s HSD test for multiple comparisons. Values presented as mean ± SEM of N = 3 independent experiments. NS – not significant, ****P < 0.0001.

Next, we sought to identify which domain of JAKMIP1 was responsible for this nuclear localization. Given the largely microtubular role of the JAKMIP1 N-terminus, we hypothesized that the C-terminus of JAKMIP1 could be responsible for this nuclear localization. To investigate this, we transfected HEK-293 cells with mCherry-tagged truncated N- and C-termini, as well as the full-length JAKMIP1 constructs (Fig. 5B). When observed, the N-terminus displays predominant cytoplasmic localization, similar to the full-length protein, and co-localizes with microtubules as expected (Fig. 5C). Furthermore, we find that overexpression of the full-length and N-terminus constructs often result in the formation of juxtanuclear puncta (Fig. 5C) probably in close proximity to the centrosome, the major microtubule-organizing center of the cell. In contrast, the JAKMIP1 C-terminus accumulates largely in the nucleus (Fig. 5C) although it also exhibits a degree of microtubule-association in some cells (Fig. S10). Importantly, the C-terminus is excluded from the nucleolar compartment, which is visible through Ki-67 staining in proliferating cells (Fig. 5C) (Sun and Kaufman, 2018). The observed localization patterns of these constructs were replicated in SH-SY5Y cells (Fig. S11) and in live cells (Fig. S12), indicating that the fixation process did not disturb the subcellular localization of the constructs.

Additionally, amino acid-based prediction of structural motifs suggests the presence of a putative nuclear localization sequence within the JAKMIP1 C-terminus and a class 1A consensus nuclear export motif for Exportin 1 at the N-terminus (Fig. S13), motifs that are consistent with, and may determine, the observed localization patterns of the N- and C-terminal fragments. The nuclear export sequence appears to exert a stronger influence on JAKMIP1 localization, as JAKMIP1 remains largely cytoplasmic and microtubule-associated throughout the cell cycle (Fig. S14) and fluorescence recovery after photobleaching experiments indicate that JAKMIP1 motility is dependent on the N-terminus (data not presented here).

Considering the nucleoplasmic localization of the JAKMIP1 C-terminus, we examined whether its overexpression could influence *STAT3* expression. We found that overexpression of the C-terminus, but not the full-length protein or N-terminus, slightly increases *STAT3* expression (Fig. 5D). This fits with the observed nucleocytoplasmic localization of each construct and provides further evidence that nuclear populations of JAKMIP1 could regulate gene expression.

### Proteomic profiling of JAKMIP1-containing protein complexes identifies multiple interactions with RBPs involved in transcription and splicing, as well as numerous proteins encoded by ASD risk genes

To further understand the mechanisms by which the JAKMIP1 C-terminal domain could regulate *STAT3* expression, we aimed to identify C-terminus-interacting proteins that could control nuclear gene expression, perhaps via transcription or splicing regulation. To accomplish this, SH-SY5Y cells were transfected with plasmids encoding the various mCherry-tagged JAKMIP1 constructs, and JAKMIP1-containing protein complexes were subsequently immunoprecipitated (Fig. 6A) for identification by liquid chromatography-mass spectrometry.

**Figure 6.**
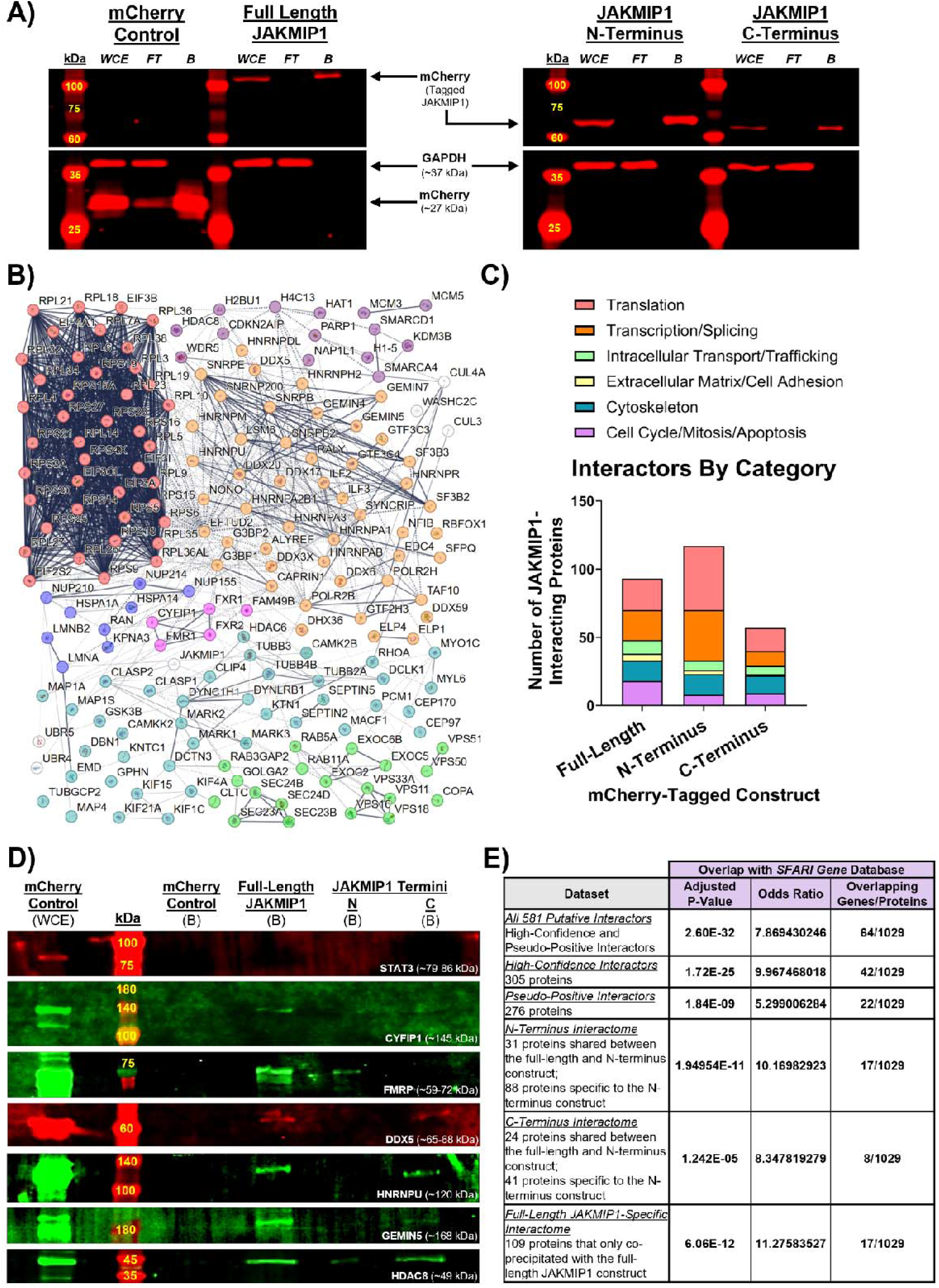
Identifying putative JAKMIP1-interacting proteins from IP-MS data and examining the extent of overlap with protein products of known ASD candidate genes. **A)** Immunoprecipitation efficiency of mCherry-tagged JAKMIP1 constructs confirmed by Western blotting prior to liquid chromatography-mass spectrometry (LC-MS). SH-SY5Y cells were transfected with plasmids encoding the JAKMIP1 mCherry fusion proteins and lysed 48 hours later for immunoprecipitation. WCE-whole cell extract, FT – flow through (or unbound fraction), B – bound fraction. All JAKMIP1 constructs were detected by mCherry at the expected molecular weight and GAPDH (an expected non-JAKMIP1-interacting protein) did not contaminate bound fractions. **B)** Visualization of the network of select JAKMIP1-interacting proteins (not separated by domain specificity), which are grouped into broad functional categories: ribosomal proteins and translation initiation factors (red nodes); regulators of transcription, splicing and mRNA stability (orange nodes); modifiers of histone and chromatin organization as well as DNA replication (purple nodes); proteins involved in nuclear import and export (blue nodes); cytoskeletal and associated proteins (dark cyan nodes); and components of intracellular trafficking pathways (green nodes) (see Table S5 for full list of proteins in each category). Though involved in translational regulation and cytoskeletal remodeling, FMRP (protein product of *FMR1*), CYFIP1 and related proteins are highlighted in a separate central category (pink nodes) due to their direct association with FXS and Dup15q. Generated with *STRING* (Szklarczyk *et al*., 2019). **C)** Number of JAKMIP1-interacting proteins separated by the mCherry-tagged constructs they co-precipitated with. The proteins were categorized by their general function into six categories: proteins involved in or part of “Translation”, “Transcription or Splicing”, “Intracellular Transport or Trafficking”, “Extracellular Matrix or Adhesion”, “Cytoskeleton”, “Cell Cycle or Mitosis or Apoptosis” (see Table S6 for full list of proteins). **D)** JAKMIP1 interactions (or lack thereof) with STAT3, CYFIP1, FMRP, DDX5, HNRNPU, GEMIN5 and HDAC8 were validated by co-immunoprecipitation (co-IP). **E)** Results of over-representation analyses to test for enrichment of the protein products of ASD-associated genes in the JAKMIP1 interactome. The JAKMIP1 interactome is split into separate categories described in the left column. Enrichment analyses were conducted with Fisher’s exact tests, and P-values were adjusted with Benjamini-Hochberg FDR correction to account for multiple testing. The “Overlapping Genes/Proteins” columns refer to the proteins present in a particular category of JAKMIP1-interacting proteins that are encoded by genes listed in *SFARI Gene*. Comparisons with an FDR-adjusted P-value that meets the threshold for statistical significance have been bolded.

This tandem IP-MS approach identified a total of 581 putative JAKMIP1-interacting proteins; with 12 common to all constructs (described as “Sextuple-Positive” interactors), 109 that specifically interact with the full-length JAKMIP1 protein, 119 that interact with the N-terminus, 65 that interact with the C-terminus, and 276 lower-confidence “Pseudo-Positive” interactors (Fig. S15 and Table S4). This putative JAKMIP1 interactome contains proteins that are components of complexes with diverse cellular functions (a select few are presented in Fig. 6B), including ribosomal proteins and translation initiation factors; regulators of transcription, splicing and mRNA stability; modifiers of Histone and chromatin organization as well as DNA replication; proteins involved in nuclear import and export; cytoskeletal and associated proteins; and components of intracellular trafficking pathways. Importantly, FMRP (protein product of *FMR1*) and CYFIP1 both co-precipitate with JAKMIP1, providing further links to FXS and Dup15q.

When comparing the proteins that interact separately with the N- or C-termini, N-terminus is responsible for the majority of JAKMIP1 protein interactions, and no specific domain possesses a sole functional category of interactors (Fig. 6C). This is unexpected as based on localization, the N-terminus should be enriched for cytoskeletal proteins and the C-terminus for nuclear proteins. We suspect that this is likely because of the capability for different JAKMIP1 domains to dimerize with endogenous JAKMIP1 (Fig. S16), which suggests that the domain-specificity of interactions reported in the IP-MS data should be interpreted with caution.

To validate the IP-MS findings, independent co-immunoprecipitation (co-IP) experiments were carried out, testing a range of interactors from expected and novel putative JAKMIP1-interacting proteins (Fig. S17). Importantly, we confirm that JAKMIP1 does not interact with STAT3, so the reduced *STAT3* expression in *JAKMIP1*^−/−^ SH-SY5Y cells is unlikely to be due to a negative feedback loop caused by loss of JAKMIP1 expression. Furthermore, we confirm that JAKMIP1 interacts with FMRP (as expected from (Berg *et al*., 2015)) but add that it also can form complexes with CYFIP1 (Fig. 6D). This reinforces a role for JAKMIP1 in translational regulation and strengthens its link to both FXS and Dup15q. We also find that the JAKMIP1 C-terminus interacts with several RNA-binding proteins (RBPs) – DDX5, HNRNPU and GEMIN5, which the N-terminus does not interact with (Fig. 6D). These RBPs are known regulators of transcription and splicing (Yong *et al*., 2010; Xiao *et al*., 2012; Dardenne *et al*., 2014; Berg *et al*., 2015; Han *et al*., 2022), offering potential ways by which the JAKMIP1 C-terminus can control the synthesis, maturation and stability of new mRNAs. Additionally, we find that JAKMIP1 may interact with epigenetic modifiers of chromatin organization such as HDAC8 (Fig. 6D) (Tang *et al*., 2020). When considered alongside the multiple interactions with Histones (Fig. 6B), these present another mechanism to define how JAKMIP1 may control transcriptional output via chromatin accessibility to transcription factors.

Taking a similar approach with the protein interactors identified by IP-MS, we also tested whether the JAKMIP1 interactome is enriched for the products of ASD risk genes. We find that JAKMIP1 interacts with a large number of proteins encoded by ASD candidate genes (Fig. 6E). Though not definitive proof, these results indicate that JAKMIP1 likely functions in cellular pathways important in neurodevelopment and dysregulation of it or its interactome may be relevant towards the etiology of ASD.

### Dup15q hiPSCs and iNeurons display enhanced IL-6/STAT3 signaling and display an altered neuritogenic response to Hyper IL-6

The data presented thus far suggest that JAKMIP1 may modulate the expression of genes relevant to both inflammatory signaling and ASD. To test whether altered JAKMIP1 expression in ASD could modify how cells respond to cytokines, we used an hiPSC line (SCC115) derived from an individual with an interstitial triplication at chromosome 15, cytoband 15q11.2-q15 (Fig. 7A) to model Dup15q. Along with a control line (CTRM336S), the hiPSCs were differentiated to produce cortical iNeurons (Fig. 7A; Fig. S18). Consistent with the triplication and as expected for a Dup15q model, the SCC115 hiPSCs and iNeurons demonstrated increased *CYFIP1* expression (Fig. S19). However, *JAKMIP1* and *STAT3* expression do not follow the trends we observed in *JAKMIP1*^−/−^ SH-SY5Y cells or that expected from previous work (Berg *et al*., 2015). Whilst *JAKMIP1* and *STAT3* mRNA are downregulated in Dup15q hiPSCs (Fig. 7B), only STAT3 protein is reduced in Dup15q hiPSCs (Fig. 7C). In Dup15q iNeurons, *JAKMIP1* mRNA and protein are both strongly upregulated (Fig. 7, D and E), whilst *STAT3* mRNA is increased (Fig. 7D) but protein reduced (Fig. 7E). Taken together, increased dosage and expression of *CYFIP1* or other 15q11.2-q15 genes in Dup15q may exert distinct effects on JAKMIP1 expression across different cell types and stages of neuralization.

**Figure 7.**
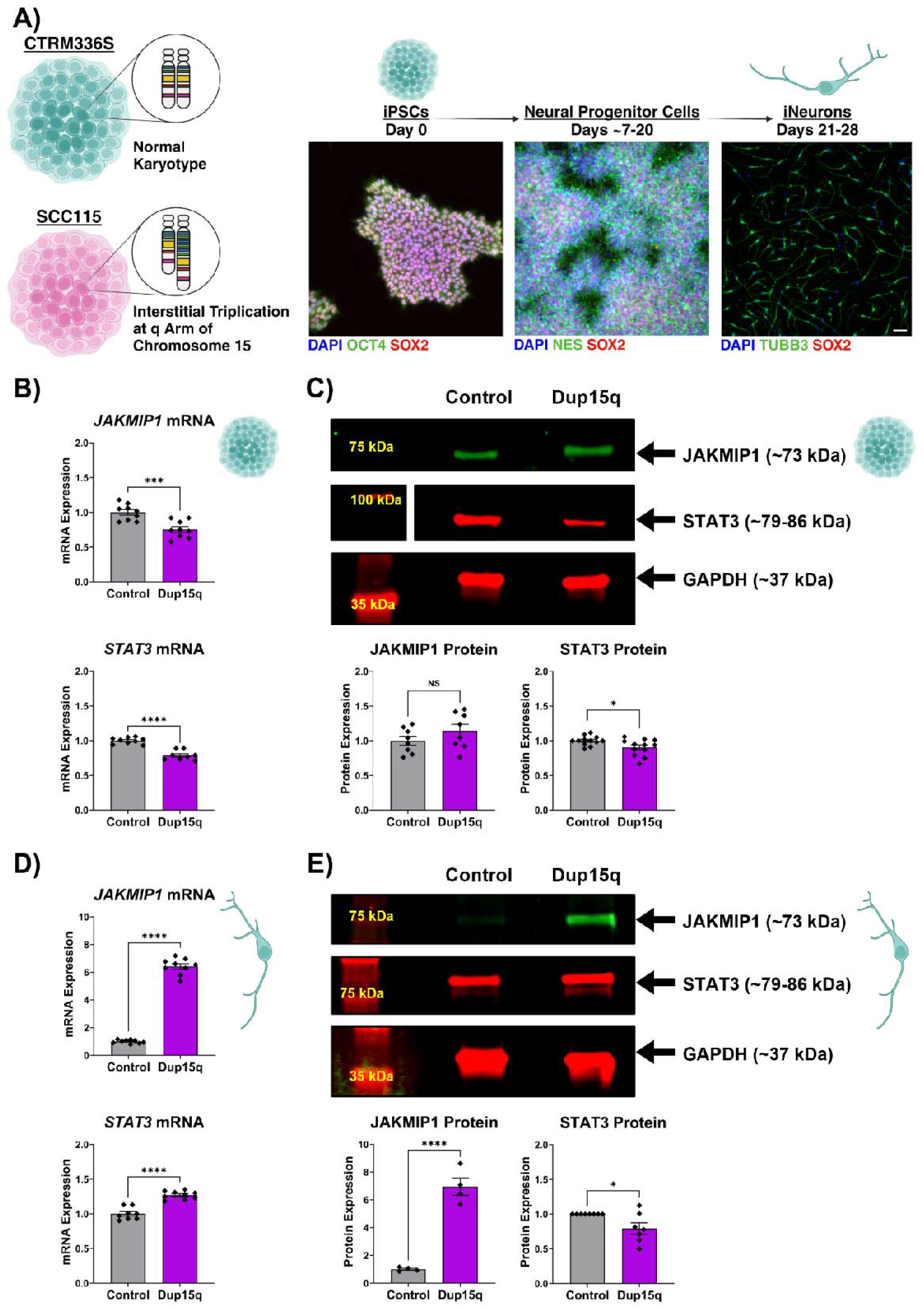
Dup15q hiPSCs and iNeurons display altered JAKMIP1 and STAT3 expression. **A)** Control (CTRM336S) and Dup15q (SCC115) hiPSCs with an interstitial triplication at the q arm of chromosome 15q were used in these experiments. hiPSCs were stained for pluripotency markers Octamer-Binding Protein 4 (OCT4; green) and SRY-Box Transcription Factor 2 (SOX2; red), then differentiated into neural progenitor cells (NPCs). NPCs were stained for Nestin (NES; green) to confirm NPC identity and SOX2 (red) on Day 19 of differentiation. Following terminal plating on Day 21 of differentiation, hiPSC-derived cortical neurons (iNeurons) were stained on Day 28 for the neuronal marker, β(III)-Tubulin (TUBB3; green) and for SOX2 (red) to confirm its lack of expression. Nuclei were counterstained with 4’,6-Diamidino-2-Phenylindole (DAPI; blue). Scale bar = 100 μm. **B)** Relative mRNA expression of JAKMIP1 and STAT3 in control and Dup15q hiPSCs measured by qRT-PCR. Unpaired Student’s T-Tests. **C)** Western blotting was performed to measure JAKMIP1 and STAT3 expression in control and Dup15q hiPSCs. Densitometric analysis of JAKMIP1 and STAT3 relative to the GAPDH loading control. Statistics as in B). Note that the three images are from three separate membranes. **D)** Relative mRNA expression of *JAKMIP1* and *STAT3* in control and Dup15q iNeurons on Day 28 measured by qRT-PCR. Statistics as in B). **E)** Western blotting was performed to measure JAKMIP1 and STAT3 expression in control and Dup15q iNeurons. Densitometric analysis of JAKMIP1 and STAT3 relative to the GAPDH loading control. Statistics as in B). Values presented as mean ± SEM of N = 3-4 independent experiments. NS – not significant, *P < 0.05, ***P < 0.001, ****P < 0.0001. Created with BioRender.com.

Since STAT3 expression is commonly reduced in both the Dup15q hiPSCs and iNeurons, we hypothesized that this may lead to impaired IL-6/STAT3 signaling as observed in *JAKMIP1*^−/−^ SH-SY5Y cells. To test neuronal STAT3 responses, Hyper IL-6 (a chimeric fusion of IL-6 and soluble IL-6Rα (sIL-6Rα)) was used for iNeuron treatments as the iNeuron cells used for this study have not had enough time to mature and express IL-6Rα (Couch *et al*., 2023), thus IL-6 does not elicit a response in these cells (Fig. S20). Instead of being impaired, however, IL-6-induced STAT3 responses appear to be enhanced in both Dup15q hiPSCs and iNeurons despite the reduced baseline STAT3 protein expression (Fig. 8, A-J). This was determined by increased STAT3 (Tyr^705^) phosphorylation following IL-6 (hiPSC) or Hyper IL-6 (iNeuron) treatment (Fig. 8, A-C and F-H) and validated by qRT-PCR for *SOCS3* and *STAT3* (Fig. 8, D-E and I-J). These results could imply that the complex genetic background of Dup15q results in a change to the regulatory networks that control STAT3 activity that supersedes the function of JAKMIP1 in the control CTRM336S cells.

**Figure 8.**
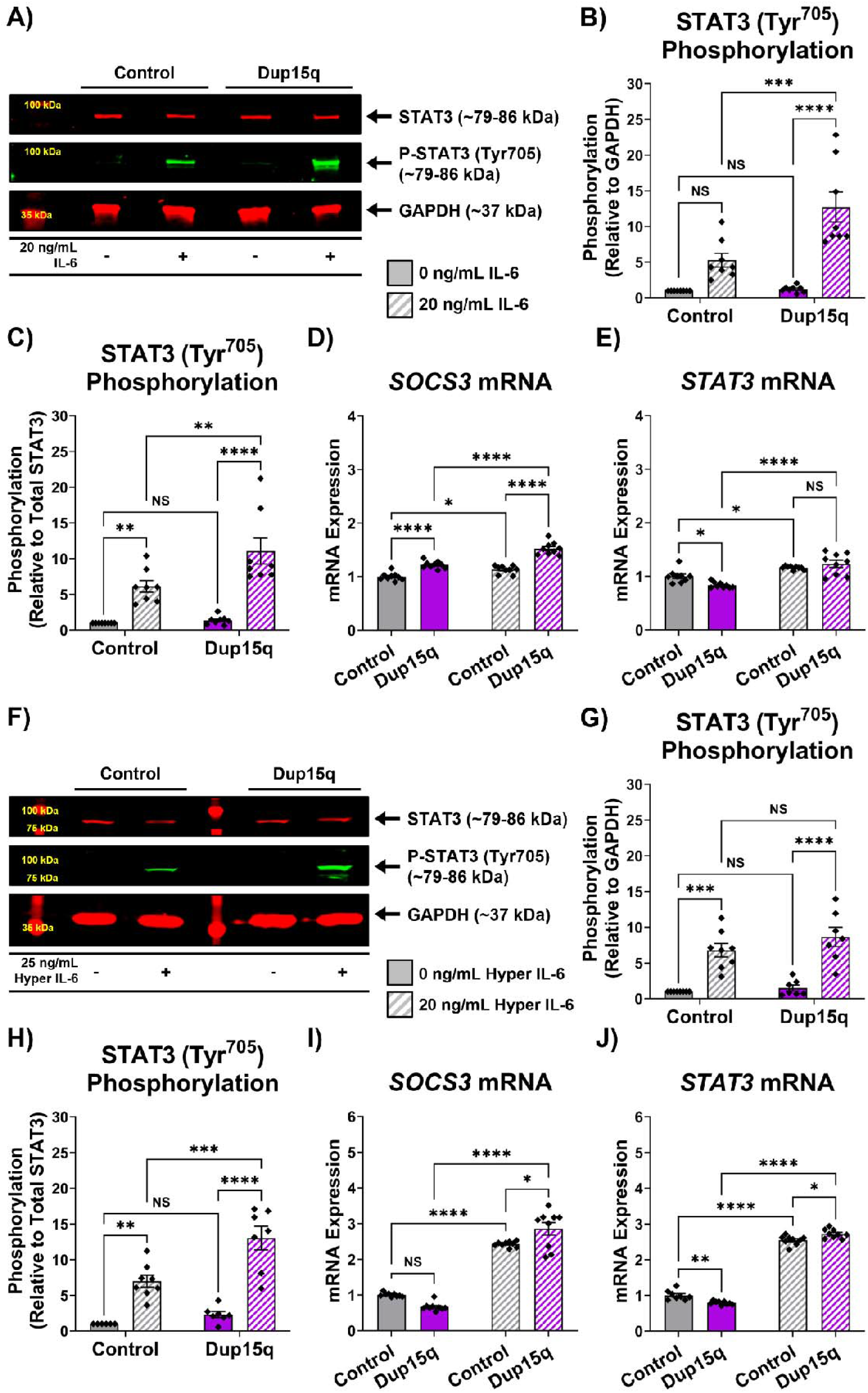
Dup15q hiPSCs and iNeurons display increased IL-6-induced STAT3 (Tyr^705^) phosphorylation and enhanced STAT3 transcriptional activity. **A)** Control (CTRM336S) and Dup15q (SCC115) hiPSCs were treated with StemFlex™ medium with or without 20 ng/mL IL-6 for 30 minutes prior to lysis for protein extraction. Western blotting was performed to measure STAT3 expression and STAT3 (Tyr^705^) phosphorylation. **B)** Densitometric analysis of phosphorylation at the Tyr^705^ residue (P-STAT3^Tyr705^) relative to the GAPDH loading control from A). Two-way ANOVA with Tukey’s HSD test for multiple comparisons. **C)** Densitometric analysis of P-STAT3^Tyr705^ relative to total STAT3 protein from A). Statistics as in B). **D)** Relative mRNA expression of *SOCS3* in control and Dup15q hiPSCs exposed to complete medium with or without 20 ng/mL IL-6 for 4 hours, measured by qRT-PCR. Statistics as in B). **E)** Relative mRNA expression of *STAT3* in control and Dup15q hiPSCs exposed to complete medium with or without 20 ng/mL IL-6 for 4 hours, measured by qRT-PCR. Statistics as in B). **F)** Control and Dup15q iNeurons were treated B-27™-supplemented Neurobasal™ medium with or without 25 ng/mL Hyper IL-6 (sIL-6Rα + IL-6 fusion protein) for 30 minutes on Day 28 prior to lysis for protein extraction. Western blotting was performed to measure STAT3 expression and STAT3 (Tyr^705^) phosphorylation. **G)** Densitometric analysis of P-STAT3^Tyr705^ relative to the GAPDH loading control from F). Statistics as in B). **H)** Densitometric analysis of P-STAT3^Tyr705^ relative to total STAT3 protein from F). Statistics as in B). **I)** Relative mRNA expression of *SOCS3* in control and Dup15q hiPSCs exposed to B-27™ Neurobasal™ medium with or without 25 ng/mL Hyper IL-6 for 4 hours on Day 28, measured by qRT-PCR. Statistics as in B). **J)** Relative mRNA expression of *STAT3* in control and Dup15q hiPSCs exposed to B-27™ Neurobasal™ medium with or without 25 ng/mL Hyper IL-6 for 4 hours on Day 28, measured by qRT-PCR. Statistics as in B). Values presented as mean ± SEM of N = 3 independent experiments. NS – not significant, *P < 0.05, **P < 0.01, ***P < 0.001, ****P < 0.0001.

To test how these signaling changes may impact neuronal development in Dup15q, we examined how exposure to Hyper IL-6 affects Dup15q iNeuron morphology, as IL-6/STAT3 signaling is purported to induce neurite outgrowth. At baseline, we observed that Dup15q iNeurons produce longer neurites (Fig. 9, A-C) and greater numbers of neurites (Fig. 9, E and F), as well as demonstrate increased neurite branching (Fig. 9, G and H). The differences in neurite number and branching are particularly striking – with more Dup15q cells producing ≥4 neurites or with ≥3 neurite branch points (Fig. 9, E and G). However, we find that the Dup15q iNeurons do not show a strong response to Hyper IL-6. In control iNeurons, exposure to Hyper IL-6 treatment following terminal plating raises longest neurite length (Fig. 9C), total neurite length (Fig. 9D), neurite number (Fig. 9F) and branch point number (Fig. 9H), however, only branch point number (Fig. 9H) is affected by Hyper IL-6 in Dup15q cells. Although the lack of change in neurite lengths or number seem to disagree with the enhanced STAT3 activation or activity, these findings do support that neuronal responses to IL-6 are altered in Dup15q.

**Figure 9.**
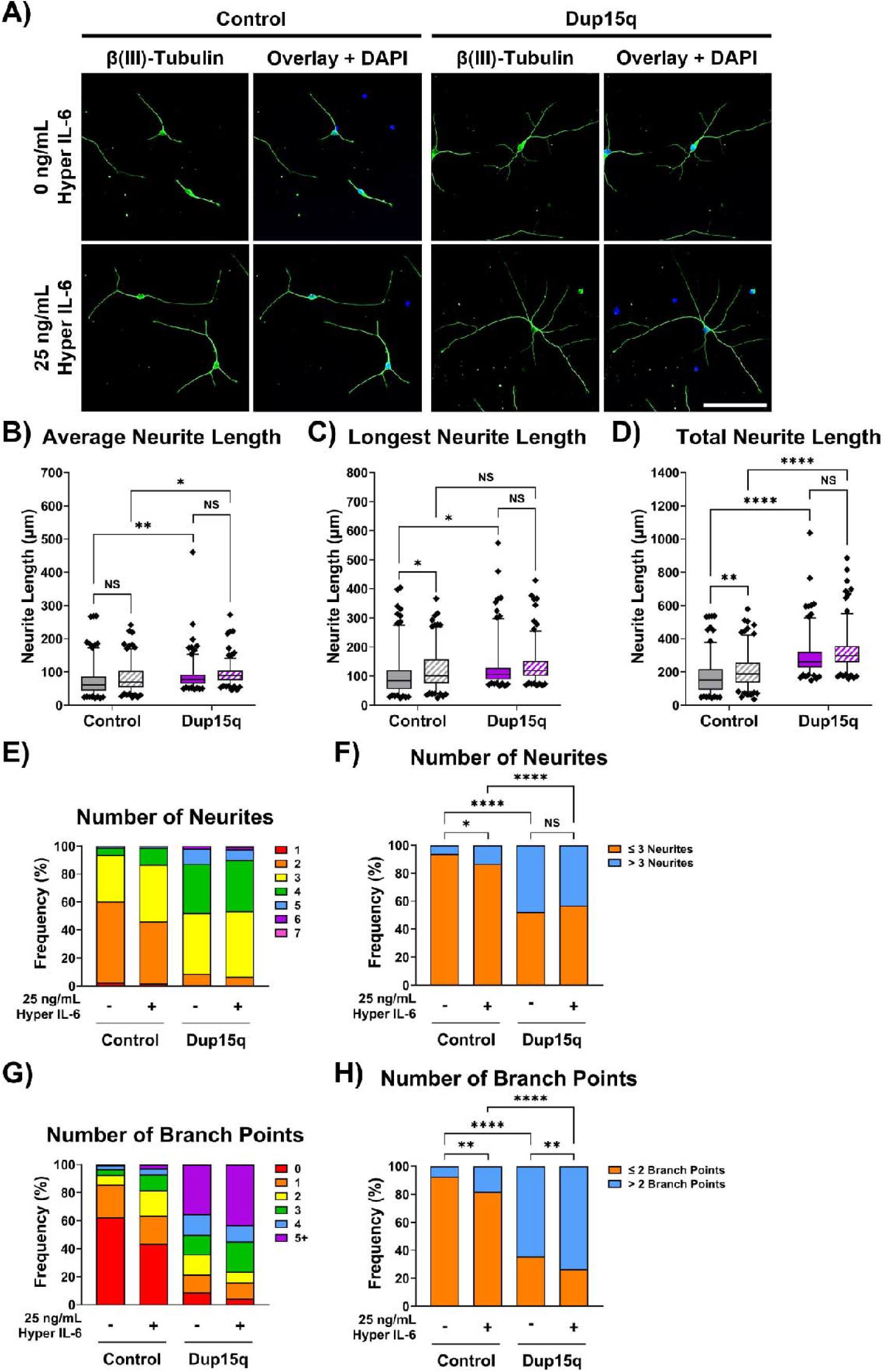
Dup15q iNeurons display increased neurite production, extension and branching but reduced neuritogenic response to Hyper IL-6. **A)** Control and Dup15q iNeurons were terminally plated on Day 21, then treated with or without 25 ng/mL Hyper IL-6 in B-27™ Neurobasal™ medium every 48 hours until fixation on Day 28. iNeurons were stained for β(III)-Tubulin (TUBB3; green). Nuclei were counterstained with 4’,6-Diamidino-2-Phenylindole (DAPI; blue). Scale bar = 100 μm. **B)** Measurement of the average neurite length (ANL) produced by individual neurons in part A). Two-way ANOVA with Tukey’s HSD test for multiple comparisons. **C)** Measurement of the length of the longest neurite (LNL) produced by individual neurons in part A). Statistics as in B). **D)** Measurement of the sum length of all neurites (TNL) produced by individual neurons in part A). Statistics as in B). Boxplots in parts B-D) display the distribution of neurite lengths, with whiskers extending to 5^th^ and 95^th^ percentiles of the data. **E)** Frequency of iNeurons producing 1, 2, 3, 4, 5, 6, or 7 neurites expressed as the percentage of total cells (%). **F)** Frequency of cells producing either ≤3 or >3 neurites following recategorization of data from part E). Comparison of the frequency distributions using Fisher’s exact tests, and P-values were adjusted with Benjamini-Hochberg FDR correction to account for multiple testing. **G)** Frequency of iNeurons with 0, 1, 2, 3, 4, or ≥5 (5+) neurite branch points expressed as the percentage of total cells (%). **H)** Frequency of cells possessing either ≤2 or >2 neurites following recategorization of data from part G). Statistics as in F). N = 3 independent experiments. NS – not significant, *P < 0.05, **P < 0.01, ****P < 0.0001.

## Discussion

### JAKMIP1 as a modulator of neuronal cytokine signaling

Based on its high neuronal expression and capability to interact with JAKs and microtubules (Steindler *et al*., 2004; Vidal *et al*., 2009), we hypothesized that JAKMIP1 may play a role in neuronal cytokine signaling. In this study, we first confirmed that JAKMIP1 is capable of associating with cytokine receptors, particularly the IL-6R complex, in the neuronal SH-SY5Y cell line and in primary chick neurons. However, association with the IL-6R complex does not necessarily mean that JAKMIP1 plays a direct role in regulating neuronal IL-6/JAK1/STAT3 signaling. JAKMIP1 localization seems to largely be regulated by its N-terminus, and it seems primarily associated with microtubules. As previously reported with GABA_B_Rs, it is possible that JAKMIP1 simply acts as an adaptor protein to traffic JAK1/TYK2-associated receptor complexes along microtubules (Vidal *et al*., 2007). Thus, we used a variety of biochemical assays to test whether *JAKMIP1*^−/−^ SH-SY5Y cells display changes in STAT3 activity downstream of IL-6 signaling.

We observed deficits in STAT3 transcriptional activity in *JAKMIP1*^−/−^ SH-SY5Y cells, which appears to stem from reduced *STAT3* mRNA expression and capacity to respond to IL-6. Importantly, the loss of JAKMIP1 expression does not appear to directly change STAT3 phosphorylation kinetics, but rather the transcriptional output of STAT3-responsive genes (such as *SOCS3* and *STAT3*) following IL-6 stimulation. Furthermore, we find that IL-6-induced neuritogenesis is impaired in *JAKMIP1*^−/−^ SH-SY5Y cells, providing a functional output for assessing the impact of altered JAKMIP1-regulated IL-6/STAT3 signaling on neuronal behavior. The neurite tracing experiments suggest that whilst JAKMIP1 is not a crucial mediator of neurite outgrowth (in contrast to a previous study (Vidal *et al*., 2012)), it may play a more central role in translating cytokine signals to the microtubule cytoskeleton. Thus, the results presented in this study point towards a role for JAKMIP1 in transcriptional regulation of *STAT3* expression, which allows JAKMIP1 to modify transcriptional and morphogenic responses to IL-6 in neuronal cells. However, it is important to note that the *JAKMIP1*^−/−^ SH-SY5Y cells still successfully form neurites and that the observed neurite outgrowth deficits are not nearly as drastic as those reported by Vidal et al. (Vidal *et al*., 2012), so it is important to consider that the SH-SY5Y cells used in this study may not behave as true neurons.

Transcriptomic profiling of the *JAKMIP1*^−/−^ SH-SY5Y cells identified additional cytokine-related pathways that could potentially be affected by JAKMIP1 deficiency, such as TGF-β/SMAD and NF-κB signaling. However, these signaling pathways tend to be related to one another, with a large degree of crosstalk, for example, existing between STAT3 and NF-κB signaling (Cauvi *et al*., 2017). Therefore, it is possible that the DEGs observed when comparing these pathways stem from long-term alterations to STAT3 expression and activity. This highlights a potential limitation of this approach wherein the RNA-seq data provide a detailed ‘snapshot’ of those cellular pathways which could be altered following long-term JAKMIP1 deficiency but does not necessarily inform the direct causes and effects on gene expression. Nevertheless, it is clear that the effects of JAKMIP1 on cytokine signaling are not limited to STAT3 signaling but likely extend to multiple pathways. We postulate that JAKMIP1 may also influence the signaling of other STAT3-agonistic cytokines beyond IL-6, though this will require further investigation to confirm.

To test our findings in a true neuronal model, we obtained Dup15q hiPSCs and generated iNeurons. Crucially, though the SH-SY5Y experiments support a role for JAKMIP1 in regulating IL-6/STAT3 signaling, similar experiments performed in Dup15q hiPSCs and iNeurons suggest this may not always be the case. Changes in *JAKMIP1* and *STAT3* mRNA expression occur variably throughout the neuralization process, and we also find that STAT3 activation and transcriptional activity are enhanced in both Dup15q hiPSCs and iNeurons. This enhancement does not seem to be caused by changes in *JAKMIP1* expression, which could mean that whilst JAKMIP1 may regulate STAT3 activity in ‘normal’ cells, it likely plays a different role in Dup15q. We hypothesize that in Dup15q, one of the genes encoded in the chromosome 15q11.2-q15 region exerts stronger control over STAT3 activation/function, ultimately causing a state where STAT3 is more responsive to IL-6 regardless of changes in JAKMIP1 expression. Although it is unclear which gene this is, we hypothesize that *CYFIP1* could be responsible for this, as we have separately demonstrated that *CYFIP1* overexpression enhances IL-6/STAT3 signaling even though it reduces *STAT3* expression (Martin *et al*., 2026), findings which are highly reminiscent of the data presented here from Dup15q hiPSCs and iNeurons.

However, unlike with *JAKMIP1*^−/−^ SH-SY5Y cells, we were unable to confirm whether this enhancement of STAT3 signaling in the Dup15q iNeurons augments neurite outgrowth. Although Hyper IL-6 exposure did induce an increase in neurite lengths, neuritogenesis and neurite branching in control iNeurons, the increase was attenuated in the Dup15q iNeurons. We suspect that this is likely because Dup15q iNeurons produce significantly greater numbers of neurites and branch points under basal conditions so that the addition of Hyper IL-6 fails to induce a major increase in neurite outgrowth. We could also interpret this drastic difference in neuritogenesis to indicate that at rest conditions, Dup15q iNeurons exist in a state that is different from the control iNeurons in such a way that exposure to Hyper IL-6 results in a limited morphogenic response compared to the control iNeurons. Without further research, it is difficult to pinpoint which gene within the Dup15q region is altering neuritogenesis in the iNeurons, most strikingly causing increased branching. Given that JAKMIP1 expression was also largely increased in Dup15q iNeurons, and that we observed only a subtle impairment in neurite extension in the JAKMIP1-/- SH-SY5Y cells, we can postulate that perhaps JAKMIP1 exerts a stronger effect on neurite branching as opposed to neurite elongation as initially expected. To determine whether these observed effects are due to changes in JAKMIP1 expression JAKMIP1-knockdown approaches should be explored.

### Potential mechanisms for JAKMIP1 control of *STAT3* expression

The mechanisms by which JAKMIP1 modulates *STAT3* expression remain unclear. However, our findings suggest that this is likely to be mediated via the C-terminus of the protein, which can interact with RBPs such as DDX5 and HNRNPU. Interestingly, DDX5 has previously been shown to modulate *STAT3* expression and activity (Sarkar, Khare and Ghosh, 2016), and though not directly linked to STAT3, HNRNPU is known to regulate various aspects of RNA biology (Sapir and Reiner, 2023) so it would not be surprising if HNRNPU were to be discovered as a modulator of STAT3 expression. Most importantly, HNRNPU is a known interactor of DDX5 (Han *et al*., 2022), which further cements that JAKMIP1 could form a component of an RBP complex, perhaps one comprised of DDX5 and HNRNPU.

As overexpression of the JAKMIP1 C-terminus only results in a modest upregulation of *STAT3* mRNA, we suspect that JAKMIP1 is unlikely to act as a transcription factor. Instead, JAKMIP1 could function as an adaptor protein that supports transcription. Costa *et al*. previously predicted a highly coiled coil structure for the JAKMIP1 protein, and that the C-terminus domain may possess a leucine zipper motif (Costa *et al*., 2007). More recently developed tools for protein structure prediction, such as *AlphaFold*, also show a highly helical structure for JAKMIP1 and predict coiled coil structures (Fig. S21). Though not proof of DNA-binding capability, helical structures and leucine zipper domains of JAKMIP1 are characteristic features of DNA-binding proteins of the Basic Leucine Zipper (bZIP) superfamily of transcription factors (Seldeen *et al*., 2010). In addition, JAKMIP1 contains several basic amino acid residues which are commonly found in DNA-binding proteins, such as bZIPs and Histones, that interact with the negatively charged DNA phosphate groups (Seldeen *et al*., 2010; Lin and Guo, 2019). These motifs may facilitate dimerization of JAKMIP1 and potential interactions with DNA-binding proteins, which fit with the observed nucleoplasmic localization.

It is also important to acknowledge that JAKMIP1 could control *STAT3* expression in more indirect ways, such as through regulation of splicing or chromatin accessibility. We find that components of the Survival of Motor Neurons complex, such as GEMIN5, co-precipitate with the JAKMIP1 C-terminus. GEMINs are integral to spliceosome assembly (Battle *et al*., 2006) and could represent another mechanism for control of *STAT3* expression, as it is known that splicing can ‘talk’ back to and modulate transcription (Tellier, Maudlin and Murphy, 2020). Additionally, JAKMIP1 seems to be able to interact with HDAC8, along with several Histones, which may allow JAKMIP1 to influence STAT3 expression via changing chromatin accessibility (Hu *et al*., 2000; Tang *et al*., 2020). Even the JAK proteins, which interact with the JAKMIP1 C-terminus (Steindler *et al*., 2004), are reported to modify Histone post-translation modifications as well (Zouein, Duhé and Booz, 2011). Considering the prior association of JAKMIP1 with euchromatin (Vidal *et al*., 2009), a role for JAKMIP1 in epigenetic regulation of gene expression is possible.

Nevertheless, our current evidence suggests that a nuclear mechanism for JAKMIP1 regulation of *STAT3* expression is most likely; and that this is mediated by the JAKMIP1 C-terminus, which primarily localizes to the nucleoplasm. Future work should use chromatin immunoprecipitation sequencing (ChIP-seq) or Assay for Transposase-Accessible Chromatin using sequencing (ATAC-seq) in JAKMIP1-deficient cells or cells overexpressing the JAKMIP1 C-terminus to investigate potential mechanisms by which the JAKMIP1 C-terminus modulates *STAT3* expression.

### JAKMIP1-regulated genes, -interacting proteins and their relevance to Dup15q and wider ASD etiology

Outside the context of cytokine signaling, we find that JAKMIP1 deficiency disproportionately affects the expression of multiple ASD risk genes (as curated by *SFARI Gene*), and DEGs identified in *JAKMIP1*^−/−^ SH-SY5Y cells show significant overlap with those differentially expressed in other ASD models. This suggests that JAKMIP1 may regulate the expression of genes relevant to the etiology of ASD. Examples of ASD-relevant DEGs observed in *JAKMIP1*^−/−^ SH-SY5Y cells include classical axon guidance molecules and their receptors (e.g., *ROBO2*, *SLIT2*, *SEMAs*, *UNCs*, etc.), synapse scaffolding proteins (e.g., *DLG2*, *DLGAP2*, *SHANKs*, etc.) and synaptic cell adhesion molecules (e.g., *CNTNs*, *CDHs*, *CNTNAP5*, *NRXNs*, *NLGN1*, etc.). Disruptions to these proteins and the genes that encode them are often associated with ASD and other neuropsychiatric disorders (Bourgeron, 2015; Van Battum, Brignani and Pasterkamp, 2015; de la Torre-Ubieta *et al*., 2016; Gandawijaya *et al*., 2021).

In addition to the genes themselves, we find the JAKMIP1 interactome is enriched for the products of multiple ASD-associated genes. As the IP-MS experiments indicate that the JAKMIP1 C-terminus can interact with a range of RBPs, we suspect that some of these JAKMIP1-interacting RBPs may have a direct role in controlling the expression of genes involved in ASD-relevant pathways, though this will require further work to confirm. Importantly, both FMRP and CYFIP1 also co-precipitate with JAKMIP1, reinforcing the concept that JAKMIP1 may form a downstream molecular link between FXS and Dup15. Furthermore, there exist overlapping gene expression signatures between *JAKMIP1*^−/−^, *Fmr1*^−/−^ and *CYFIP1*-knockdown models, which imply that JAKMIP1 may control the expression of a subset of genes sensitive to *FMR1* and *CYFIP1* disruption. We also discover that genes likely induced by maternal immune activation overlap with components of the JAKMIP1 interactome, which indicates that JAKMIP1 may be involved in the neural immune/inflammatory response and provides further evidence to support JAKMIP1-regulated genes as likely relevant to ASD development. Yet, considering that JAKMIP1 expression is highly raised in the Dup15q iNeurons, it is difficult to relate the RNA-seq findings presented here directly to Dup15q. According to Nishimura et al. (Nishimura *et al*., 2007), *JAKMIP1* expression appeared to be raised in lymphoblastoid cell lines from cases of FXS and Dup15q but was instead reduced in neural models of altered *FMR1* and *CYFIP1* expression. Thus, additional work is required to confirm whether *JAKMIP1* expression is truly raised in ASD (particularly in the nervous system), and if so, to discover the impact of *JAKMIP1* expression changes.

In maternal immune activation models, upregulated IL-6 signaling plays a central part in the development of sociability deficits and restrictive behaviors (Wu *et al*., 2017; Cieślik *et al*., 2020; Zawadzka, Cieślik and Adamczyk, 2021). Whilst the GSOA suggests that JAKMIP1 may regulate genes and pathways that are impacted by neuroinflammation, our results demonstrate that JAKMIP1 deficiency restrains IL-6 signaling. If reduced *JAKMIP1* expression is associated with ASD, then it should in theory be protective against sustained IL-6 signaling, which does not fit with the maternal immune activation or post-mortem tissue studies that implicate IL-6 in ASD development. Coincidentally, the enhanced IL-6/STAT3 signaling observed in the Dup15q hiPSCs and iNeurons fits very well with these other studies, highlighting increased levels of IL-6 or enhanced responsiveness to IL-6 as a convergent pathway that may link different risk factors for ASD (CNVs such as in Dup15q and neuroinflammation). As of now, further research is required to determine how (or if) JAKMIP1 may be related to the increased/enhanced IL-6/STAT3 signaling in ASD.

### Limitations and future directions

There are several limitations to consider when interpreting the findings of this study, particularly regarding the cell lines used. A large proportion of the mechanistic work was conducted in SH-SY5Y cells and although SH-SY5Y cells are a tractable neural model for investigating cytokine signaling pathways, they do not fully recapitulate the properties of neurons. Furthermore, as a neuroblastoma cell line, it is possible that SH-SY5Y cells may display altered baseline cytokine signalling responses. Therefore, to validate our findings and to explore the effects of altered JAKMIP1 expression on STAT3 expression and activity in an ASD-relevant model, iNeurons from a Dup15q model were used. However, for all hiPSC-based experiments, a single control and Dup15q line were used; whilst these experiments provide insight into how Dup15q altered neuronal cytokine signaling pathways, it does not account for individual genetic differences. Expanding these experiments to include multiple independent control and Dup15q lines, and ideally alongside the use of isogenic controls, may help to better define the contribution of the various genes at the Dup15q locus to neuroinflammatory signaling in ASD.

Importantly, although the data presented here are consistent with a role for JAKMIP1 in modulating IL-6/STAT3 signaling in SH-SY5Y cells, a direct causal relationship has not been established in human iNeurons. The altered cytokine responses observed in Dup15q cells are interpreted in relation to dysregulated JAKMIP1 expression, however, considering our findings and the complex genetic landscape of Dup15q, it remains unclear to what extent IL-6/STAT3 signaling is modulates by JAKMIP1 in Dup15q. Further studies employing targeted manipulation of JAKMIP1 expression in Dup15q iNeurons may help to clarify its contribution and determine whether it directly regulates neuronal responses to cytokines in Dup15q and more broadly, syndromic ASDs.

## Materials and Methods

Cell culture, CRISPR gene-editing, RNA and protein isolation, qRT-PCR, RNA-seq, Western blotting, and immunocytochemical staining were performed as previously described (Harlow *et al*., 2022; Martin, Gandawijaya and Oguro-Ando, 2022). Sections below provide a brief overview of general methods; see File S3 for additional details.

### General cell culture and maintenance

Human neuroblastoma SH-SY5Y and embryonic kidney HEK-293 cells (#CRL-2266™ and #CRL-1573™, *American Type Culture Collection*; HEK-293 cells were a gift from Dr. John K. Chilton) were maintained in a 1:1 mixture of Dulbecco’s modified eagle medium and Ham’s nutrient mixture F-12 supplemented with GlutaMAX™ (#31331093, Gibco™; this is referred to simply as DMEM/F-12 in later sections) containing 10% fetal bovine serum (FBS; #10500064, *Gibco*™) (FBS-supplemented DMEM/F-12 is henceforth referred to as DMEM/F-12 + 10% FBS) at 37°C, 5% CO2, 95% humidity. Upon reaching 70-80% confluency, SH-SY5Y or HEK-293 cells were dissociated at using TrypLE™ (#12604013, *Gibco*™), which was inactivated with an equal volume of DMEM/F-12 + 10% FBS, and the cells, and cells passaged or counted and seeded into new culture vessels for experiments. Cell passage numbers were kept as close to each other among experimental replicates. At least three independently passaged wells/dishes of cells were used for all experiments.

### hiPSC cell culture, maintenance and neuralization

In this work, we used two human induced pluripotent cell lines (hiPSCs): a control cell line, CTRM336S, gifted by Prof. Deepak Srivastava; and a Dup(15q) cell line, SCC115 (carries an interstitial triplication of the chromosome 15q11.2-q15 region), obtained from and authenticated by the *University of Connecticut Stem Cell Core*. Validation of pluripotency marker expression was performed on receipt and checked again prior to neuralization.

hiPSCs were maintained in StemFlex™ media (#A3349401, *Gibco*™) at 37°C, 5% CO2, 95% humidity in 6-well plates coated with 1% Geltrex™ (#A1413301, *Gibco*™). Cells were passaged at 70% confluency by incubation with Versene™ solution (#15040066, *Gibco*™) and detached by washing with StemFlex™ media and using a cell lifter to retain intact colonies. For all hiPSC experiments, cell passage numbers were kept as close to each other among experimental replicates, between passage 20-30. A minimum of three independently passaged wells of cells were used for all experiments.

Upon reaching 100% confluency, hiPSCs were differentiated into neural progenitor cells (NPCs) and cortical neurons (iNeurons) as previously reported (Warre-Cornish *et al*., 2020) following a dual SMAD inhibition protocol and Wnt signaling inhibition protocol. At the start of neuralization (Day 0), StemFlex™ was replaced with a N-2:B-27 media, a 1:1 mixture of N-2 (#17502048, *Gibco*™) and B-27™ (#17504044, *Gibco*™) supplemented media (individual medias prepared according to manufacturer instructions). From Days 1-6, N-2:B-27 was supplemented with was supplemented with the SMAD inhibitors SB431542 (#SM33-10, *Cambridge Bioscience*) and Dorsomorphin (#P5499-5MG, *Sigma-Aldrich*®), and the Wnt inhibitor XAV939 (#X3004-5MG, *Sigma-Aldrich*®). On Day 7, cells were passaged in a 1:1 ratio using Accutase™ (#A1110501, *Gibco*™) and plated in SB431542-, Dorsomorphin- and XAV939-supplemented N-2:B-27 media containing Y-27632 (#Y0503-1MG, *Sigma-Aldrich*®). SB431542, Dorsomorphin and XAV939 are removed from the next day onwards and the cells cultured in N-2:B-27. On Days 12, 16 and 19, NPCs are dissociated with Accutase™ and replated. On Day 21, NPCs are dissociated again and terminally plated onto poly-D-Lysine and Laminin-coated plates and cultured in B-27™ media supplemented with DAPT (#D5942-5MG, *Sigma-Aldrich*®). DAPT was refreshed daily until Day 28, after which the iNeurons were ready for experiments. A minimum of three independently differentiated wells of iNeurons were used for all experiments.

### Plasmid DNA replication, preparation and transfection

DH5α *Escherichia coli* (gift from Dr. John K. Chilton) were cultured in Luria-Bertani (LB) broth (#L3022-1KG, *Sigma-Aldrich*®) or on LB agar (#L2897-1KG, *Sigma-Aldrich*®) plates for molecular cloning and plasmid DNA replication. To replicate plasmids, chemically competent DH5α E. coli were transformed by heat shock and plated onto antibiotic containing LB agar plates to select for successfully transformed colonies. Following overnight incubation at 37°C, individual colonies were inoculated into larger volumes of antibiotic containing LB broth to replicate the plasmid of interest. The QIAprep® Spin Miniprep (#27106, *Qiagen*) and ZymoPURE II Plasmid Midiprep (#D4201, *Zymo Research*) kits were then used to harvest plasmids from broth cultures, following manufacturer instructions. The quality and concentration of eluted plasmid was determined using a NanoDrop™ ND-8000 spectrophotometer. Plasmid DNA was then transfected into cells with a variety of methods, either with reagents such as Lipofectamine™ LTX (#15338100, *Invitrogen*™) and jetOPTIMUS® (#117-07, *Polyplus-transfection*®) or by Nucleofection™ (#V4XC-2012, *Lonza Bioscience*).

### RNA extraction, cDNA synthesis, PCR and qRT-PCR

TRI Reagent™ (#R2050-1-50, *Cambridge Bioscience*) was used to lyse cells prior to RNA isolation by either the Direct-zol™ RNA MiniPrep kit (#R2052, *Zymo Research*), following manufacturer instructions, or by precipitation following chloroform phase separation. The quality and concentration of eluted/reconstituted RNA was determined using a NanoDrop™ ND-8000 spectrophotometer. RNA was converted to cDNA with the PrimeScript™ RT Reagent kit (#RR038A, *Takara Bio*), following manufacturer instructions, with RNA input into the cDNA synthesis reaction kept consistent among replicates of a given experiment. 100% RNA-to-cDNA conversion efficiency was assumed to estimate cDNA concentration.

Sequences of primers for qRT-PCR can be found in Table S7. Primers were designed using a combination of *Benchling* (Benchling [Biology Software]. (2022). Retrieved from https://benchling.com), *Primer3Plus* (Untergasser *et al*., 2012) and the *UCSC In-Silico PCR* (Kent *et al*., 2002) tools. The coding sequence of a target gene was imported into *Benchling* from *Ensembl* (Cunningham *et al*., 2022). Non-quantitative end-point PCR was performed with the HOT FIREPol® DNA Polymerase (#01-02-00500, *Solis BioDyne*), whereas quantitative reverse transcription PCR (RT-PCR) was performed with the HOT FIREPol® EvaGreen® qPCR Mix Plus (ROX) (#08-24-00001-10, *Solis Biodyne*) in a QuantStudio 12K Flex Real-Time PCR machine (*Thermo Fisher Scientific*).

qRT-PCR data was analyzed by the Pfaffl method (Pfaffl, 2001). The expression of each target gene was normalized to two housekeeping genes, *GAPDH* and *POLR2A*. The resulting gene expression ratios were normalized to control samples of each experiment to express a fold change in mRNA expression value.

### Protein extraction and Western blotting

Adherent SH-SY5Y cells, HEK-293 cells, hiPSCs or iNeurons were washed with ice-cold phosphate-buffered saline (PBS) then lysed on ice with radioimmunoprecipitation assay (RIPA) buffer supplemented with PMSF, Protease Inhibitor Cocktail (#P8340-1ML, *Sigma-Aldrich*®), Phosphatase Inhibitor Cocktail 2 (#P5726-1ML, *Sigma-Aldrich*®) and Phosphatase Inhibitor Cocktail 3 (#P0044-1ML, *Sigma-Aldrich*®). Cell lysates were cleared by centrifugation and protein concentration estimated with the Pierce™ bicinchoninic acid (BCA) protein assay kit (#23227, *Thermo Scientific*™). Absorbance was measured with a PHERAstar FS microplate reader (*BMG Labtech*).

50 μg of total protein was diluted in Laemmli buffer and heated at 70°C for 10 minutes (for phosphoprotein analysis) or 95°C for 5 minutes, then subjected to sodium dodecyl sulfate (SDS) polyacrylamide gel electrophoresis (SDS-PAGE). Following SDS-PAGE, proteins were transferred onto polyvinylidene difluoride (PVDF) membranes by a wet transfer method. PVDF membranes were then sequentially incubated in blocking buffer and antibody solutions, with thrice 0.1% Tween® 20 supplemented tris-buffered saline (TBS-T) washes in between. GAPDH was used as a loading control to normalize the expression of target proteins across different samples. Immunolabelled proteins were then visualized using the Odyssey® CLx Imager (*LI-COR Biosciences*) with near-infrared DyLight™ dye-conjugated secondary antibodies. The Image Studio software (Version 5.2; *LI-COR Biosciences*) was used to perform densitometric analyses comparing the abundance of target proteins between samples. Antibodies used for Western blotting can be found in Table S8.

### Immunocytochemical staining and fluorescence microscopy

SH-SY5Y, HEK-293, hiPSCs or NPCs were seeded onto 13-mm-diameter glass coverslips pre-coated with extracellular matrix proteins. Following overnight incubation (or a specified treatment or differentiation), cells were fixed with 4% paraformaldehyde (PFA; #P6148-1KG, *Sigma-Aldrich*®) for 30 minutes at room temperature. Fixed cells were permeabilized and then sequentially incubated in blocking buffer and antibody solutions, with three PBS washes in between each step. Immunolabelled proteins were visualized with fluorophore-conjugated secondary antibodies. Antibodies used for immunocytochemical staining can be found in Table S8. Nuclei were counterstained with 200 ng/mL DAPI for 15 minutes, then coverslips mounted onto glass microscope slides with ProLong™ Diamond Antifade Mountant (#P36970, *Invitrogen*™). Microscope slides were imaged either on the upright DM4B LED (*Leica Microsystems*) or inverted EVOS FLoid™ (*Invitrogen*™), DMi8 widefield (*Leica Microsystems*) and TCS SP8 confocal microscopes (*Leica Microsystems*).

### Generation of *JAKMIP1*^−/−^ SH-SY5Y lines with CRISPR-Cas9 technology

A CRISPR guide RNA (gRNA) sequence was designed to exon 5 of transcript ENST00000282924.9 of the *JAKMIP1* gene using the CRISPR tool available in *Benchling*. The gRNA and its complementary sequence were purchased from *Integrated DNA Technologies* with sticky ends to facilitate type IIS restriction enzyme cloning (full details of the cloning process are described in File S3). The gRNA sequence was cloned into the pU6-(BbsI)_CBh-Cas9-T2A-mCherry plasmid (a gift from Dr. Ralf Kuehn; *AddGene* plasmid #64324; http://n2t.net/addgene:64324; RRID: Addgene_64324) and the resultant plasmid transfected into SH-SY5Y cells.

Following clonal isolation, genomic DNA was extracted using the PureLink™ Genomic DNA Mini Kit (#K182002, *Invitrogen*™), following manufacturer instructions, and the “*JAKMIP1* (Exon 4-6)” primer was used to amplify the genomic sequence of the JAKMIP1 gene containing the gRNA by PCR. Potential *JAKMIP1*^−/−^ SH-SY5Y lines were then genotyped by Sanger sequencing (through *GENEWIZ*, *Azenta Life Sciences*). Two separate *JAKMIP1*^−/−^ lines were identified, named “KO-A” and “KO-B”, and these were carried forward for further validation of successful gene-knockout by qRT-PCR and Western blotting for *JAKMIP1* expression.

### Assessment of STAT3 activity following IL-6 treatment

Multiple techniques were used to assess STAT3 activation and transcriptional activity downstream of IL-6 stimulation. SH-SY5Y cells, HEK-293 cells and hiPSCs were treated with 20 ng/mL of IL-6 (#7270-IL-025, *R&D Systems*) in their corresponding media. This concentration has previously been shown to be sufficient to induce STAT3 activation (Russell *et al*., 2013), however. iNeurons were treated with 25 ng/mL of Hyper IL-6 (human IL-6/secreted IL-6Rα chimeric protein) (#8954-SR-025, *R&D Systems*) as immature neurons have previously been shown to express very low levels of the IL-6Rα subunit (only detectable in one study around Day 50 of differentiation (Couch *et al*., 2023)). We also confirmed that our iNeurons did not show increased STAT3 (Tyr^705^) phosphorylation or increased *SOCS3* mRNA expression following stimulation with 20 ng/mL IL-6 (Fig. S20).

To capture short-term, rapid changes in STAT3 (Tyr^705^) phosphorylation, cells were exposed to IL-6/Hyper IL-6 for 30 minutes prior to protein extraction and Western blotting.

To measure downstream expression of STAT3-responsive genes, SH-SY5Y cells were treated with IL-6/Hyper IL-6 for four hours prior to cell lysis with TRI Reagent™. For dual-luciferase® reporter (DLR™; #E1960, *Promega*) assays, cells were treated with IL-6 for 18 hours post-transfection with Cignal Reporter plasmids (#336841, GeneGlobe ID: CCS-9028L, *Qiagen*), then lysed to extract the Firefly and Renilla luciferases. Detailed methodology for the DLR™ assays can be found in File S3.

To assess the effects of IL-6 on neurite outgrowth in SH-SY5Y cells, IL-6 was added to the differentiation medium as described in the next section (“SH-SY5Y differentiation to measure neurite outgrowth”). To test the effects of IL-6 on neurite outgrowth and branching in iNeurons, 25 ng/mL of Hyper IL-6 was added to B-27™ medium on Day 21 during terminal plating, and medium was replaced every day until Day 28.

### SH-SY5Y differentiation for neurite outgrowth assessment and RNA-seq

SH-SY5Y cells were differentiated with Brain-Derived Neurotrophic Factor (BDNF) and retinoic acid (RA) under serum starved conditions as previously described (Martin, Gandawijaya and Oguro-Ando, 2022) to induce neurite outgrowth. After 48 hours, differentiating SH-SY5Y cells were fixed with 4% PFA for morphological assessment.

For the RNA sequencing (RNA-seq) experiments, SH-SY5Y cells were differentiated for a longer period (nine days in total) with a higher concentration of BDNF to obtain a more mature, neuron-like population of SH-SY5Y cells (see File S3 for details).

### Neurite tracing for SH-SY5Y cells and iNeurons

Following fixation with 4% PFA, cells were stained for β(III)-Tubulin to visualize gross cell morphology and neurites through the underlying microtubule cytoskeleton. For SH-SY5Y cells, the length of the longest neurite produced by a single cell (i.e., longest neurite length (LNL); a measure of neurite extension) and the sum length of all neurites produced by a single cell (i.e., total neurite length (TNL); a measure of neuritogenesis) were manually measured using the *FIJI* software (ImageJ Version 2.9.0; (Schindelin *et al*., 2012)). For iNeurons, the LNL, TNL and average neurite length (ANL) were manually measured using the FIJI software. Additionally, the number of neurites produced per neuron and the number of branch points per neuron were manually counted. Prior to neurite length analysis, a scale bar was used to set a pixel to mm ratio for each image, after which the “Segmented Line” tool was used to measure the distance (in μm) from the border of the nucleus to this distal end of each neurite.

### Transcriptomic profiling by RNA-seq

9-day differentiated SH-SY5Y cells were lysed with TRI Reagent™ and RNA purified with the Direct-zol™ RNA MiniPrep kit. RNA quality, purity and concentration were determined with the Qubit® 2.0 Fluorometer (*Invitrogen*™; with #Q32866) and the Agilent 2200 TapeStation (*Agilent Technologies*; with #5067-5576, #5067-5577 and #5067-5578) A total of four RNA samples per differentiated SH-SY5Y cell line (12 samples total across the EV control, KO-A and KO-B cell lines) with high concentration and RNA integrity number ≥ 8 were selected to be taken forward for RNA-seq. 1 μg of total RNA was prepared for each sample and submitted to the Exeter Sequencing Service for library preparation with the TruSeq DNA Library Prep Kit HT with oligo(dT) primers against the 3’-end poly(A)-tail (#FC-121-2003, *Illumina*) and next generation sequencing on the HiSeq 2500 platform (*Illumina*), producing 50-bp paired-end sequencing data.

All bioinformatic analyses were performed on the University of Exeter’s *ISCA* high-performance computing (HPC) environment or in *RStudio* (Version 4.0.0; (RStudio Team, 2020)) with Version 4.0.0 of *R* (R Core Team, 2020). Scripts written to perform these analyses and an overview of the analysis workflow can be found in File S3. Adapter sequences were pre-trimmed by the *Exeter Sequencing Service* with *Cutadapt* and quality of sequencing reads was checked with *FastQC* (Version 0.11.7; (Andrews, 2010)) and consolidated with *MultiQC* (Version 1.2; (Ewels *et al*., 2016)). *FastQC* showed that sequencing adapters were successfully removed; that base calling was accurate across the length of sequencing reads (Phred scores ≥ 30), with ≤ 4% undetermined “N” nucleotides at any base position; that all reads were of the expected 50-bp length; and that the reads show expected G-C content (∼50% across all reads).

Both ends of sequenced cDNA fragments were aligned to the *Homo sapiens* Genome Reference Consortium Human Build 38 (GRCh38/hg38) reference genome (Schneider *et al*., 2017) as read pairs using STAR (Version 2.7.3; (Dobin *et al*., 2013)) using GTF gene and transcript annotation files from *Ensembl* (GRCh38/hg38 reference genome release 103). A genome index file was generated using the “*genomeGenerate*” function, and the “*sjdbOverhang*” option set to a length value of 49 bases. Both ends of the sequenced cDNA fragments were aligned to the reference genome as read pairs, and *MultiQC* was used to visualize the success of the alignment. Across all samples, ∼80% of reads mapped to unique genomic loci, with the remaining reads either mapping to multiple loci or not successfully mapped due to lack of overlap with the GRCh38/hg38 reference (these unmapped reads were excluded from downstream analysis). Following alignment, SAMtools (Version 1.9; (Li *et al*., 2009)) was used to generate BAM files for the STAR alignment with the “*index*” function, and the aligned reads sorted by read name rather than coordinate using the “*sort*” function. *featureCounts* (Version 2.0.0; (Liao, Smyth and Shi, 2014)) was used to quantify the number of sequenced reads that mapped to a given gene. Transcripts with a mean count across all samples < 10 were removed.

The *DESeq2* package (Version 1.30.1; (Love, Huber and Anders, 2014)) was then used to identify gene expression changes in neuronal cells specific to JAKMIP1 deficiency. Read counts were normalized by the regularized-logarithmic transformation method (“*rLogTransformation*” function). Principal component analyses (PCA) were first conducted with the “*prcomp*” function to check for similarities and differences between samples and check for sources of variance. Statistical significance for the differential gene expression analyses were calculated for using the Wald test, with Benjamini-Hochberg false discovery rate (FDR) correction for multiple testing. Differentially expressed genes (DEGs) with an adjusted P-value < 0.05 were considered statistically significant. To minimize potentially confounding off-target effects from CRISPR gene-editing and obtain a list of DEGs more likely to reflect the true effects of JAKMIP1 deficiency in SH-SY5Y cells, only statistically significant DEGs identified in separate EV against KO-A and EV against KO-B analyses were accepted for the final DEG list (termed “overlapping” significant DEGs; compiled with the “*venn.diagram*” function).

The RNA-seq FASTQ files generated and analyzed in this study can be found at NCBI Sequence Read Archive (BioProject ID: PRJNA1467645).

### Co-immunoprecipitation (co-IP), immunoprecipitation mass spectrometry (IP-MS) and analysis of liquid chromatography-mass spectrometry (LC-MS) data

SH-SY5Y cells were maintained in complete medium and grown to 80-90% confluency prior to transfection with Lipofectamine™ LTX. For each SH-SY5Y cell co-IP/IP-MS experiment, the combined total SH-SY5Y cells from two 10-cm-diameter Petri dishes were pooled by centrifugation into a single pellet due to relatively low transfection efficiency (∼20-60% depending on the plasmid of interest).

48 hours post-transfection, cells were dissociated with TrypLE™, washed with ice-cold PBS, and re-suspended in IP-MS lysis buffer supplemented with PMSF, Protease Inhibitor Cocktail, and Phosphatase Inhibitor Cocktails. Prior to immunoprecipitation, lysates were cleared by centrifugation at 4°C. mCherry-tagged JAKMIP1-containing protein complexes were immunoprecipitated with RFP-Trap® Agarose beads (#rta-20, *Proteintech*) (see File S3 for full immunoprecipitation procedure). For co-IP experiments, immunocomplexes bound to the beads were dissociated by heating in Laemmli buffer for 5 minutes at 95°C, after which Western blotting was performed on the supernatant to analyze proteins that co-precipitated with each JAKMIP1 construct. For the IP-MS experiments, immunocomplexes were not dissociated and instead left bound to the RFP-Trap® Agarose beads, which were then shipped to the *University of Bristol Proteomics Facility* for LC-MS. Each IP-MS experiment was performed in duplicate, producing replicates “A” and “B”, which signify two separate sets of mass spectrometry-identified peptides from proteins that co-precipitated with the mCherry-tagged JAKMIP1 constructs (or the mCherry control) from two independently performed immunoprecipitation reactions.

LC-MS and subsequent analysis of LC-MS data with *Proteome Discoverer* (*Thermo Fisher Scientific*) was performed by the *University of Bristol Proteomics Facility*. Peptides from immunoprecipitated proteins were identified with a SEQUEST search against the *UniProt* human proteome. Only protein hits with a minimum 5% FDR were included in the final list of identified proteins. Final lists of proteins that co-precipitated with the mCherry-tagged constructs were provided by the *University of Bristol Proteomics Facility*. Several pieces of information accompanied each protein ‘hit’, including unique *UniProtKB* accession IDs, sequence coverage of the protein by detected peptides; number of distinct peptide sequences identified; the number of peptide spectrum matches; whether the peptide is a common contaminant routinely observed in similar proteomics experiments; and a “Score Sequest HT” value, which is calculated from the sum of the scores of individual peptides – the higher the “Score Sequest HT” value, the better the quality of protein detection.

A filtering strategy was then implemented to remove ‘sticky’, contaminating proteins that were non-specifically pulled down by either the overexpressed mCherry protein or the RFP-Trap® Agarose beads, as well as to identify the likely ‘true’ JAKMIP1-interacting proteins (the full strategy is explained in File S3). In brief, proteins that co-precipitated with both replicates of the mCherry control were compiled to create a list of the non-specific, sticky proteins. If a protein co-precipitated with an mCherry-tagged JAKMIP1 construct and was not in the list of sticky proteins, then it was considered a likely true JAKMIP1 interactor. Furthermore, if a protein was also in the sticky list but showed a ≥ 2.5-fold increase in Score Sequest HT value when pulled down with a JAKMIP1 construct, then it was also considered a likely true JAKMIP1 interactor. If a likely JAKMIP1-interacting protein was detected in both A and B replicates, then it was classed as a “Double-Positive” or “High-Confidence” putative JAKMIP1-interacting protein. If a potential interacting protein was only detected in one of the A or B replicates, then it was classed as a “Single-Positive” or “Low-Confidence” putative JAKMIP1-interacting protein. These putative JAKMIP1-interacting proteins were then validated by independent co-IP experiments.

### Gene ontology pathway and gene-set over-representation analyses

Functional gene ontology (GO) pathway enrichment analyses were performed to infer cellular pathways that may be regulated by JAKMIP1 in neuronal cells. This was performed on the list overlapping significant DEGs with the *g:GOSt* tool from *g:Profiler* (available at https://biit.cs.ut.ee/gprofiler/gost) (Raudvere *et al*., 2019). GO pathway enrichment analyses were only conducted with annotated genes against the GO Biological Process 2021 database and a statistical significance threshold of adjusted P-value < 0.05 was set, with Benjamini-Hochberg FDR correction for multiple testing (final q < 0.05 was accepted as statistically significant). The GO pathway enrichment analyses were carried out separately for up- and downregulated DEGs as suggested in (Hong *et al*., 2014).

The list of overlapping significant DEGs were also subjected to gene-set over-representation analyses (GSOA). GSOA was performed to test several hypotheses, specifically 1) whether JAKMIP1-regulated genes are enriched for ASD risk genes; 2) whether there exists a subset of genes co-regulated by JAKMIP1, FMRP and CYFIP1; and 3) whether there exist overlapping gene expression signatures between *JAKMIP1*^−/−^ SH-SY5Y cells and human models of maternal immune activation. Where possible, gene sets were obtained from studies utilizing neural models (such as NPCs and neurons) and ideally, of human origin. Though neuronal models are preferred to compare with the population of differentiated SH-SY5Y cells, NPCs can still offer valuable perspective on earlier neurodevelopment and stages of neuronal differentiation. Details of the datasets chosen for GSOA are provided in File S3. GSOA were performed in *RStudio*. Two-by-two contingency tables were first constructed, comparing frequencies of overlapping and non-overlapping DEGs from *JAKMIP1*^−/−^ SH-SY5Y cells and genes/DEGs from each dataset (these were matched and filtered by *Ensembl* gene ID as opposed to gene symbol to prevent mistakes caused by previous or alias gene symbols across different publications) against a background gene set of the entirety of human genes in the GRCh38/hg38 reference genome (release 103 from *Ensembl*). Enrichment of different gene sets was carried out with one-sided Fisher’s exact tests (the “*fisher.test*” function), performed by specifying the “*alternative*” argument to “*greater*” (i.e., testing whether the odds ratio is greater than 1). P-values were then adjusted by Benjamini-Hochberg FDR correction with the “*p.adjust*” function (final q < 0.05 was accepted as statistically significant). Enrichment analyses were carried out separately for up- and downregulated DEGs identified in the *JAKMIP1*^−/−^ SH-SY5Y cells.

In parallel with the DEGs, the list of putative JAKMIP1-interacting proteins from the IP-MS experiments were subjected to similar GSOA as well to test whether JAKMIP1 appears to be more likely to interact with proteins encoded by ASD candidate genes (from *SFARI Gene* (Abrahams *et al*., 2013)). This was performed in the same manner as described above.

### Statistical Analyses

For all wet laboratory experiments, data were analyzed, and statistical significance determined using *GraphPad Prism* (GraphPad Prism version 9.0.0 for Windows 11, GraphPad Software, San Diego, California United States, www.graphpad.com).

The differences between any two groups were assessed by unpaired student’s t-tests. The differences between multiple groups were assessed by ordinary one-way ANOVA with Tukey’s HSD tests for multiple comparisons with the exception of neurite tracing. Neurite lengths are not typically normally distributed (and can be seen in this study by skewness of the range of data in box and whisker plots in Fig. 3 and 9), and hence these data do not satisfy the assumption of data normality required for parametric tests such as ANOVA. Instead, the Kruskal-Wallis test (the non-parametric counterpart to the ANOVA) is used to compare the medians of neurite length measurements (followed by Dunn’s multiple comparisons test). Box and whisker plots are presented to provide a clearer picture of the distribution of neurite lengths. The whiskers of these plots extend to the 5^th^ and 95^th^ percentiles of the data, with neurite length measurements outside these percentiles displayed as individual ‘outlier’ data points.

The differences between grouped data sets where the influence of two different variables and the interaction between them is in question (e.g., both JAKMIP1 deficiency and IL-6 treatment and their impact on neurite outgrowth), a two-way ANOVA is used as there is no suitable non-parametric counterpart that examines interaction effects. This is followed by Tukey’s HSD tests for multiple comparisons.

## Supplementary Figures (File S1)

**Supplementary Figure 1.**
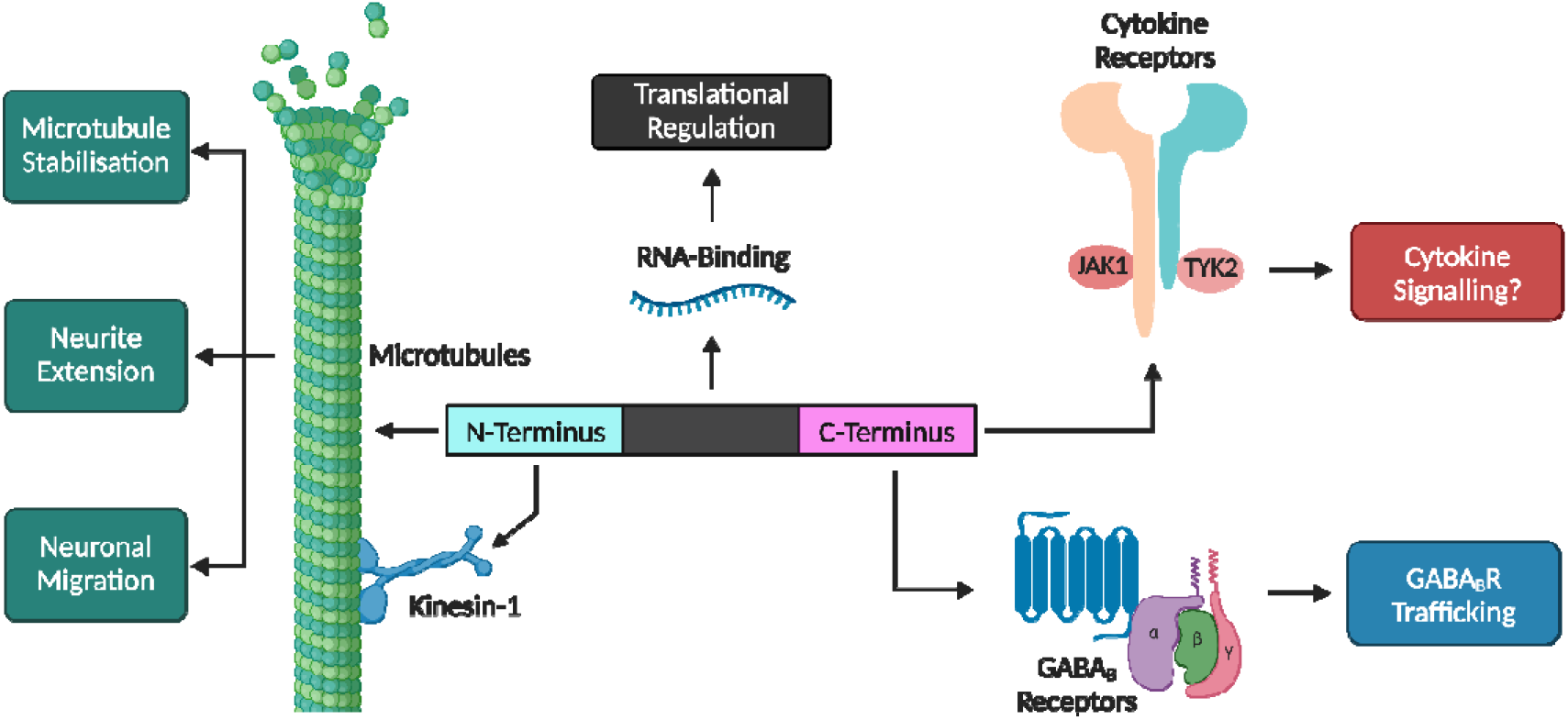
Distinct functional domains of the JAKMIP1 protein. Schematic overview of the known functional domains of JAKMIP1 as well as their interactions. The JAKMIP1 N-terminus associates with microtubules and motor proteins; a central region interacts with mRNAs and is involved in translational regulation; and the C-terminus interacts with JAKs and GABA_B_R1. Adapted from (Steindler *et al*., 2004; Vidal *et al*., 2007, 2012; Berg *et al*., 2015). Created with BioRender.com.

**Supplementary Figure 2.**
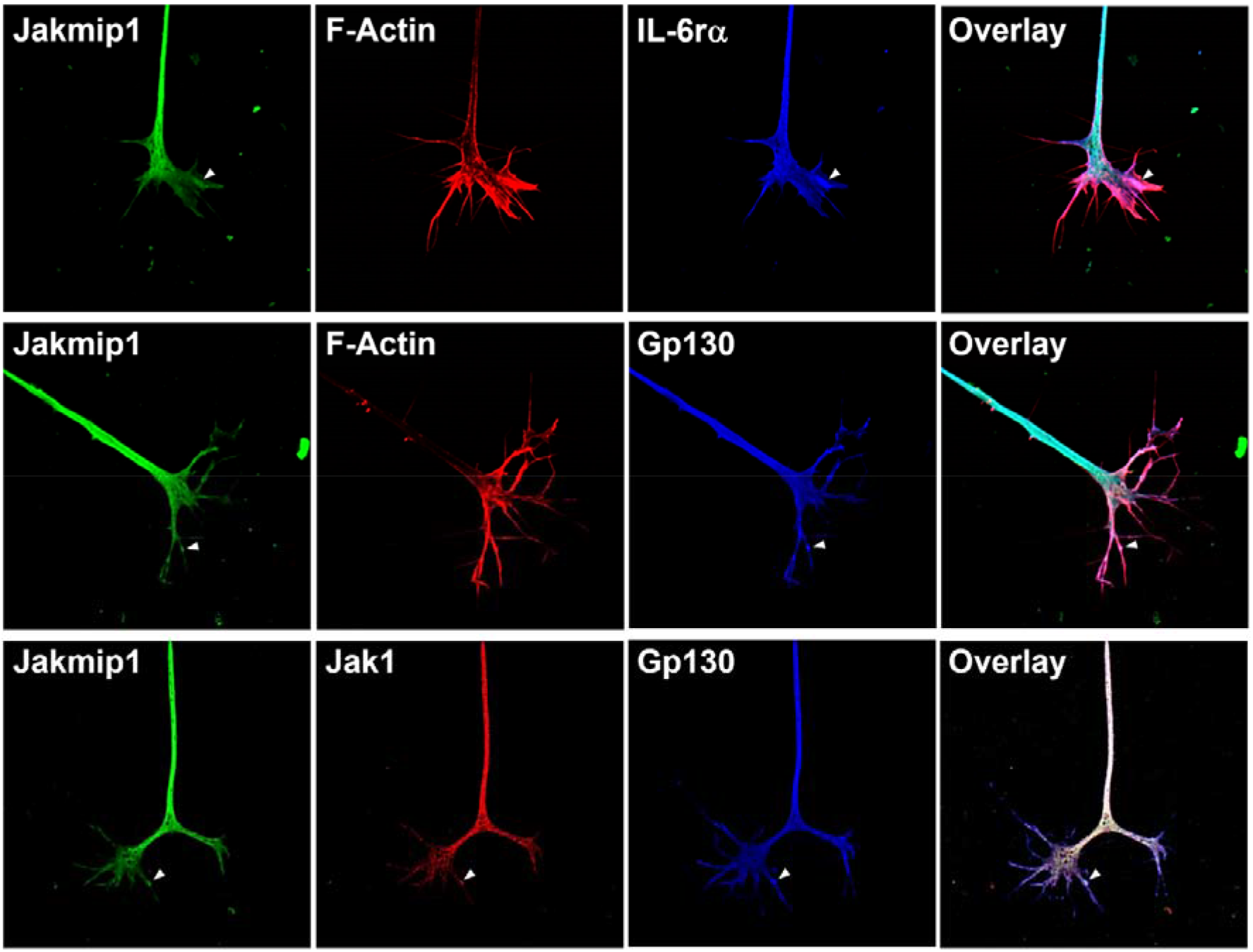
Jakmip1 co-localizes with components of the IL-6r complex in chick neurons. Chick dorsal root ganglion neurons were explanted at embryonic day 7 and cultured in 100 ng/mL IL-6, fixed, permeabilized and stained for varying combinations of Jakmip1, IL-6rα, Gp130, Jak1 and F-Actin. Arrowheads indicate examples of co-localization between Jakmip1 and IL-6rα, Gp130 or Jak1.

**Supplementary Figure 3.**
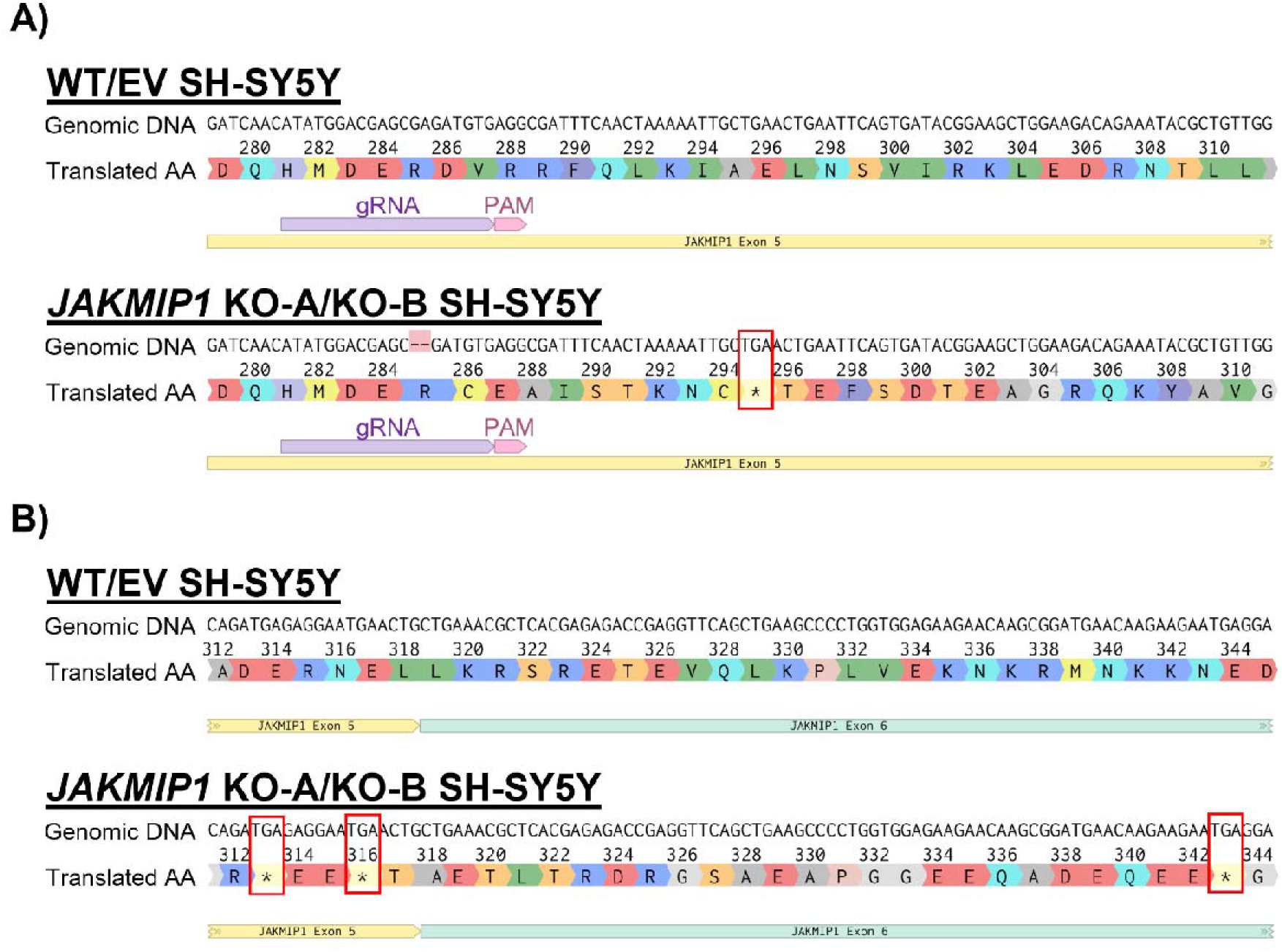
Sanger sequencing reveals 2-bp deletion in *JAKMIP1*^−/−^ SH-SY5Y cells. **A)** Visualization of Sanger sequencing results from *GENEWIZ*, showing the genomic DNA sequence of WT, EV control and both *JAKMIP1*^−/−^ SH-SY5Y cell lines (named KO-A and KO-B) and the subsequent translated amino acid sequence. A 2-bp deletion is identified (mismatch in DNA alignment shown in red) within the gRNA sequence (in purple) in exon 5 of the *JAKMIP1* gene following clonal isolation and expansion. This disrupts the reading frame of the gene, leading to a premature TGA (i.e., UGA in mRNA) stop codon (shown by an asterisk in the amino acid sequence) instead of an amino acid at position 295 in the KO-A and KO-B cell lines. Stop codons are also outlined with red rectangles for clarity. **B)** Effects of the 2-bp deletion shown in part A) on the downstream amino acid sequence of JAKMIP1. Additional UGA stop codons are found encoded towards the end of exon 5 (amino acid^313^ and amino acid^316^) and in exon 6 (amino acid^343^). Created with *Benchling* (Benchling [Biology Software]. (2022). Retrieved from https://benchling.com.). EV – empty vector; gRNA – guide RNA; KO-A – *JAKMIP1*-knockout A; KO-B – *JAKMIP1*-knockout B; WT – wild-type.

**Supplementary Figure 4.**
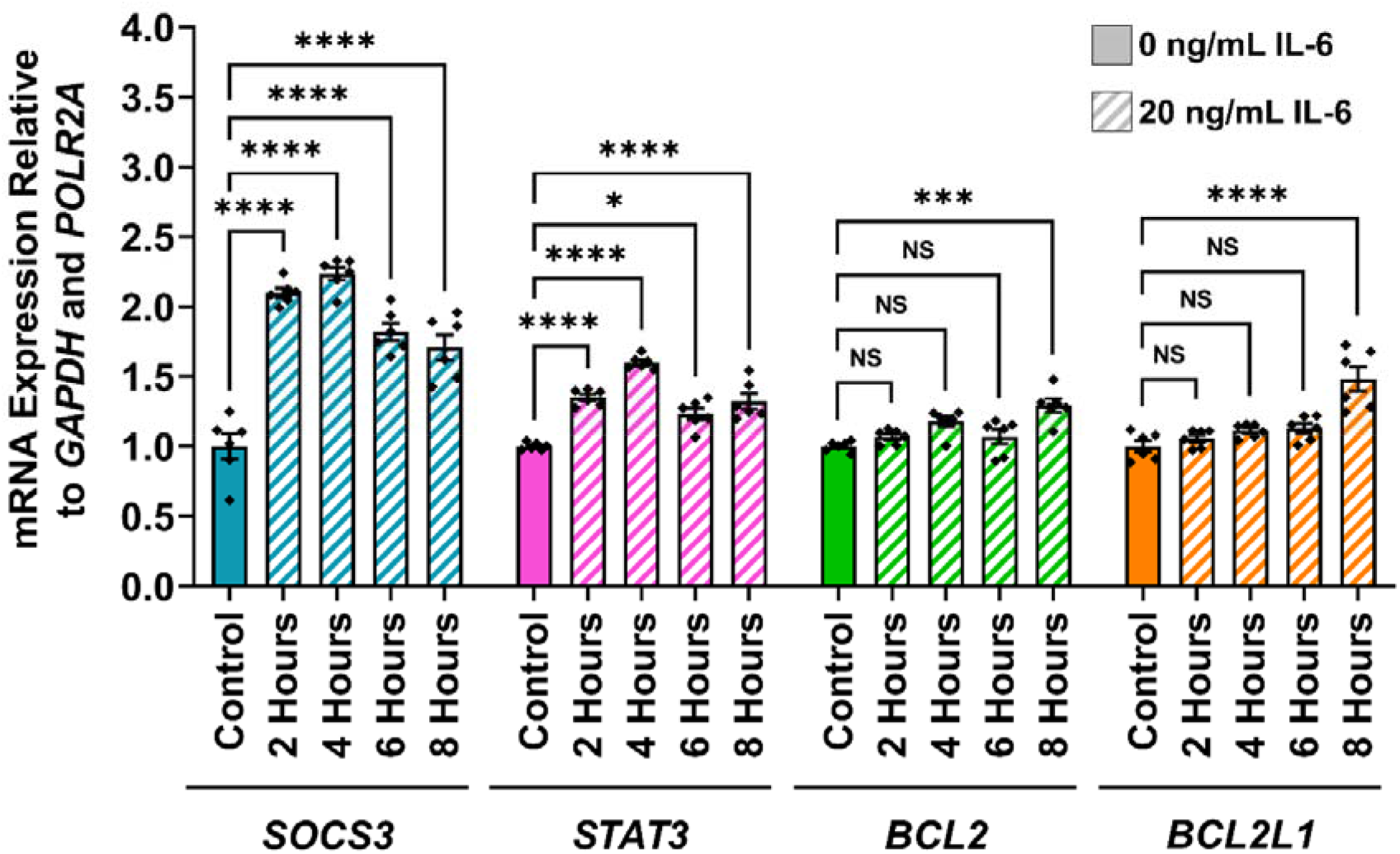
Identifying STAT3-responsive genes induced by IL-6 treatment in SH-SY5Y cells. Relative mRNA expression of typical STAT3-target genes (*SOCS3*, *STAT3*, *BCL2* and *BCL2L1*) in WT SH-SY5Y cells following treatment with 20 ng/mL IL-6 for various timepoints indicated (2-8 hours), measured by qRT-PCR. Cells treated with complete medium (i.e., culture medium was refreshed) for 8 hours were used as a control. All values are normalized to the non-IL-6-treated control. Values presented as mean ± SEM of N = 2 independent experiments, 3 replicates each. One-way ANOVA with Tukey’s HSD test for multiple comparisons. Statistical significance displayed against the non-IL-6-treated control; NS – not significant (P > 0.05); ***P < 0.001; ****P < 0.0001). ANOVA – analysis of variance; mRNA – messenger RNA; qRT-PCR – quantitative reverse transcription PCR; SEM – standard error of the mean; WT – wild-type.

**Supplementary Figure 5.**
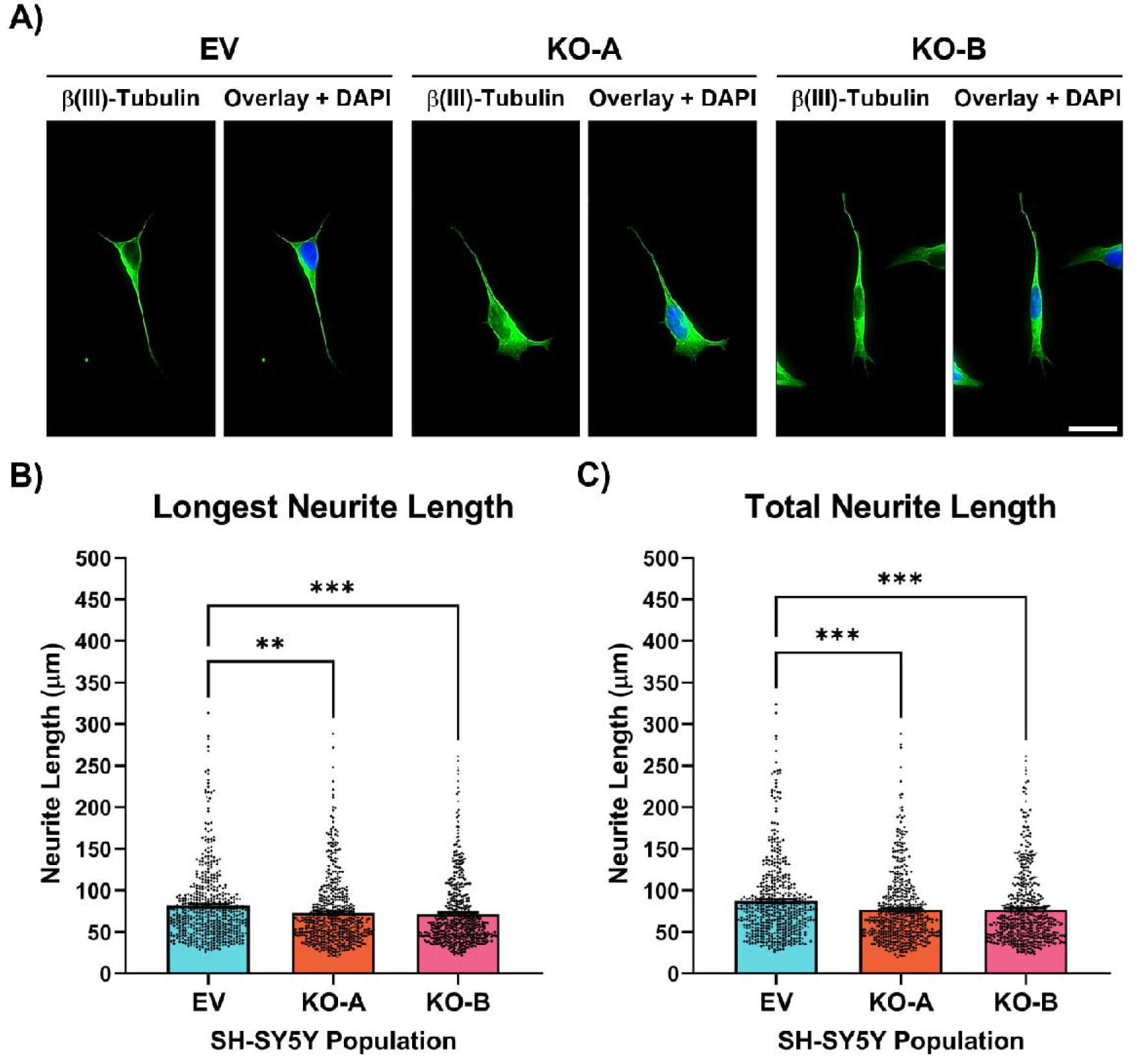
JAKMIP1-/- SH-SY5Y cells display subtle deficits in neurite extension and neuritogenesis. **A)** EV, KO-A and KO-B SH-SY5Y cells were differentiated on PDL-coated glass coverslips prior to fixation and immunostaining for β(III)-Tubulin (Alexa Fluor® 488; seen in green). Images representative of N = 3 independent differentiations, 2 separate coverslips per differentiation experiment and 515-522 SH-SY5Y cells traced per genotype. Scale bar = 50 μm. B) Measurement of the length of the longest neurite (LNL) produced by individual SH-SY5Y cells in part A). Kruskal-Wallis with Dunn’s multiple comparisons test for multiple comparisons. Statistical significance displayed against the EV control; NS – not significant (P > 0.05); ***P < 0.001; ****P < 0.0001. C) Measurement of the sum length of all neurites (TNL) produced by individual SH-SY5Y cells in part A). Kruskal-Wallis with Dunn’s multiple comparisons test for multiple comparisons. Statistical significance displayed as in B). EV – empty vector; KO-A – *JAKMIP1*-knockout A; KO-B – *JAKMIP1*-knockout B; PDL – poly-D-Lysine.

**Supplementary Figure 6.**
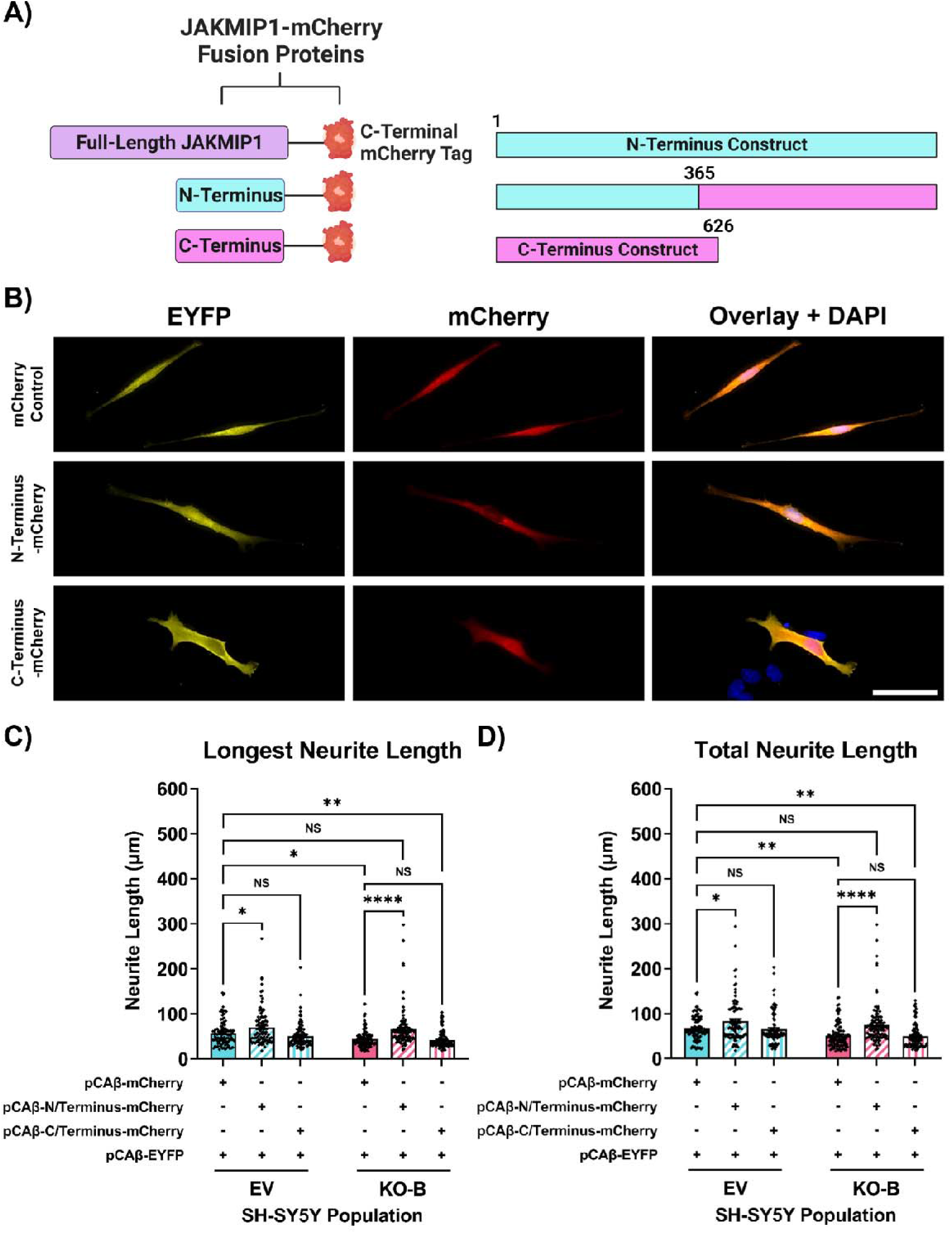
The JAKMIP1 N-terminus is responsible for the neurite outgrowth-promoting effects of JAKMIP1. **A)** Left: schematic representation of JAKMIP1-mCherry fusion proteins, including the full-length and truncated N- and C-termini domains. Right: Outline of amino acid composition of truncated JAKMIP1 constructs. The N-terminus is composed of amino acid^1-365^, and the C-terminus is composed of amino acid^365-626^. Created with BioRender.com. **B)** Representative fluorescence micrographs of differentiated EV and KO-B SH-SY5Y cells transfected with plasmid vectors encoding truncated JAKMIP1-mCherry fusion constructs (in conjunction with a separate vector encoding EYFP for neurite tracing) by Nucleofection™. Transfected SH-SY5Y cells were differentiated for 48 hours before fixation. Images representative of N = 3 independent differentiations, 2 separate coverslips per differentiation experiment and 97-110 SH-SY5Y cells traced per genotype/transfection combination. Scale bar = 50 μm. **C)** Measurement of the length of the longest neurite (LNL) produced by individual transfected SH-SY5Y cells in B). Two-way ANOVA with Tukey’s HSD test for multiple comparisons. Statistical significance displayed against the mCherry-OE EV control; NS – not significant (P > 0.05); *P < 0.05; **P < 0.01; ***P < 0.001; ****P < 0.0001. **D)** Measurement of the sum length of all neurites (TNL) produced by individual SH-SY5Y cells in B). Two-way ANOVA with Tukey’s HSD test for multiple comparisons. Statistical significance displayed as in C). ANOVA – analysis of variance; EV – empty vector; EYFP – enhanced yellow fluorescent protein; KO-B – *JAKMIP1*-knockout B; OE – overexpression; PDL – poly-D-Lysine.

**Supplementary Figure 7.**
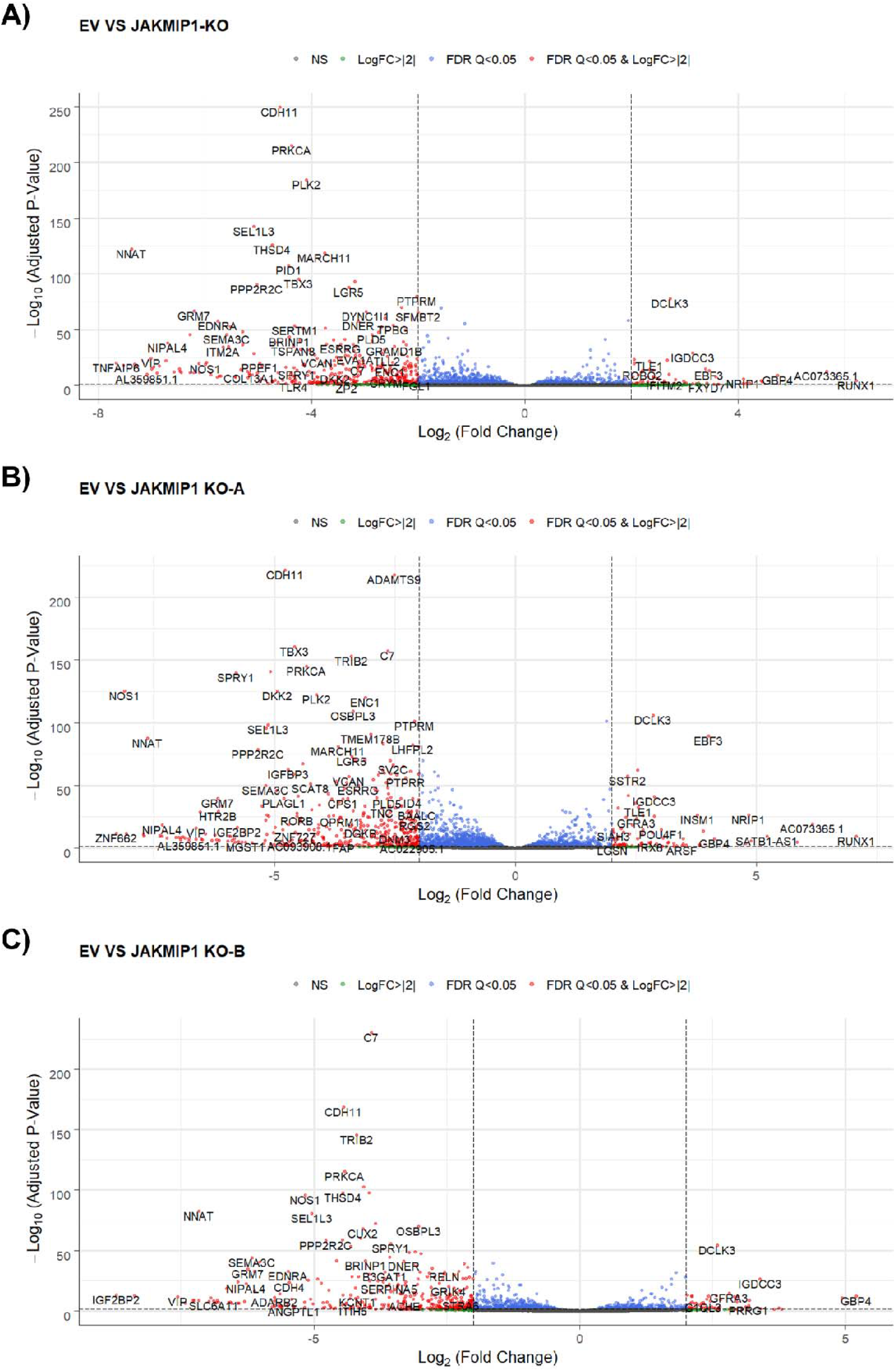
Global gene expression changes in differentiated *JAKMIP1*^−/−^ SH-SY5Y cells. Volcano plots for DEGs identified in **A)** both KO-A and KO-B combined; or **B)** KO-A only; or **C)** KO-B only compared to the EV control. Grey dots = non-significant DEGs (adjusted P-value > 0.05); green dots = significant DEGs (adjusted P-value < 0.05) with an absolute Log_2_FC ≥ 2; blue dots = significant DEGs (adjusted P-value < 0.05) with Q-value following Benjamini-Hochberg FDR correction ≤ 0.05; red dots = significant DEGs with both an absolute Log_2_FC ≥ 2 and Q-value following Benjamini-Hochberg FDR correction ≤ 0.05. DEG – differentially expressed gene; EV – empty vector control; FDR – false discovery rate; KO-A – *JAKMIP1*-knockout A; KO-B – *JAKMIP1*-knockout B; Log_2_FC – base 2 logarithm of differential gene expression fold change (FC).

**Supplementary Figure 8.**
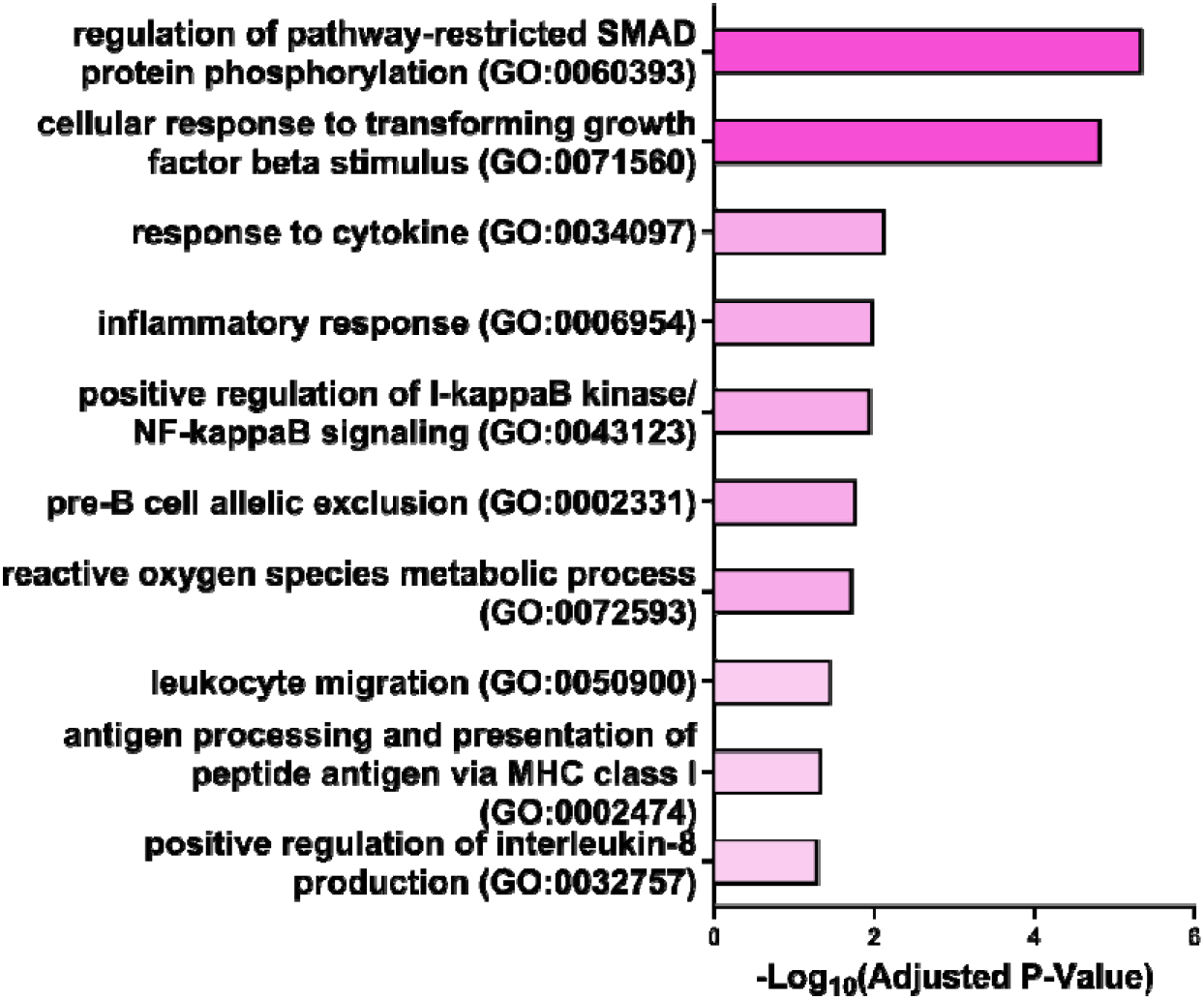
Enrichment of cytokine signaling, inflammation and immune function-related GO Biological Process 2021 terms in downregulated DEGs. Additional results of GO analysis performed on the separated downregulated DEGs relating to cytokine signaling, inflammation and immune cell function. 10 terms of interest are reported with the adjusted P-value following Benjamini-Hochberg FDR correction for multiple testing. Terms are sorted by significance of adjusted P-value. DEG – differentially expressed gene; FDR – false discovery rate.

**Supplementary Figure 9.**
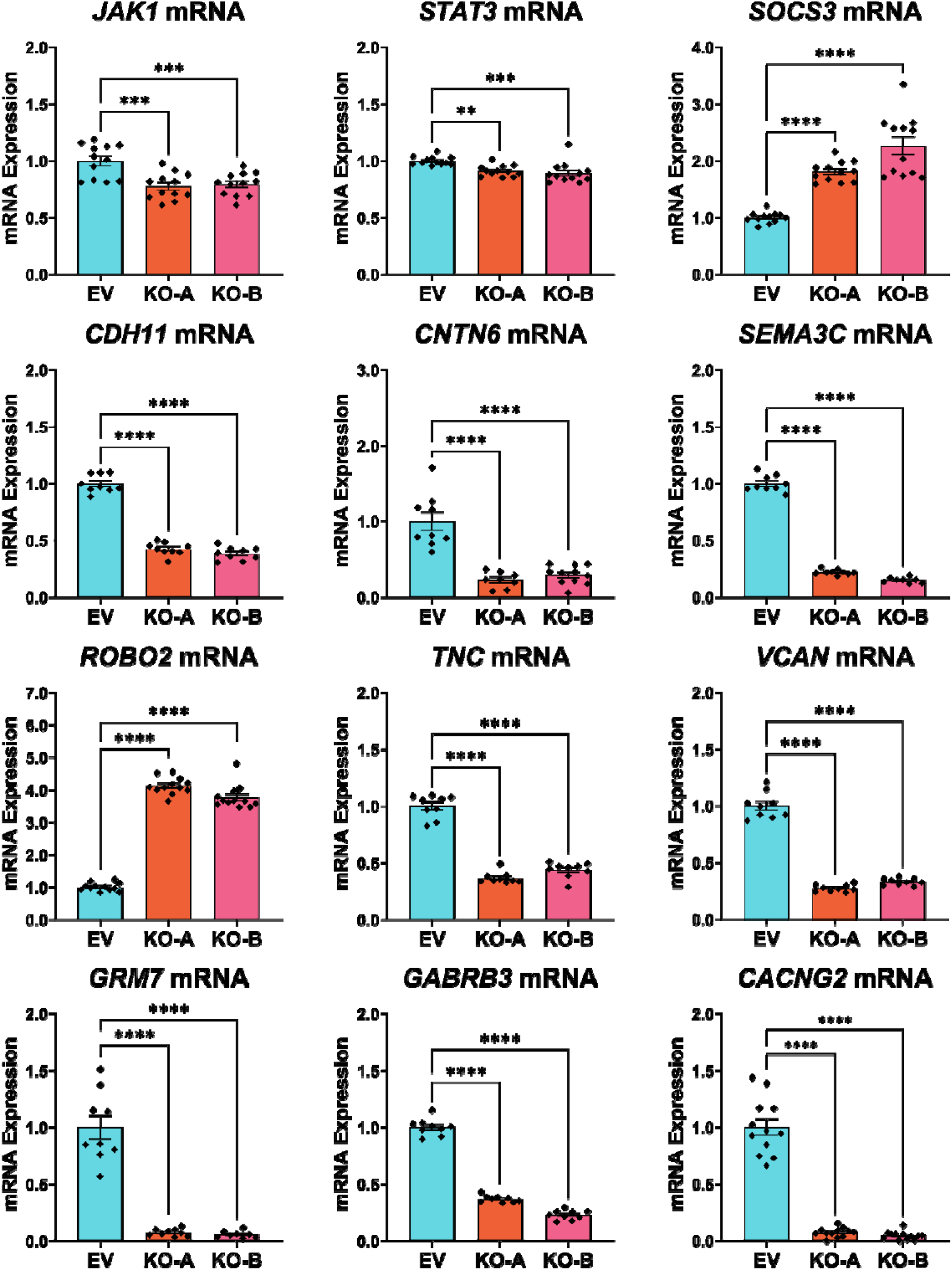
Validation of differential gene expression analysis by qRT-PCR. Relative mRNA expression of various identified DEGs from the RNA-seq in differentiated EV, KO-A and KO-B SH-SY5Y cells measured by qRT-PCR. DEGs fall into several broad categories, namely IL-6/JAK1/STAT signaling components (*JAK1*, *STAT3* and *SOCS3*); CAMs (*CDH11* and *CNTN6*); axon guidance ligands and receptors (*SEMA3C* and *ROBO2*); ECM components (*TNC* and *VCAN*); and synaptic receptors and ion channels (*GRM7*, *GABRB3* and *CACNG2*). *GAPDH* and *POLR2A* were used as housekeeping genes for relative mRNA expression quantification. All values are normalized to the EV control. Values presented as mean ± SEM of N = 3 independent experiments, 3-4 replicates each. Statistical significance displayed against the EV control; **P < 0.01; ***P < 0.001; ****P < 0.0001. One-way ANOVA with Tukey’s HSD test for multiple comparisons. DEG – differentially expressed gene; EV – empty vector control; KO-A – *JAKMIP1*-knockout A; KO-B – *JAKMIP1*-knockout B; mRNA – messenger RNA; RNA-seq – RNA sequencing; qRT-PCR – quantitative reverse transcription polymerase chain reaction; SEM – standard error of the mean.

**Supplementary Figure 10.**
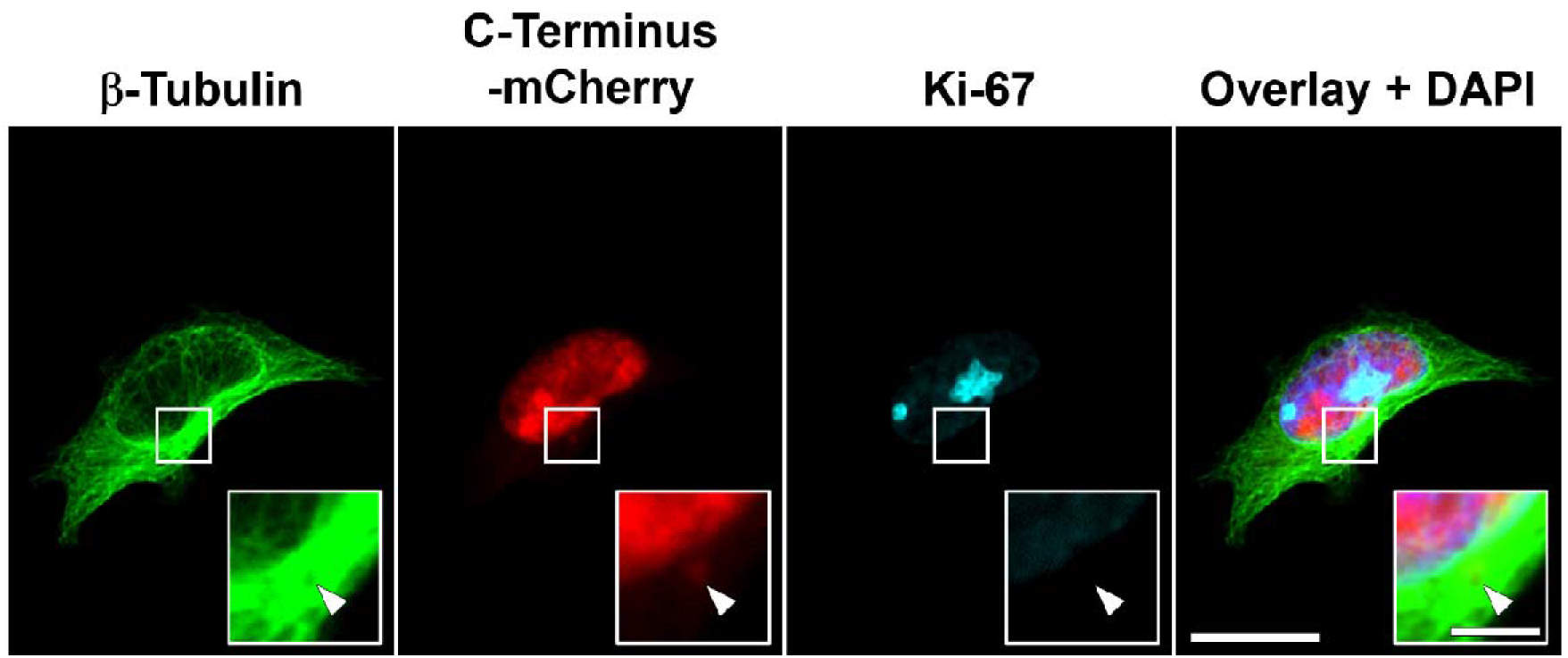
The JAKMIP1 C-terminus can display centrosomal localization. HEK-293 cells transfected with the mCherry-tagged JAKMIP1 C-terminus construct. Immunostaining for β-Tubulin and Ki-67. Scale bar = 20 μm. Insets show enlarged view of regions of interest outlined with white squares. Arrowhead indicates centrosomal localization of the JAKMIP1 C-terminus construct observed in a subset of transfected cells. Inset scale bar = 5 μm.

**Supplementary Figure 11.**
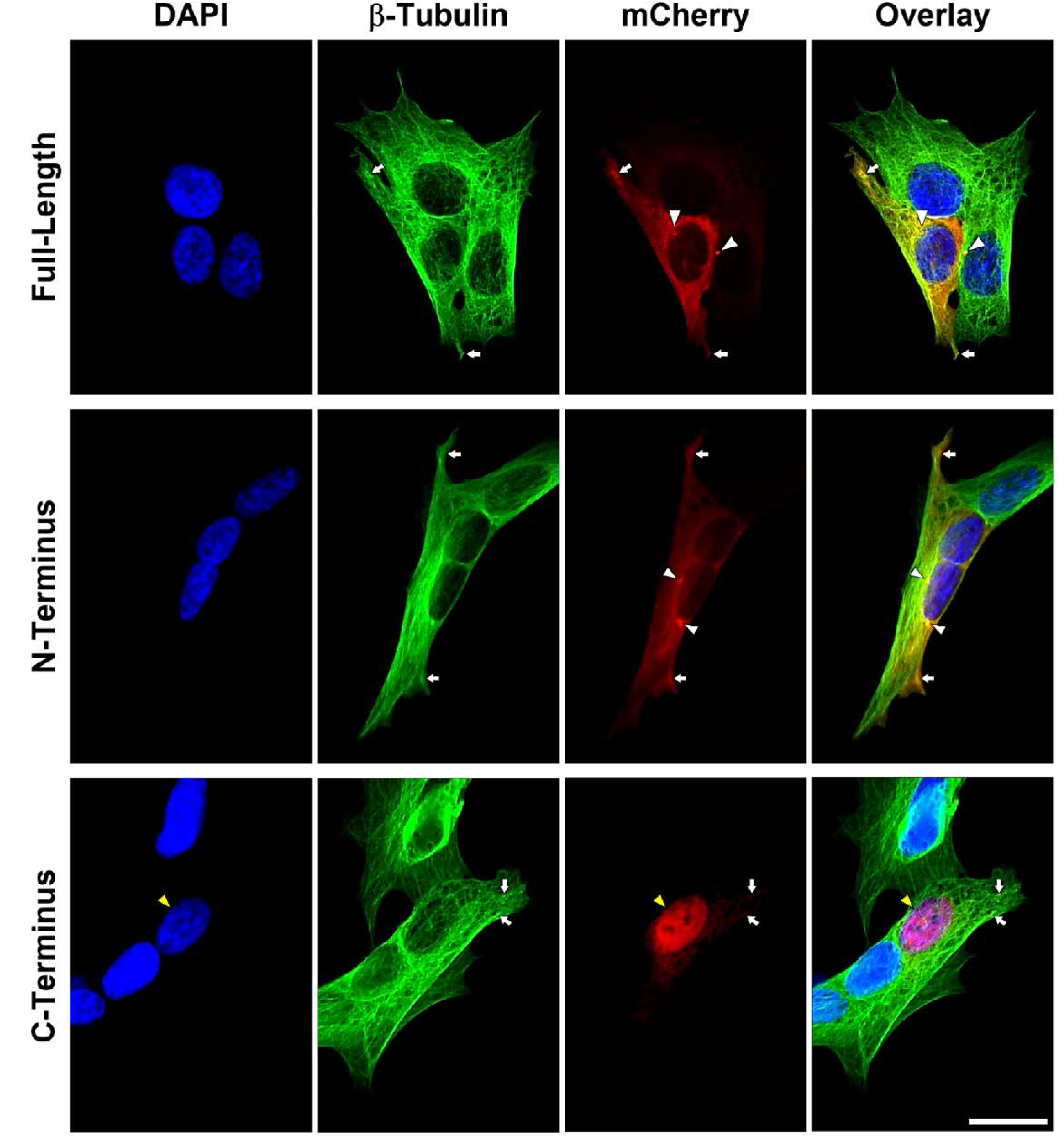
The JAKMIP1 C-terminus displays predominantly nuclear localization. Representative fluorescence micrographs of SH-SY5Y cells transfected with full-length or truncated JAKMIP1 constructs tagged with mCherry (using jetOPTIMUS®). Images representative of N = 3 independently fixed plates of cells, 1 separate coverslip stained per plate. Transfected cells were fixed 48 hours post transfection. Immunostaining for β-Tubulin (Alexa Fluor® 488; seen in green). White arrowheads indicate juxtanuclear puncta visible in both the full-length and N-terminus constructs; white arrows indicate co-localization with microtubules; and yellow arrowheads indicate nuclear localization of the C-terminus construct. Scale bar = 20 μm. PDL – poly-D-Lysine; WT – wild-type.

**Supplementary Figure 12.**
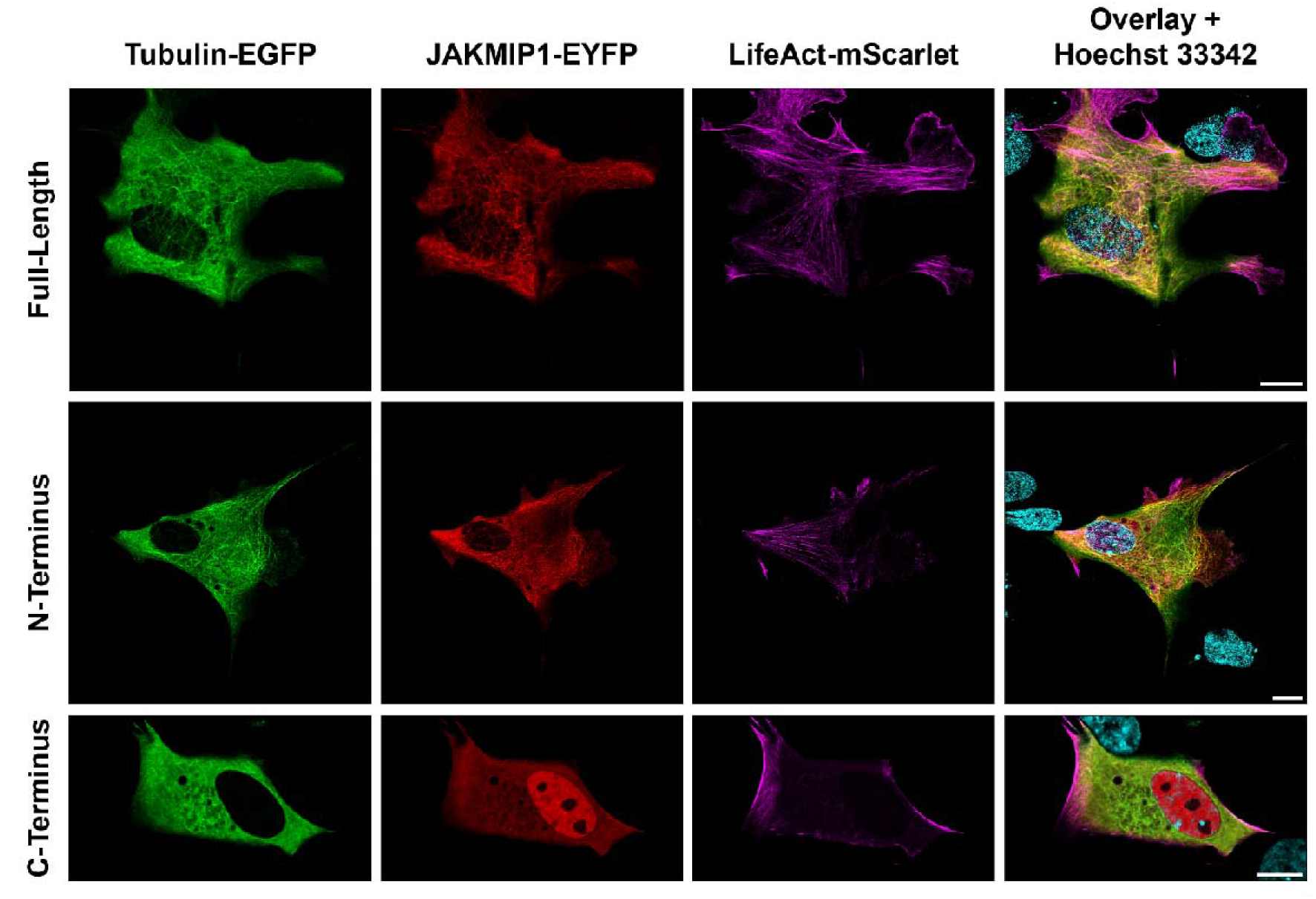
Live-cell imaging confirms nuclear localization of the JAKMIP1 C-terminus. HEK-293 cells transfected with plasmid vectors encoding EGFP-tagged Tubulin (green), EYFP-tagged full-length or truncated JAKMIP1 constructs (red) and mScarlet-tagged LifeAct (magenta) with Lipofectamine™ LTX. Nuclei were counterstained with 50 ng/mL Hoechst 33342 (cyan) for at least 30 minutes prior to imaging. Images captured with a Leica STELLARIS 8 confocal microscope by Paul McCormick. Scale bars = 10 μm.

**Supplementary Figure 13.**
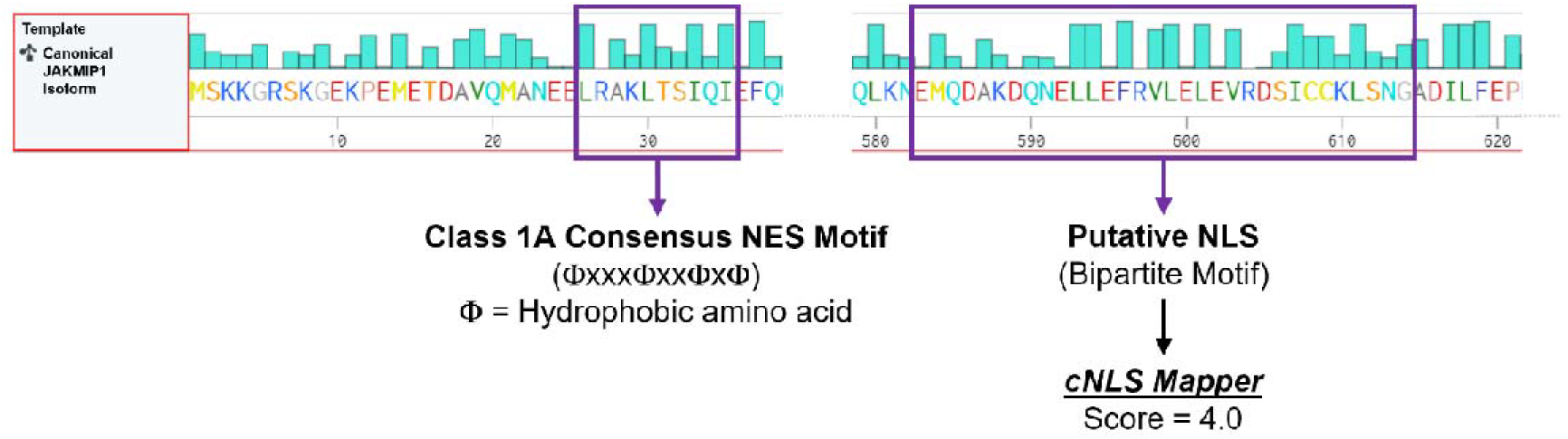
Putative nuclear localization and export sequences in JAKMIP1. Amino acid sequence of JAKMIP1. Cyan bars above each amino acid represent the hydrophobicity of the residue (longer bar = more hydrophobic). *cNLS Mapper* (available at: https://nls-mapper.iab.keio.ac.jp/cgi-bin/NLS_Mapper_form.cgi) identifies a putative bipartite NLS motif at the C-terminus of JAKMIP1 (located at amino acid^583-614^) with a score of 4.0. A class 1A consensus NES motif at the N-terminus of JAKMIP1 (located at amino acid^26-35^). Created with *Benchling* (Benchling [Biology Software]. (2022). Retrieved from https://benchling.com.). Φ – hydrophobic amino acid; NES – nuclear export sequence; NLS – nuclear localization sequence.

**Supplementary Figure 14.**
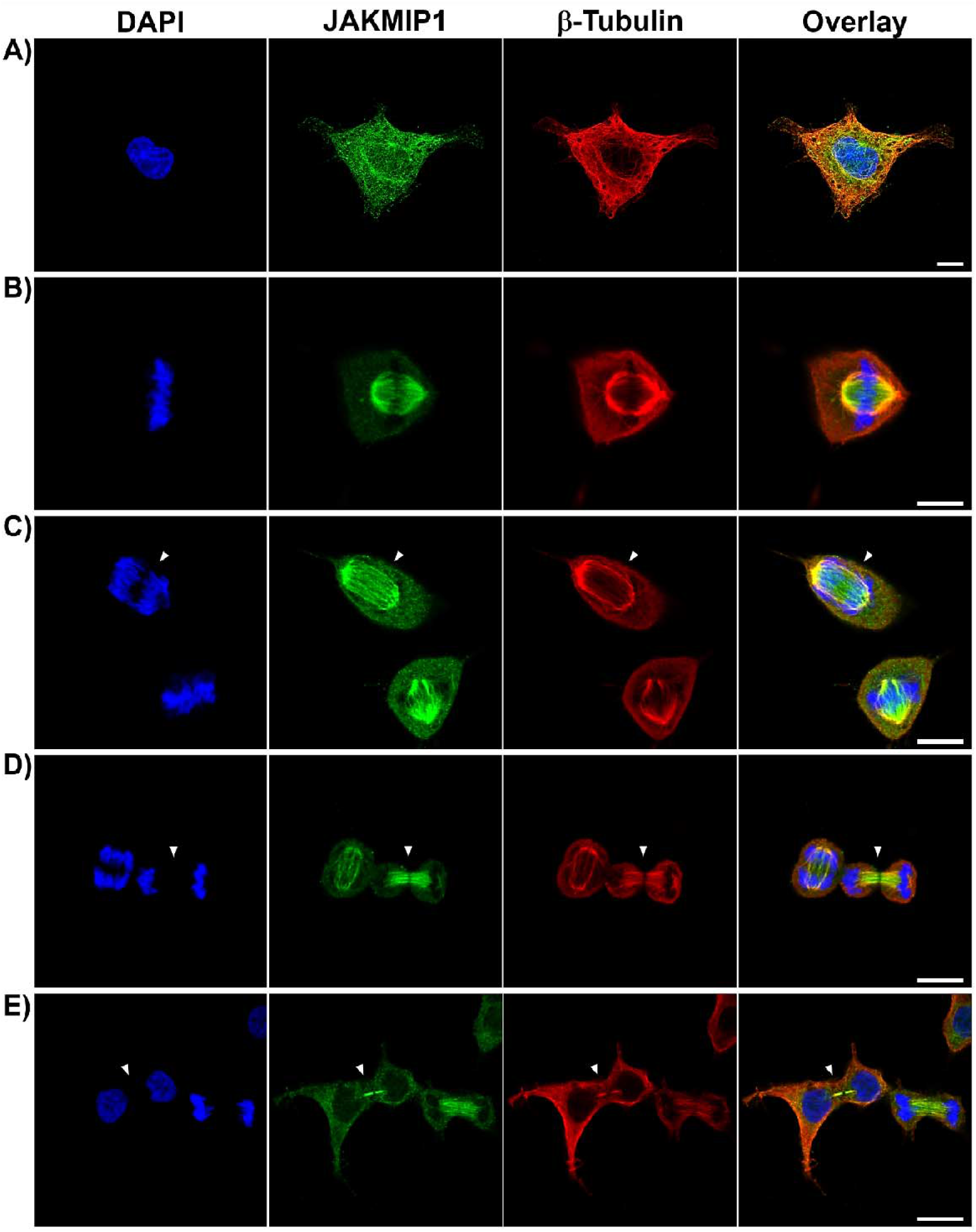
JAKMIP1 is localized to spindle microtubules during mitosis. HEK-293 cells were cultured on PDL-coated coverslips prior to fixation with PFA and immunostaining for JAKMIP1 (Alexa Fluor® 488; seen in green) and β-Tubulin (Alexa Fluor® 568; seen in red). Scale bars = 10 μm. **A)** Non-dividing HEK-293 cell in interphase. **B)** HEK-293 cell in metaphase. **C)** HEK-293 cell in early anaphase (indicated by arrowhead). **D)** HEK-293 cells in late anaphase (indicated by arrowhead. **E)** HEK-293 cells in late telophase prior to completion of cytokinesis (indicated by arrowhead). PDL – poly-D-Lysine; PFA – paraformaldehyde.

**Supplementary Figure 15.**
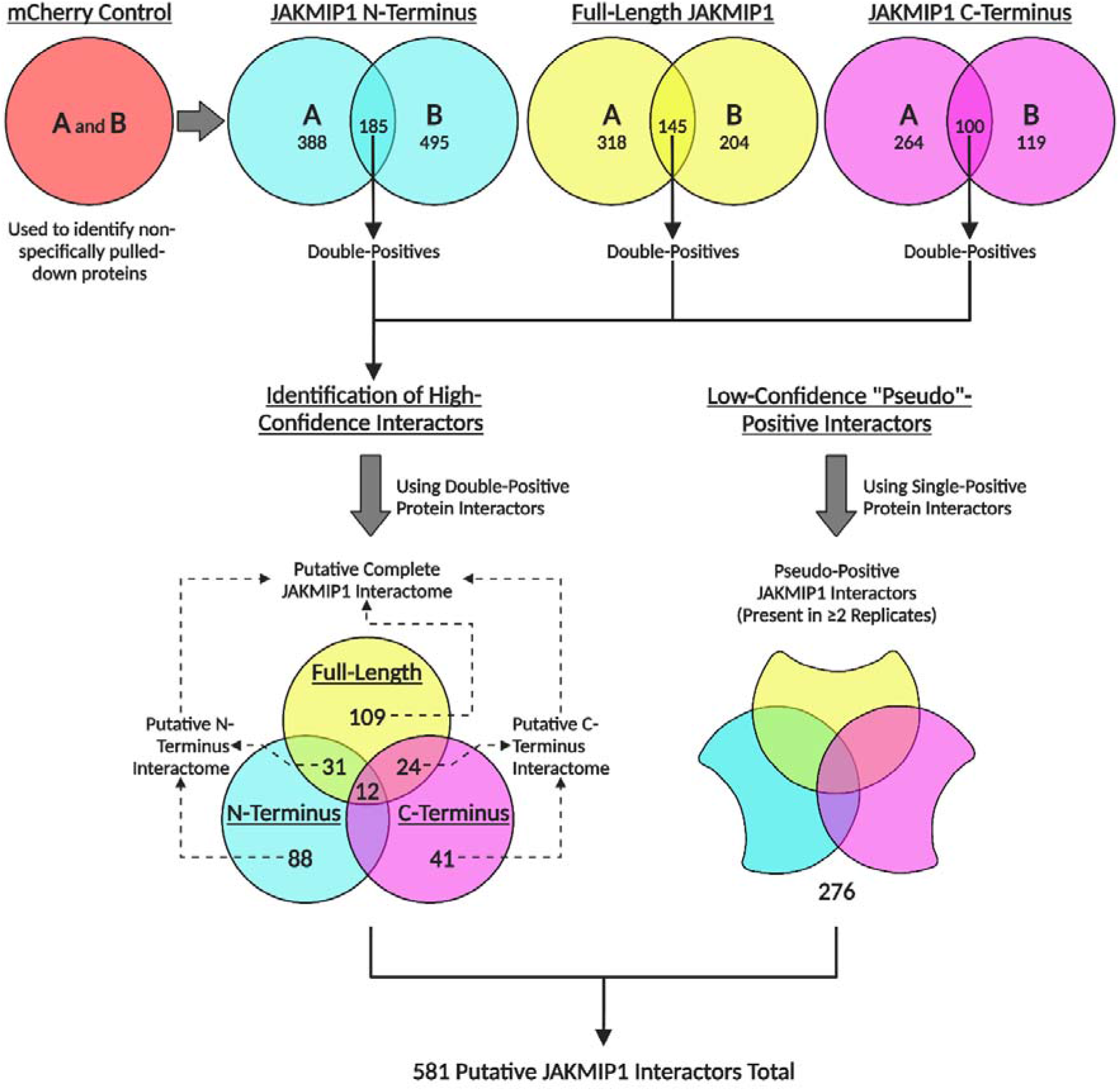
Identification of putative JAKMIP1-interacting proteins from IP-MS data. Overview of IP-MS analysis results, illustrating how the final set of 581 putative JAKMIP1-interacting proteins were identified. Created with BioRender.com.

**Supplementary Figure 16.**
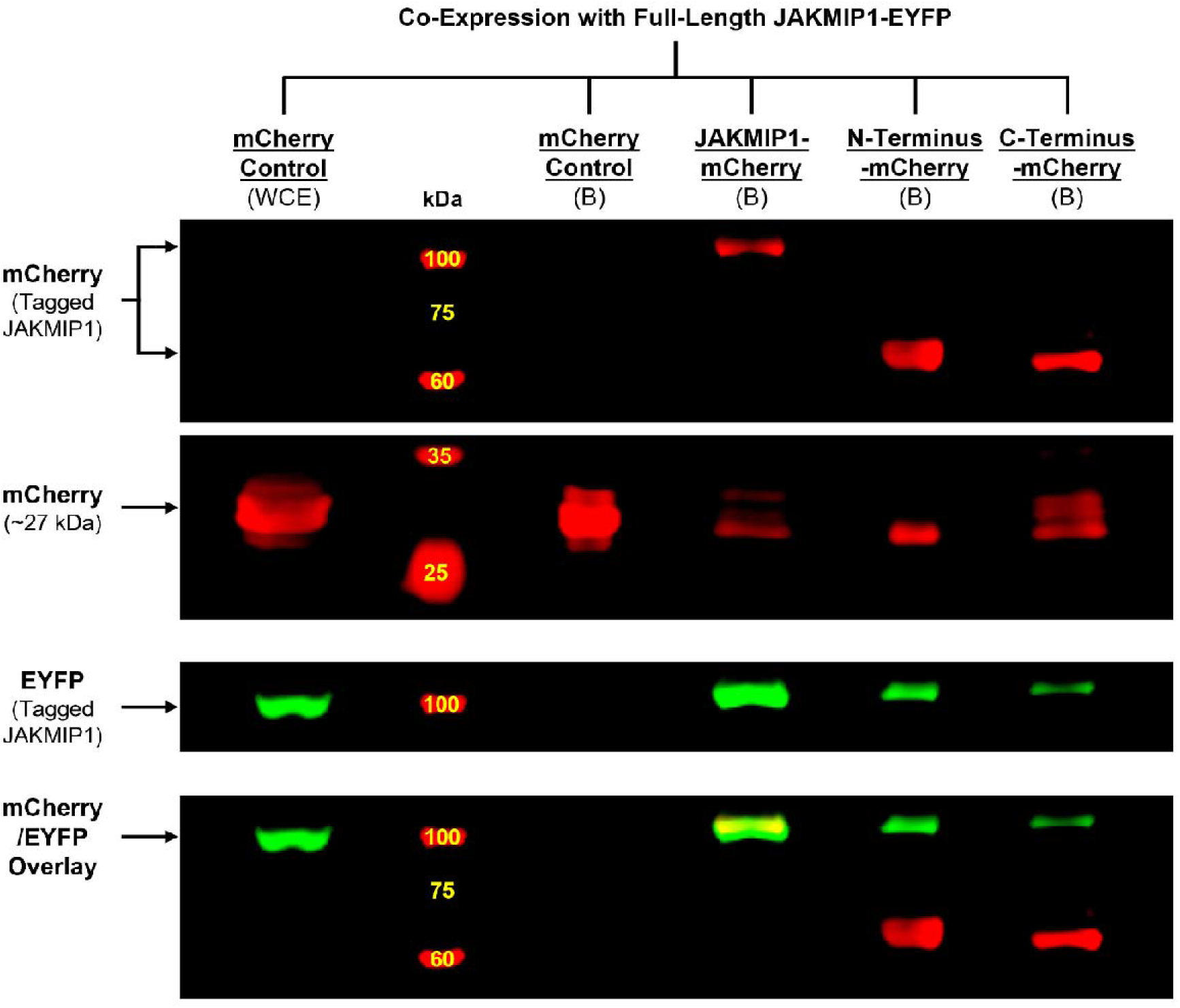
JAKMIP1 can dimerize at both its N- and C-termini. HEK-293 cells were transfected (using Lipofectamine™ LTX) with plasmid vectors encoding mCherry-tagged JAKMIP1 constructs in conjunction with an EYFP-tagged JAKMIP1 construct, allowed to express fusion proteins for 48 hours and then lysed for co-IP. JAKMIP1-EYFP/mCherry-OE SH-SY5Y cells were used as a control. Immunoprecipitation of mCherry-containing protein complexes was carried out using RFP-Trap® Agarose beads. Images representative of N = 2 independent samples generated from separate co-IP experiments; each sample subjected to Western blotting once. Lanes indicative of bound (“B”) fractions from cells expressing JAKMIP1-EYFP in conjunction with mCherry or JAKMIP1-mCherry constructs as indicated above each lane. The whole-cell extract (“WCE”) of JAKMIP1-EYFP/mCherry-OE HEK-293 cells was included as a positive control for protein-of-interest detection. Each fraction indicated was loaded into wells of polyacrylamide gels for SDS-PAGE. PVDF membranes were sequentially probed for EYFP and mCherry. JAKMIP1-EYFP is co-precipitated by the RFP-Trap® Agarose beads in HEK-293 cells expressing all the JAKMIP1-mCherry fusion constructs but not mCherry only. An overlay of the mCherry/EYFP signal is provided to demonstrate the similarity in molecular weight of the JAKMIP1-mCherry and JAKMIP1-EYFP fusion proteins. co-IP – co-immunoprecipitation; EYFP – enhanced yellow fluorescent protein; OE – overexpression; PVDF – polyvinylidene difluoride; SDS-PAGE – sodium dodecyl sulfate polyacrylamide gel electrophoresis.

**Supplementary Figure 17.**
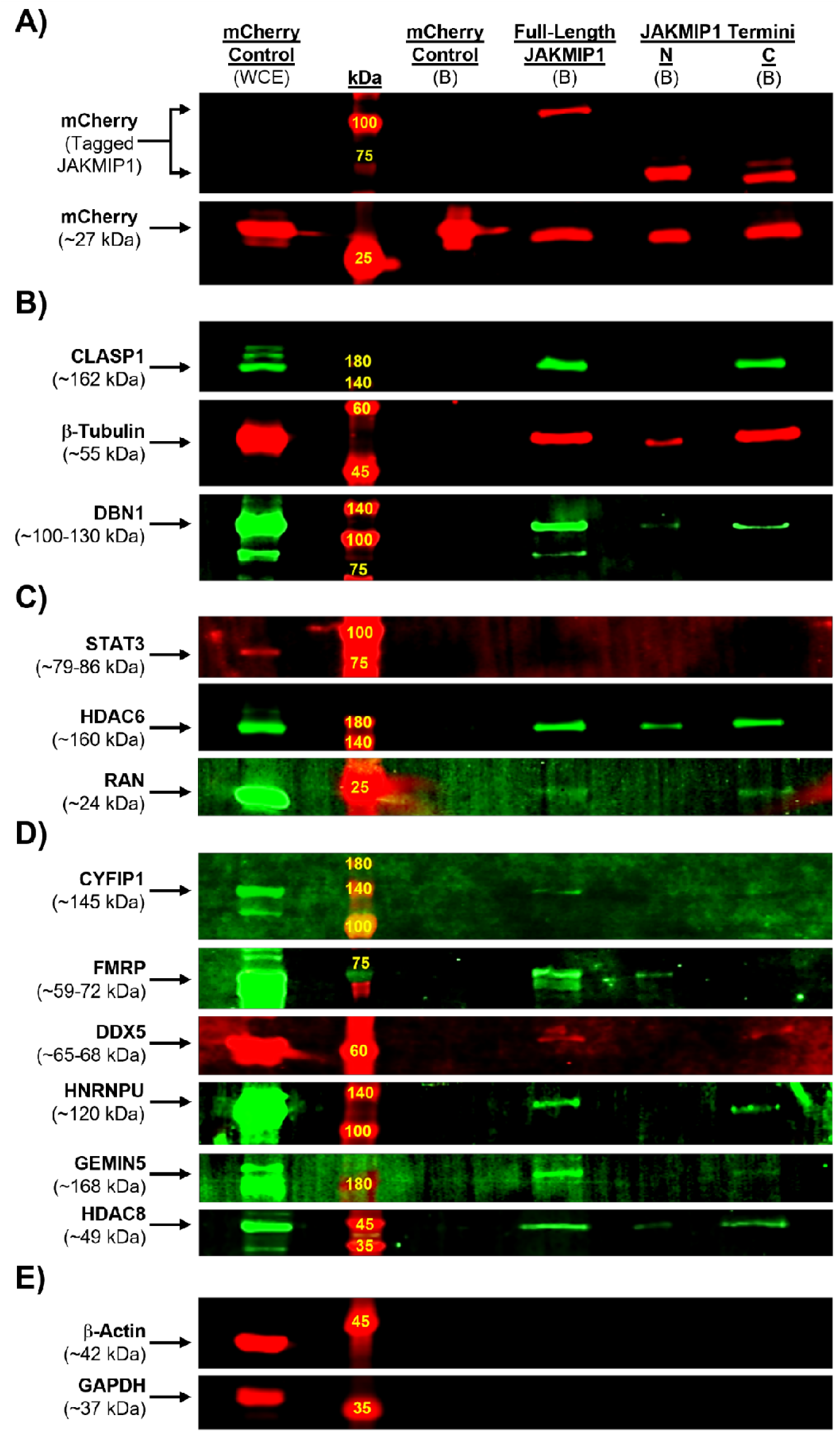
Confirmation of putative JAKMIP1 interactors by co-IP. Known and novel JAKMIP1-interacting proteins from IP-MS data were validated by co-IP. WT SH-SY5Y cells were transfected (using Lipofectamine™ LTX) with plasmid vectors encoding mCherry-tagged JAKMIP1 constructs and allowed to express fusion proteins for 48 hours prior to lysis and co-IP. mCherry-OE SH-SY5Y cells were used as a control. N = 2-3 independent samples generated from separate co-IP experiments; each sample subjected to Western blotting 1-2 times. Lanes indicative of bound (“B”) fractions from cells expressing each construct or mCherry only. The whole-cell extract (“WCE”) of mCherry-OE cells was included as a positive control for protein-of-interest detection. Proteins of interest targeted by primary antibodies are indicated to the left of the images. **A)** All JAKMIP1 constructs can be detected by mCherry at the appropriate molecular weight. **B)** All constructs interact with b-Tubulin and DBN1 but only the full-length and C-terminus pulls down CLASP1. **C)** No detectable interaction between the JAKMIP1 constructs and STAT3, but all constructs are capable of interacting with HDAC6, and RAN co-precipitates with the full-length or C-terminus constructs. **D)** CYFIP1 interacts with the full-length JAKMIP1 construct only, whereas FMRP interacts with the full-length and N-terminus constructs; and DDX5, HNRNPU and GEMIN5 interact with the full-length and C-terminus constructs. HDAC8 co-precipitates with all three constructs. **E)** No detectable contamination of the bound fractions by b-Actin or GAPDH, known non-JAKMIP1-interacting proteins. Co-IP – co-immunoprecipitation; IP-MS – immunoprecipitation mass spectrometry; OE – overexpression; SDS-PAGE – sodium dodecyl sulfate polyacrylamide gel electrophoresis; WT – wild-type.

**Supplementary Figure 18.**
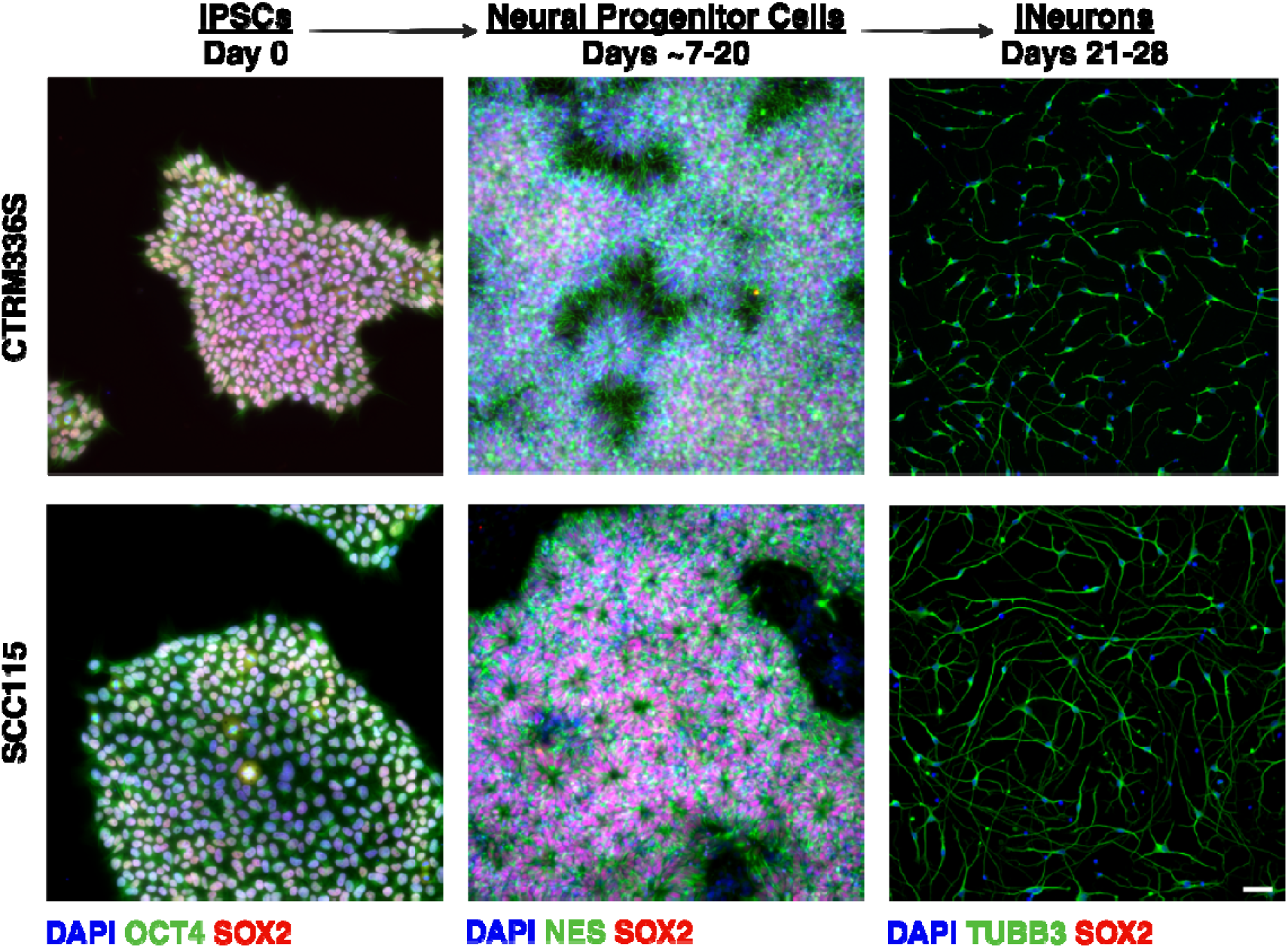
Confirmation of successful neuralization of CTRM336S and SCC115 hiPSCs into iNeurons. Representative fluorescence micrographs of CTRM336S and SCC115 cells fixed and stained for various markers for the neuralization process: prior to neuralization as hiPSCs (OCT4 (green) and SOX2 (red)), neural progenitor cells (NES (green) and SOX2 (red)) and iNeurons (TUBB3). hiPSC – human induced pluripotent stem cell; iNeuron – hiPSC-derived neuron. Scale bar = 100 μm.

**Supplementary Figure 19.**
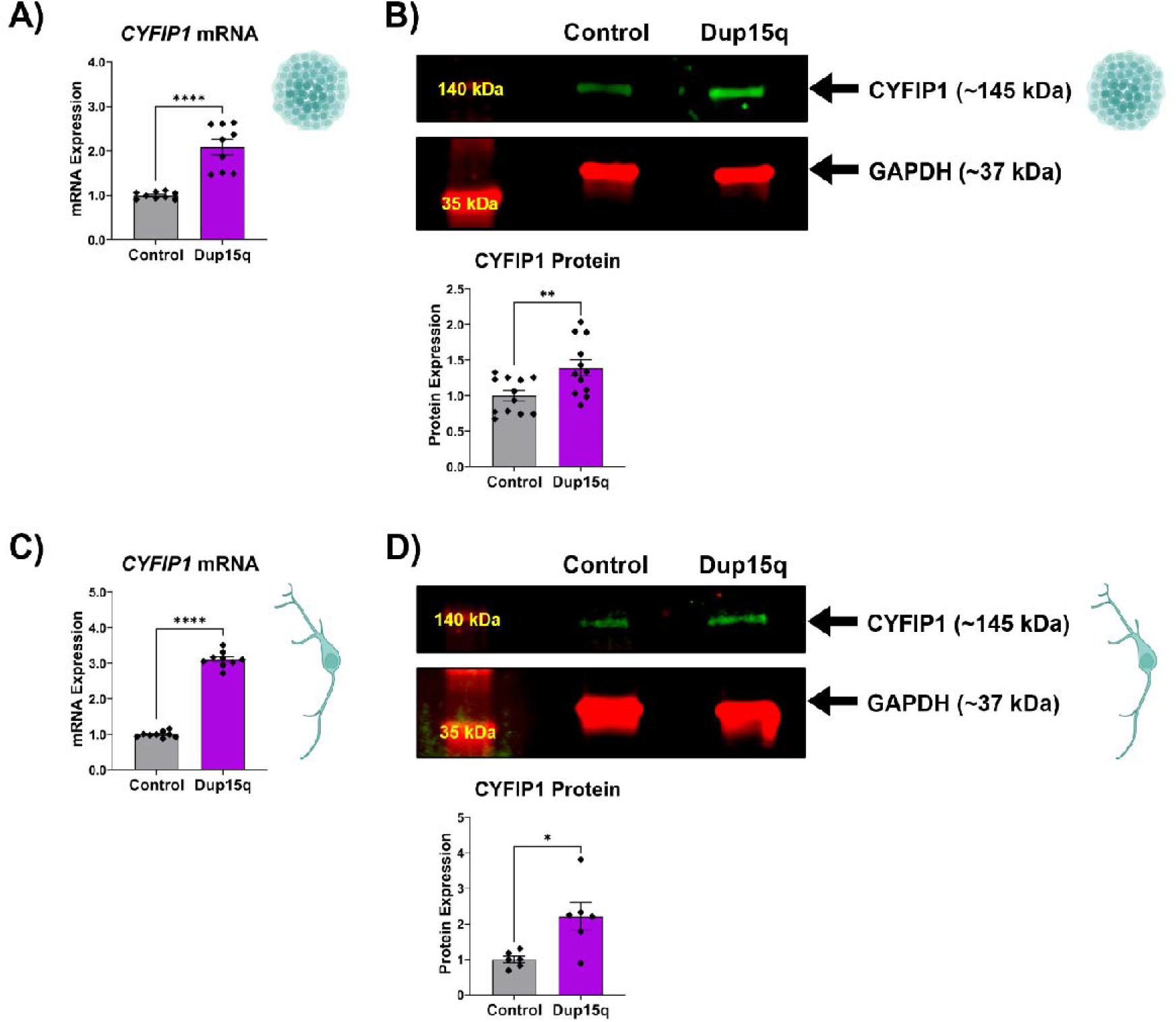
*CYFIP1* expression is raised in Dup15q hiPSCs and iNeurons. Relative mRNA and protein expression of *CYFIP1* in control and Dup15q **A-B)** hiPSCs and **C-D)** iNeurons. **A)** *CYFIP1* mRNA expression in control and Dup15q hiPSCs measured by qRT-PCR. Values presented as mean ± SEM of N = 3 independent experiments, 2-3 replicates each. One-way ANOVA with Tukey’s HSD test for multiple comparisons. Statistical significance displayed against the control; *P < 0.05; **P < 0.01; ****P < 0.0001. **B)** CYFIP1 protein expression in control and Dup15q hiPSCs measured by Western blotting. Values presented as mean ± SEM of N = 4 independent experiments, 3 replicates each. One-way ANOVA with Tukey’s HSD test for multiple comparisons. Statistical significance displayed as in A). **C)** *CYFIP1* mRNA expression in control and Dup15q iNeurons measured by qRT-PCR. Values presented as mean ± SEM of N = 3 independent experiments, 2-3 replicates each. One-way ANOVA with Tukey’s HSD test for multiple comparisons. Statistical significance displayed as in A). **D)** CYFIP1 protein expression in control and Dup15q iNeurons measured by Western blotting. Values presented as mean ± SEM of N = 3 independent experiments, 2 replicates each. One-way ANOVA with Tukey’s HSD test for multiple comparisons. Statistical significance displayed as in A). ANOVA – analysis of variance; Dup15q – chromosome 15q-duplication syndrome; mRNA – messenger RNA; qRT-PCR – quantitative reverse transcription PCR; SEM – standard error of the mean.

**Supplementary Figure 20.**
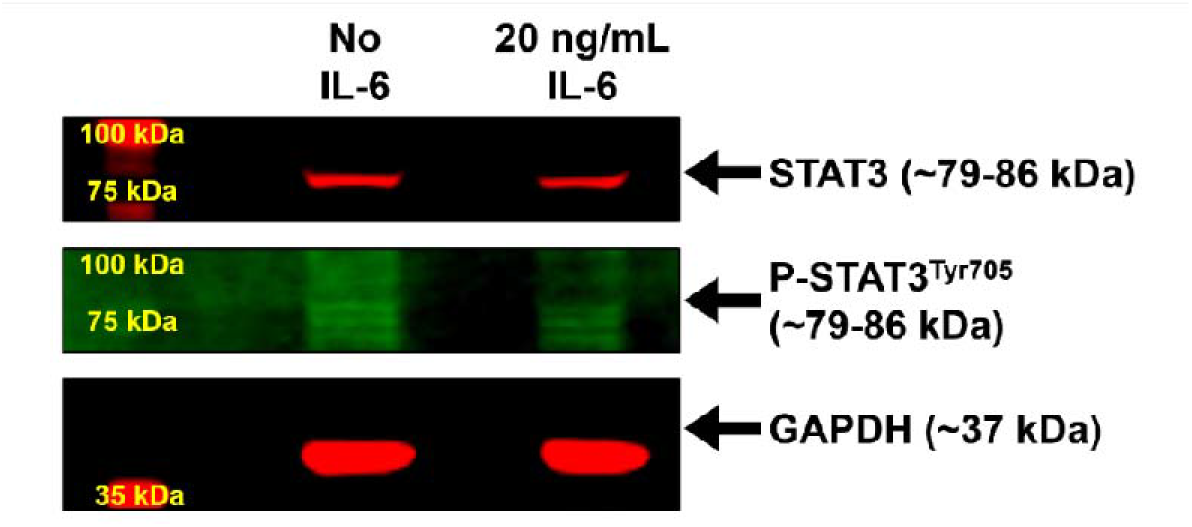
IL-6 does not induce STAT3 (Tyr^705^) phosphorylation in iNeurons. CTRM336S iNeurons were treated with 20 ng/mL IL-6 or B-27™ medium for 30 minutes prior to lysis for protein extraction. Western blotting was performed to measure STAT3 (Tyr^705^) phosphorylation relative to the GAPDH loading control. Note that no clear band for P-STAT3^Tyr705^ is visible (the brightness has been raised to the point of revealing background autofluorescence). Tyr^705^ – tyrosine residue at amino acid position 705.

**Supplementary Figure 21.**
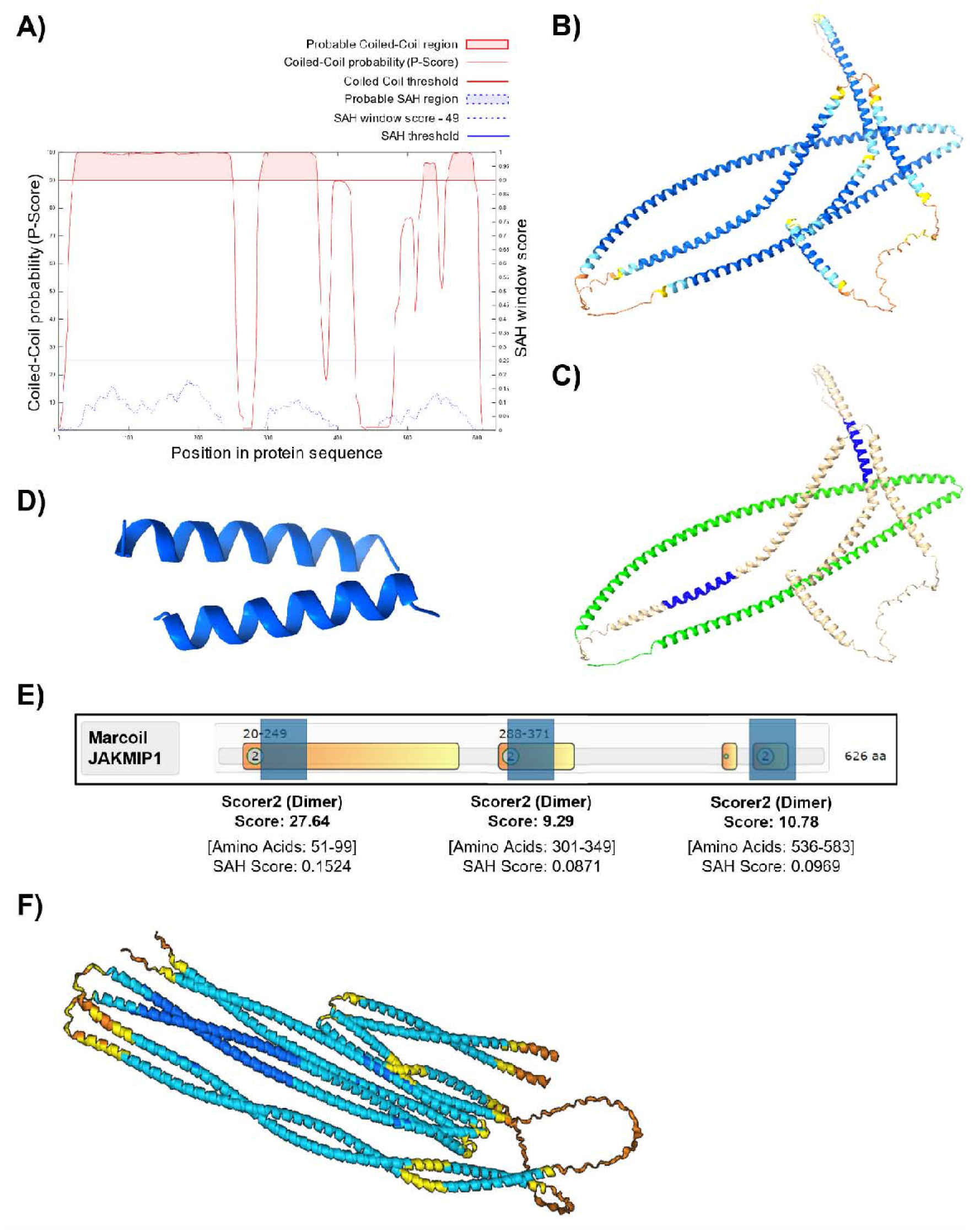
JAKMIP1 is composed of highly helical structures forming coiled coils. **A)** *Marcoil*-based prediction of coiled coils in the canonical JAKMIP1 isoform performed through *Waggawagga* (available at: https://waggawagga.motorprotein.de/). Results from *Marcoil* suggest JAKMIP1 is likely to have three main groups of coiled coils (red shaded areas in upper region of the graph) located at its N-terminus (amino acids^20-249^), RNA-binding domain (amino acids^288-371^) and C-terminus (amino acids^523-541^ and amino acids^556-595^). **B)** Predicted three-dimensional model structure of JAKMIP1 obtained from *AlphaFold* (available at: https://alphafold.ebi.ac.uk/), which are largely composed of long helices. Confidence of the model is reflected by the color of the amino acid residues (blue = very high confidence; cyan = confident; yellow = low confidence; orange = very low confidence). **C)** The N-terminal coiled coil domain and leucine zipper motif of the JAKMIP1 structure from B), highlighted in green and blue respectively. **D)** Predicted model of the leucine zipper region highlighted in C) by *AlphaFold 3* (available at: https://www.alphafoldserver.com/). **E)** *Marcoil*-based prediction of JAKMIP1 dimerization at three predicted coiled coil regions from part A). High *Scorer2* scores indicates that JAKMIP1 may form dimers mediated by the coiled coils highlighted in orange-yellow. For reference, MYH2, a cytoskeletal protein with known coiled-coils and is established to dimerize, has *Scorer2* scores between 6.83-18.55. SAH scores for the regions highlighted in blue are also provided. **F)** Predicted model of a JAKMIP1-JAKMIP1 dimer by *AlphaFold 3*, with colors of amino acid residues meaning the same as in B). The predicted structure demonstrates a well-fitting JAKMIP1 dimer with an even more coiled structure than singular JAKMIP1, reinforcing dimerization and the presence of functional coiled coils. SAH – single alpha helix.

## Supplementary Tables (File S2)

Supplementary tables are contained in File S2, which can be accessed online in the *figshare* repository at the following DOI: 10.6084/m9.figshare.30402832.

## Author Contributions (CRediT)

J.G.M – Conceptualization, Methodology, Validation, Formal Analysis, Investigation, Writing – Original Draft, Review & Editing, Visualization, Funding Acquisition

E-R.M – Conceptualization, Methodology, Validation, Formal Analysis, Investigation, Writing – Original Draft, Review & Editing, Visualization

N.T – Methodology, Investigation

C.E.H – Methodology, Resources

R.A.B – Methodology, Validation

Z.K.K.M – Data Curation

R.G.S –Supervision

N.G.M – Resources, Writing – Review & Editing

S.I-I – Investigation, Visualization

J.M – Resources

D.P.S – Resources

H.R.D – Conceptualization, Supervision

J.K.C – Conceptualization, Methodology, Resources

M.A.R – Conceptualization, Methodology, Writing – Review & Editing, Supervision

A.O-A – Conceptualization, Supervision, Writing – Review & Editing, Funding Acquisition

## Data Availability Statement

The datasets used and/or analyzed during the current study are available from the corresponding author on reasonable request.

## Funding

Wellcome Trust funded Translational Research Exchange @ Exeter (TREE) pump-priming fund (Project 121528) (J.G.M) and the Bristol Japanese Cultural Showcase (A.O-A).

## Ethics Approval and Consent to Participate

Not applicable.

## Declaration of competing interests

The authors declare that they have no known competing financial interests or personal relationships that could have appeared to influence the work reported in this paper.

## Acknowledgements

The research was carried out at the National Institute for Health and Care Research (NIHR) Exeter Biomedical Research Centre (BRC). We gratefully acknowledge the RILD Imaging Facility, University of Exeter, as well as the RILD Technical Team for their support and assistance with this work. We also extend our sincere gratitude to the generous support of the charitable organizations that contributed to this work, including the Wellcome Trust Translational Research Exchange @ Exeter (TREE) pump-priming fund and the Bristol Japanese Cultural Showcase. We would like to thank Dr. Kate Heesom at the University of Bristol Proteomics Facility for performing the LC-MS and Dr. Lauren Adams for her support, advice and guidance with the planning of the IP-MS and co-IP experiments. We would also extend our deep appreciation to Dr. Roland Naggy for his guidance with iNeuron differentiation, and Liming Yang for his assistance with iPSC culture. Finally, we would like to acknowledge the insightful discussions with Prof. Lorna Harries, Prof. Antony Vernon, Dr. Emma Dempster, Dr. Craig Beall, Dr. Nicholas Clifton, and Dr. Kaiyven Afi Leslie.

## Notes

https://doi.org/10.6084/m9.figshare.30402832.v1

### Competing Interest Statement

The authors have declared no competing interest.

### Summary of Updates

Discussion section updated to further discuss limitations of the study

## References

1. Abrahams, B.S. et al. (2013) “SFARI Gene 2.0: A community-driven knowledgebase for the autism spectrum disorders (ASDs),” Molecular Autism, 4(1), p. 36. Available at: 10.1186/2040-2392-4-36.

2. Andrews, S. (2010) “FastQC: A Quality Control Tool for High Throughput Sequence Data.” Available at: http://www.bioinformatics.babraham.ac.uk/projects/fastqc/.

3. Battle, D.J. et al. (2006) “The Gemin5 Protein of the SMN Complex Identifies snRNAs,” Molecular Cell, 23(2), pp. 273–279. Available at: 10.1016/J.MOLCEL.2006.05.036.

4. Van Battum, E.Y., Brignani, S. and Pasterkamp, R.J. (2015) “Axon guidance proteins in neurological disorders,” The Lancet Neurology, 14(5), pp. 532–546. Available at: 10.1016/S1474-4422(14)70257-1.

5. Bella, A.J. et al. (2006) “Brain-derived neurotrophic factor (BDNF) acts primarily via the JAK/STAT pathway to promote neurite growth in the major pelvic ganglion of the rat: Part I,” Journal of Sexual Medicine, 3(5), pp. 815–820. Available at: 10.1111/j.1743-6109.2006.00291.x.

6. Berg, J.M. et al. (2015) “JAKMIP1, a Novel Regulator of Neuronal Translation, Modulates Synaptic Function and Autistic-like Behaviors in Mouse,” Neuron, 88(6), pp. 1173–1191. Available at: 10.1016/j.neuron.2015.10.031.

7. Bourgeron, T. (2015) “From the genetic architecture to synaptic plasticity in autism spectrum disorder,” Nature Reviews Neuroscience 2015 16:9, 16(9), pp. 551–563. Available at: 10.1038/nrn3992.

8. Cauvi, D.M. et al. (2017) “From the Cover: Interplay Between IFN-γ and IL-6 Impacts the Inflammatory Response and Expression of Interferon-Regulated Genes in Environmental-Induced Autoimmunity,” Toxicological Sciences, 158(1), pp. 227–239. Available at: 10.1093/TOXSCI/KFX083.

9. Cervera-Juanes, R. et al. (2017) “Alcohol-dose-dependent DNA methylation and expression in the nucleus accumbens identifies coordinated regulation of synaptic genes.,” Translational Psychiatry, 7(1), pp. e994–e994. Available at: 10.1038/TP.2016.266.

10. Chen, H. et al. (2016) “IL-10 Promotes Neurite Outgrowth and Synapse Formation in Cultured Cortical Neurons after the Oxygen-Glucose Deprivation via JAK1/STAT3 Pathway,” Scientific Reports 2016 6:1, 6(1), pp. 1–16. Available at: 10.1038/srep30459.

11. Cieślik, M. et al. (2020) “Maternal Immune Activation Induces Neuroinflammation and Cortical Synaptic Deficits in the Adolescent Rat Offspring,” International Journal of Molecular Sciences 2020, Vol. 21, Page 4097, 21(11), p. 4097. Available at: 10.3390/IJMS21114097.

12. Costa, V. et al. (2007) “Identification and expression analysis of novel Jakmip1 transcripts,” Gene, 402(1–2), pp. 1–8. Available at: 10.1016/j.gene.2007.07.001.

13. Couch, A.C.M. et al. (2023) “Acute IL-6 exposure triggers canonical IL6Ra signaling in hiPSC microglia, but not neural progenitor cells,” Brain, Behavior, and Immunity, 110, p. 43. Available at: 10.1016/J.BBI.2023.02.007.

14. Cunningham, F. et al. (2022) “Ensembl 2022,” Nucleic Acids Research, 50(D1), pp. D988–D995. Available at: 10.1093/NAR/GKAB1049.

15. Dardenne, E. et al. (2014) “RNA Helicases DDX5 and DDX17 Dynamically Orchestrate Transcription, miRNA, and Splicing Programs in Cell Differentiation,” Cell Reports, 7(6), pp. 1900–1913. Available at: 10.1016/J.CELREP.2014.05.010.

16. Dobin, A. et al. (2013) “STAR: ultrafast universal RNA-seq aligner,” Bioinformatics (Oxford, England), 29(1), pp. 15–21. Available at: 10.1093/BIOINFORMATICS/BTS635.

17. Donnard, E., Shu, H. and Garber, M. (2022) “Single cell transcriptomics reveals dysregulated cellular and molecular networks in a fragile X syndrome model,” PLoS Genetics, 18(6). Available at: 10.1371/JOURNAL.PGEN.1010221.

18. Esquivel-Rendón, E. et al. (2019) “Interleukin 6 Dependent Synaptic Plasticity in a Social Defeat-Susceptible Prefrontal Cortex Circuit,” Neuroscience, 414, pp. 280–296. Available at: 10.1016/J.NEUROSCIENCE.2019.07.002.

19. Ewels, P. et al. (2016) “MultiQC: summarize analysis results for multiple tools and samples in a single report,” Bioinformatics (Oxford, England), 32(19), pp. 3047–3048. Available at: 10.1093/BIOINFORMATICS/BTW354.

20. Gandawijaya, J. et al. (2021) “Cell Adhesion Molecules Involved in Neurodevelopmental Pathways Implicated in 3p-Deletion Syndrome and Autism Spectrum Disorder,” Frontiers in Cellular Neuroscience, 14, p. 466. Available at: 10.3389/fncel.2020.611379.

21. Guzman-Martinez, L. et al. (2019) “Neuroinflammation as a common feature of neurodegenerative disorders,” Frontiers in Pharmacology, 10(SEP), p. 452270. Available at: 10.3389/FPHAR.2019.01008/BIBTEX.

22. Han, B. yue et al. (2022) “HNRNPU promotes the progression of triple-negative breast cancer via RNA transcription and alternative splicing mechanisms,” Cell Death & Disease 2022 13:11, 13(11), pp. 1–15. Available at: 10.1038/s41419-022-05376-6.

23. Han, V.X. et al. (2021) “Maternal immune activation and neuroinflammation in human neurodevelopmental disorders,” Nature Reviews Neurology 2021 17:9, 17(9), pp. 564–579. Available at: 10.1038/s41582-021-00530-8.

24. Harlow, C.E. et al. (2022) “Identification and single-base gene-editing functional validation of a cis-EPO variant as a genetic predictor for EPO-increasing therapies,” American journal of human genetics, 109(9), pp. 1638–1652. Available at: 10.1016/J.AJHG.2022.08.004.

25. Hedges, D.J. et al. (2012) “Evidence of novel fine-scale structural variation at autism spectrum disorder candidate loci,” Molecular Autism, 3(1), p. 2. Available at: 10.1186/2040-2392-3-2.

26. Helbig, I. et al. (2020) “Whole-exome and HLA sequencing in Febrile infection-related epilepsy syndrome.,” Annals of Clinical and Translational Neurology, 7(8), pp. 1429–1435. Available at: 10.1002/ACN3.51062.

27. Hong, G. et al. (2014) “Separate enrichment analysis of pathways for up- and downregulated genes,” Journal of the Royal Society Interface, 11(92). Available at: 10.1098/RSIF.2013.0950.

28. Hu, E. et al. (2000) “Cloning and characterization of a novel human class I histone deacetylase that functions as a transcription repressor,” The Journal of biological chemistry, 275(20), pp. 15254–15264. Available at: 10.1074/JBC.M908988199.

29. Ihara, S. et al. (1996) “Identification of Interleukin-6 as a Factor That Induces Neurite Outgrowth by PC12 Cells Primed with NGF,” Journal of Biochemistry, 120(5), pp. 865–868. Available at: 10.1093/oxfordjournals.jbchem.a021499.

30. Islam, O., et al. (2009) “Interleukin-6 and Neural Stem Cells: More Than Gliogenesis,” Molecular Biology of the Cell. Edited by C.-H. Heldin, 20(1), pp. 188–199. Available at: 10.1091/mbc.e08-05-0463.

31. Kent, W.J. et al. (2002) “The human genome browser at UCSC.,” Genome research, 12(6), pp. 996–1006. Available at: 10.1101/gr.229102.

32. de la Torre-Ubieta, L., et al. (2016) “Advancing the understanding of autism disease mechanisms through genetics,” Nature Medicine, 22(4), pp. 345–361. Available at: 10.1038/nm.4071.

33. Li, H. et al. (2009) “The Sequence Alignment/Map format and SAMtools,” Bioinformatics, 25(16), pp. 2078–2079. Available at: 10.1093/BIOINFORMATICS/BTP352.

34. Li, Q.S., Sun, Y. and Wang, T. (2020) “Epigenome-wide association study of Alzheimer’s disease replicates 22 differentially methylated positions and 30 differentially methylated regions.,” Clinical Epigenetics, 12(1), pp. 149–149. Available at: 10.1186/S13148-020-00944-Z.

35. Liao, Y., Smyth, G.K. and Shi, W. (2014) “featureCounts: an efficient general purpose program for assigning sequence reads to genomic features,” Bioinformatics (Oxford, England), 30(7), pp. 923–930. Available at: 10.1093/BIOINFORMATICS/BTT656.

36. Lin, M. and Guo, J.T. (2019) “New insights into protein–DNA binding specificity from hydrogen bond based comparative study,” Nucleic Acids Research, 47(21), pp. 11103–11113. Available at: 10.1093/NAR/GKZ963.

37. Love, M.I., Huber, W. and Anders, S. (2014) “Moderated estimation of fold change and dispersion for RNA-seq data with DESeq2,” Genome Biology, 15(12), pp. 1–21. Available at: 10.1186/S13059-014-0550-8/FIGURES/9.

38. Martin, E.-R. et al. (2026) “CYFIP1 overexpression amplifies IL-6/STAT3 and IFN-γ/STAT1 signaling: potential implications for neuroinflammation and autism spectrum disorder,” Brain, Behavior, & Immunity - Health, 55, p. 101293. Available at: 10.1016/J.BBIH.2026.101293.

39. Martin, E.-R., Gandawijaya, J. and Oguro-Ando, A. (2022) “A novel method for generating glutamatergic SH-SY5Y neuron-like cells utilizing B-27 supplement,” Frontiers in Pharmacology, 13, p. 943627. Available at: 10.3389/FPHAR.2022.943627.

40. Mill, J. et al. (2008) “Epigenomic profiling reveals DNA-methylation changes associated with major psychosis.,” American Journal of Human Genetics, 82(3), pp. 696–711. Available at: 10.1016/J.AJHG.2008.01.008.

41. Narimatsu, M. et al. (2001) “Tissue-Specific Autoregulation of the stat3 Gene and Its Role in Interleukin-6-Induced Survival Signals in T Cells,” Molecular and Cellular Biology, 21(19), p. 6615. Available at: 10.1128/MCB.21.19.6615-6625.2001.

42. Nebel, R.A., et al. (2016) “Reduced CYFIP1 in Human Neural Progenitors Results in Dysregulation of Schizophrenia and Epilepsy Gene Networks,” PLOS ONE. Edited by C. Walss-Bass, 11(1), p. e0148039. Available at: 10.1371/journal.pone.0148039.

43. Nishimura, Y. et al. (2007) “Genome-wide expression profiling of lymphoblastoid cell lines distinguishes different forms of autism and reveals shared pathways,” Human Molecular Genetics, 16(14), pp. 1682–1698. Available at: 10.1093/hmg/ddm116.

44. Parker-Athill, E.C. and Tan, J. (2011) “Maternal immune activation and autism spectrum disorder: Interleukin-6 signaling as a key mechanistic pathway,” NeuroSignals, 18(2), pp. 113–128. Available at: 10.1159/000319828.

45. Pfaffl, M.W. (2001) “A new mathematical model for relative quantification in real-time RT–PCR,” Nucleic Acids Research, 29(9), p. e45. Available at: 10.1093/NAR/29.9.E45.

46. R Core Team (2020) “R: A Language and Environment for Statistical Computing.” Vienna, Austria: R Foundation for Statistical Computing. Available at: https://www.R-project.org (Accessed: March 2, 2023).

47. Raudvere, U. et al. (2019) “g:Profiler: a web server for functional enrichment analysis and conversions of gene lists (2019 update),” Nucleic acids research, 47(W1), pp. W191–W198. Available at: 10.1093/NAR/GKZ369.

48. Recasens, M. et al. (2021) “Chronic exposure to IL-6 induces a desensitized phenotype of the microglia,” Journal of Neuroinflammation, 18(1), pp. 1–22. Available at: 10.1186/S12974-020-02063-1/TABLES/3.

49. RStudio Team (2020) “RStudio: Integrated Development for R.” Boston, MA: RStudio, PBC. Available at: https://www.rstudio.com/ (Accessed: November 22, 2022).

50. Russell, M.A. et al. (2013) “Differential effects of interleukin-13 and interleukin-6 on Jak/STAT signaling and cell viability in pancreatic β-cells,” Islets, 5(2), p. 95. Available at: 10.4161/ISL.24249.

51. Sapir, T. and Reiner, O. (2023) “HNRNPU’s multi-tasking is essential for proper cortical development,” BioEssays : news and reviews in molecular, cellular and developmental biology, 45(9). Available at: 10.1002/BIES.202300039.

52. Sarieva, K. et al. (2023) “Human brain organoid model of maternal immune activation identifies radial glia cells as selectively vulnerable,” Molecular Psychiatry 2023, pp. 1–13. Available at: 10.1038/s41380-023-01997-1.

53. Sarkar, M., Khare, V. and Ghosh, M.K. (2016) “The DEAD box protein p68: a novel coactivator of Stat3 in mediating oncogenesis,” Oncogene 2017 36:22, 36(22), pp. 3080–3093. Available at: 10.1038/onc.2016.449.

54. Schäfer, K.H. et al. (1999) “The IL-6/sIL-6R fusion protein hyper-IL-6 promotes neurite outgrowth and neuron survival in cultured enteric neurons,” Journal of Interferon and Cytokine Research, 19(5), pp. 527–532. Available at: 10.1089/107999099313974.

55. Schindelin, J. et al. (2012) “Fiji: an open-source platform for biological-image analysis,” Nature Methods, 9(7), pp. 676–682. Available at: 10.1038/nmeth.2019.

56. Schneider, V.A. et al. (2017) “Evaluation of GRCh38 and de novo haploid genome assemblies demonstrates the enduring quality of the reference assembly,” Genome Research, 27(5), pp. 849–864. Available at: 10.1101/GR.213611.116.

57. Seldeen, K.L. et al. (2010) “Dissecting the Role of Leucine Zippers in the Binding of bZIP Domains of Jun Transcription Factor to DNA,” Biochemical and biophysical research communications, 394(4), p. 1030. Available at: 10.1016/J.BBRC.2010.03.116.

58. Sims, N.A. (2020) “The JAK1/STAT3/SOCS3 axis in bone development, physiology, and pathology,” Experimental & Molecular Medicine 2020 52:8, 52(8), pp. 1185–1197. Available at: 10.1038/s12276-020-0445-6.

59. Steindler, C. et al. (2004) “Jamip1 (Marlin-1) Defines a Family of Proteins Interacting with Janus Kinases and Microtubules,” Journal of Biological Chemistry, 279(41), pp. 43168–43177. Available at: 10.1074/jbc.M401915200.

60. Sun, X. and Kaufman, P.D. (2018) “Ki-67: more than a proliferation marker,” Chromosoma, 127(2), p. 175. Available at: 10.1007/S00412-018-0659-8.

61. Szklarczyk, D. et al. (2019) “STRING v11: protein-protein association networks with increased coverage, supporting functional discovery in genome-wide experimental datasets,” Nucleic acids research, 47(D1), pp. D607–D613. Available at: 10.1093/NAR/GKY1131.

62. Tanaka, T., Narazaki, M. and Kishimoto, T. (2014) “IL-6 in Inflammation, Immunity, and Disease,” Cold Spring Harbor Perspectives in Biology, 6(10), pp. 16295–16296. Available at: 10.1101/CSHPERSPECT.A016295.

63. Tang, X. et al. (2020) “HDAC8 cooperates with SMAD3/4 complex to suppress SIRT7 and promote cell survival and migration,” Nucleic Acids Research, 48(6), pp. 2912–2923. Available at: 10.1093/NAR/GKAA039.

64. Tellier, M., Maudlin, I. and Murphy, S. (2020) “Transcription and splicing: A two-way street,” Wiley interdisciplinary reviews. RNA, 11(5). Available at: 10.1002/WRNA.1593.

65. Untergasser, A. et al. (2012) “Primer3—new capabilities and interfaces,” Nucleic Acids Research, 40(15), p. e115. Available at: 10.1093/NAR/GKS596.

66. Vargas, D.L. et al. (2005) “Neuroglial activation and neuroinflammation in the brain of patients with autism,” Annals of Neurology, 57(1), pp. 67–81. Available at: 10.1002/ana.20315.

67. Vidal, R.L. et al. (2007) “Marlin-1 and conventional kinesin link GABAB receptors to the cytoskeleton and regulate receptor transport,” Molecular and Cellular Neuroscience, 35(3), pp. 501–512. Available at: 10.1016/j.mcn.2007.04.008.

68. Vidal, R.L. et al. (2009) “Cellular and subcellular localization of Marlin-1 in the brain,” BMC Neuroscience, 10, p. 37. Available at: 10.1186/1471-2202-10-37.

69. Vidal, R.L. et al. (2012) “RNA interference of Marlin-1/Jakmip1 results in abnormal morphogenesis and migration of cortical pyramidal neurons,” Molecular and Cellular Neuroscience, 51(1–2), pp. 1–11. Available at: 10.1016/j.mcn.2012.07.007.

70. Warre-Cornish, K. et al. (2020) “Interferon-γ signaling in human iPSC–derived neurons recapitulates neurodevelopmental disorder phenotypes,” Science Advances, 6(34). Available at: 10.1126/SCIADV.AAY9506.

71. Wei, H. et al. (2011) “IL-6 is increased in the cerebellum of autistic brain and alters neural cell adhesion, migration and synaptic formation,” Journal of Neuroinflammation, 8, p. 52. Available at: 10.1186/1742-2094-8-52.

72. Wu, W.L. et al. (2017) “The placental interleukin-6 signaling controls fetal brain development and behavior,” Brain, Behavior, and Immunity, 62, pp. 11–23. Available at: 10.1016/j.bbi.2016.11.007.

73. Wu, Y.Y. and Bradshaw, R.A. (1996) “Induction of neurite outgrowth by interleukin-6 is accompanied by activation of Stat3 signaling pathway in a variant PC12 cell (E2) line,” The Journal of biological chemistry, 271(22), pp. 13023–13032. Available at: 10.1074/JBC.271.22.13023.

74. Xiao, R. et al. (2012) “Nuclear matrix factor hnRNP U/SAF-A exerts a global control of alternative splicing by regulating U2 snRNP maturation,” Molecular cell, 45(5), pp. 656–668. Available at: 10.1016/J.MOLCEL.2012.01.009.

75. Yang, G. and Tang, W.Y. (2017) “Resistance of interleukin-6 to the extracellular inhibitory environment promotes axonal regeneration and functional recovery following spinal cord injury,” International Journal of Molecular Medicine, 39(2), pp. 437–445. Available at: 10.3892/IJMM.2017.2848/HTML.

76. Yang, W.X. et al. (2016) “Analyses of Genotypes and Phenotypes of Ten Chinese Patients with Wolf-Hirschhorn Syndrome by Multiplex Ligation-dependent Probe Amplification and Array Comparative Genomic Hybridization.,” Chinese Medical Journal, 129(6), pp. 672–678. Available at: 10.4103/0366-6999.177996.

77. Yehia, L. et al. (2020) “Copy Number Variation and Clinical Outcomes in Patients With Germline PTEN Mutations,” JAMA Network Open, 3(1), pp. e1920415–e1920415. Available at: 10.1001/JAMANETWORKOPEN.2019.20415.

78. Yong, J. et al. (2010) “Gemin5 delivers snRNA precursors to the SMN complex for snRNP biogenesis,” Molecular cell, 38(4), p. 551. Available at: 10.1016/J.MOLCEL.2010.03.014.

79. Zawadzka, A., Cieślik, M. and Adamczyk, A. (2021) “The Role of Maternal Immune Activation in the Pathogenesis of Autism: A Review of the Evidence, Proposed Mechanisms and Implications for Treatment,” International Journal of Molecular Sciences 2021, Vol. 22, Page 11516, 22(21), p. 11516. Available at: 10.3390/IJMS222111516.

80. Zouein, F.A., Duhé, R.J. and Booz, G.W. (2011) “JAKs go nuclear: Emerging role of nuclear JAK1 and JAK2 in gene expression and cell growth,” Growth Factors, 29(6), pp. 245–252. Available at: 10.3109/08977194.2011.614949.

