## Supplementary File 1 for "Exploring the functions of JAKMIP1 in neuronal IL-6/STAT3 signaling and its relevance to chromosome 15q-duplication syndrome"

Supplementary Figures (File S1)

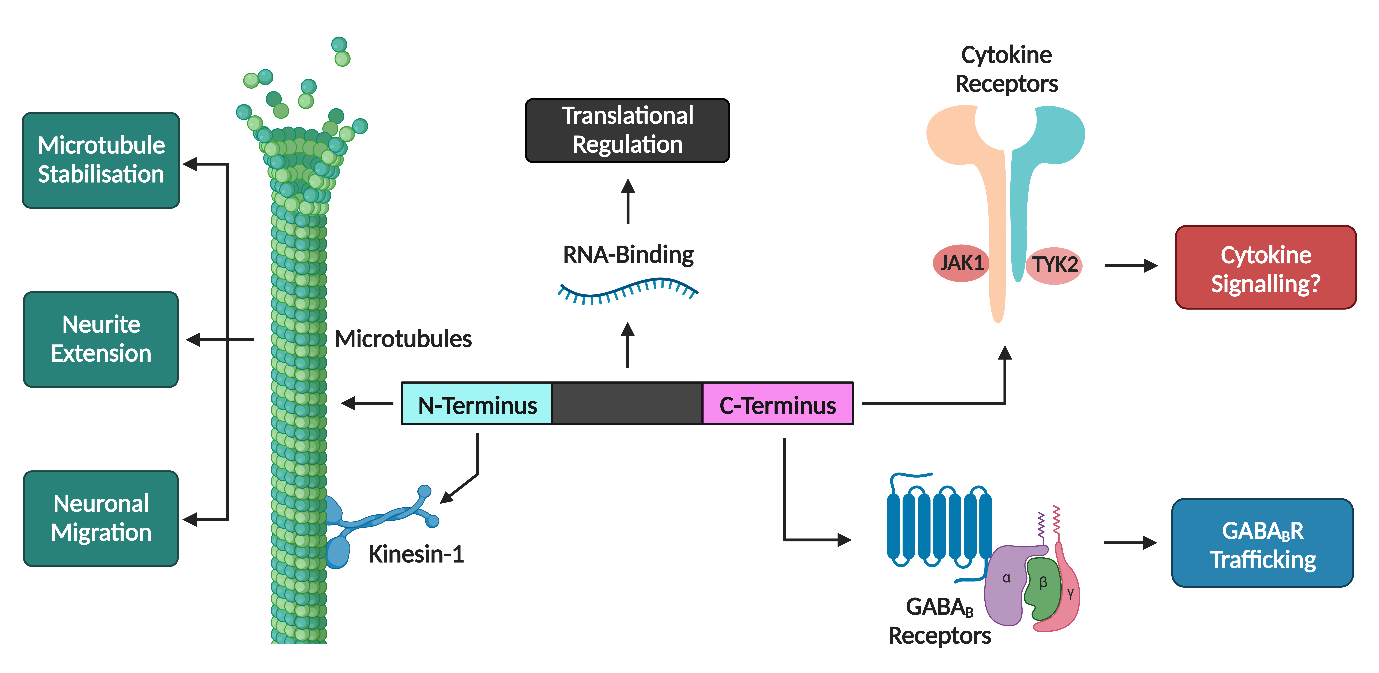

**Supplementary Figure 1 | Distinct functional domains of the JAKMIP1 protein.** Schematic overview of the known functional domains of JAKMIP1 as well as their interactions. The JAKMIP1 N-terminus associates with microtubules and motor proteins; a central region interacts with mRNAs and is involved in translational regulation; and the C-terminus interacts with JAKs and GABA_B_R1. Adapted from (*16*–*18*, *46*). Created with BioRender.com.

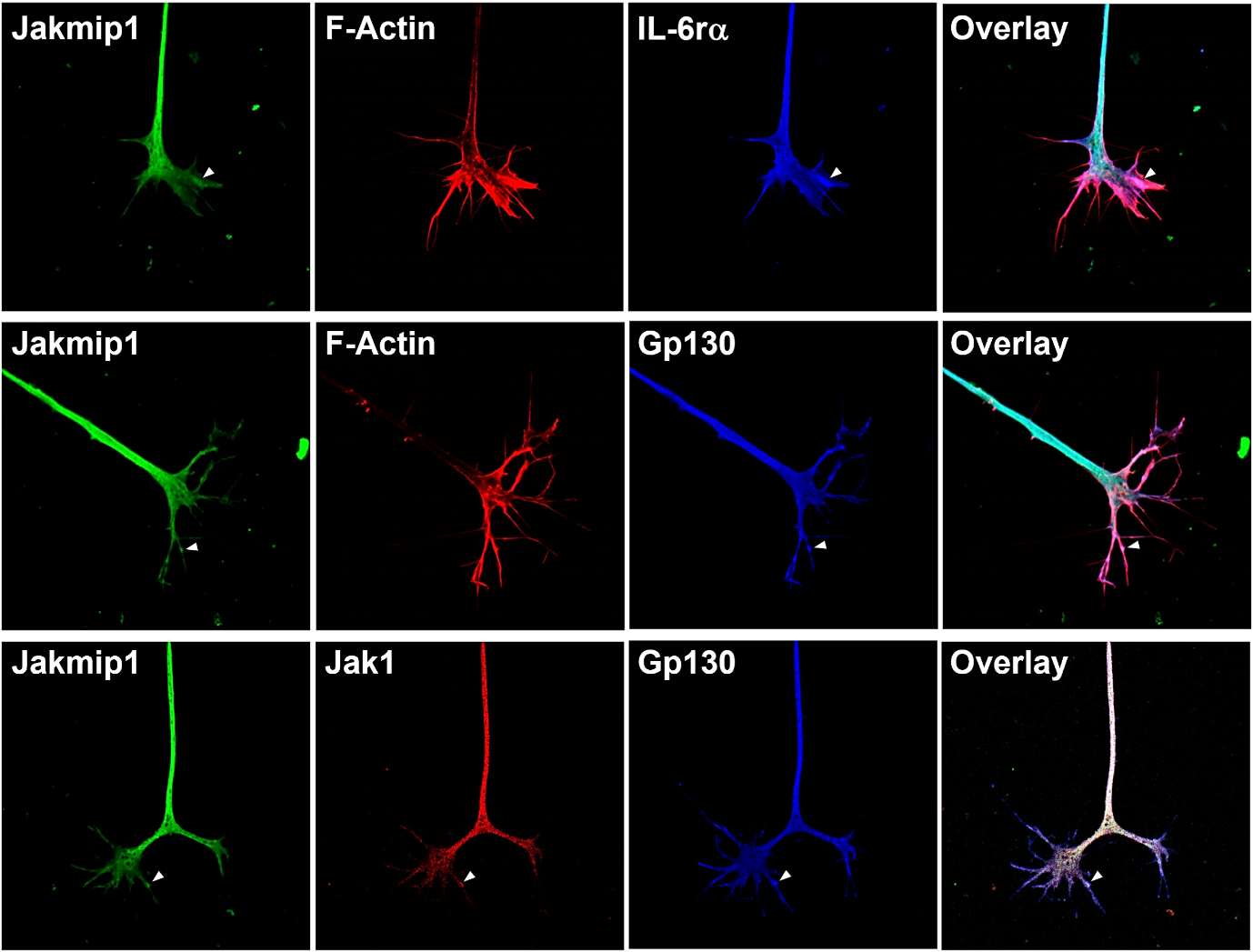

**Supplementary Figure 2 | Jakmip1 co-localizes with components of the IL-6r complex in chick neurons.** Chick dorsal root ganglion neurons were explanted at embryonic day 7 and cultured in 100 ng/mL IL-6, fixed, permeabilized and stained for varying combinations of Jakmip1, IL-6rα, Gp130, Jak1 and F-Actin. Arrowheads indicate examples of co-localization between Jakmip1 and IL-6rα, Gp130 or Jak1.

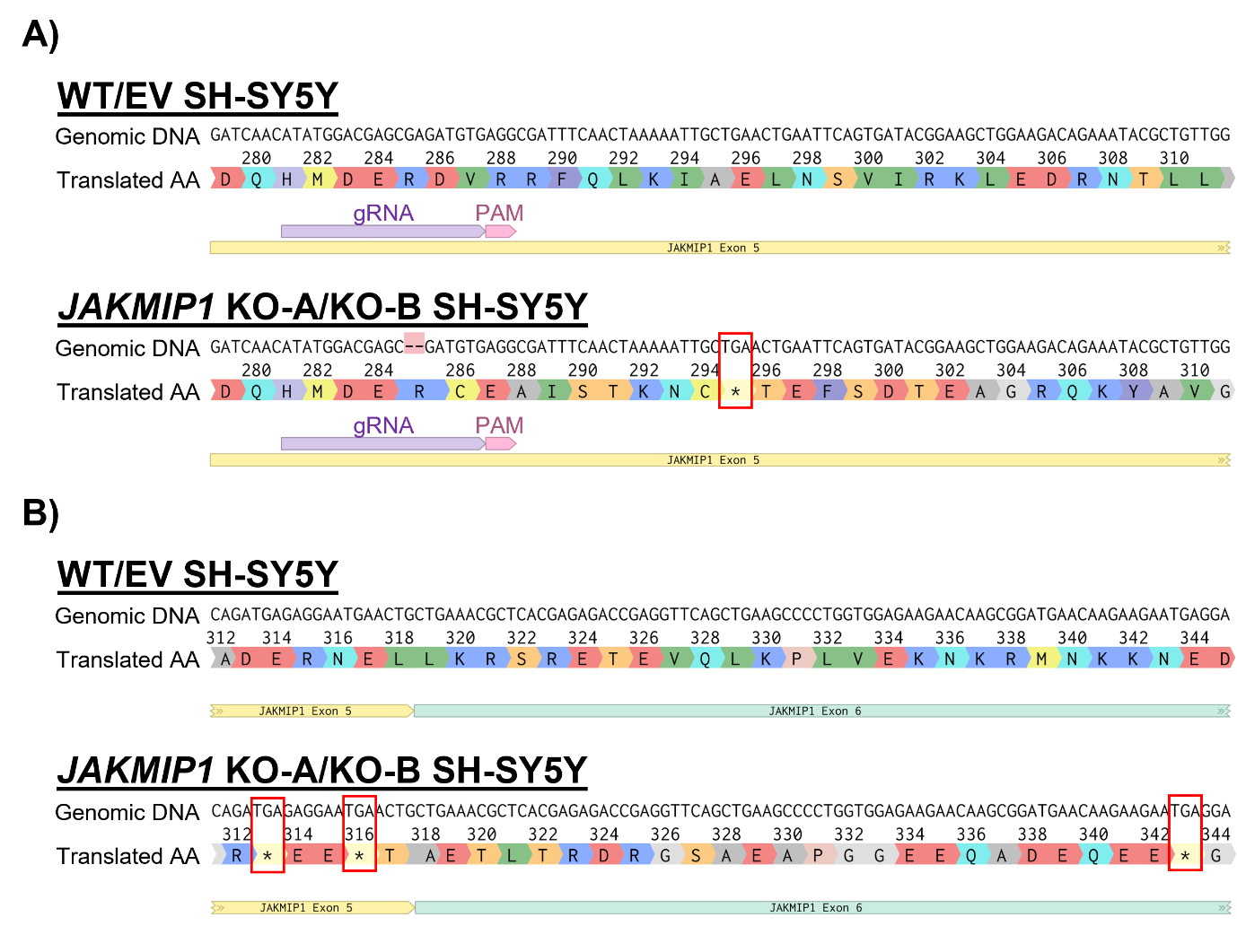

**Supplementary Figure 3 | Sanger sequencing reveals 2-bp deletion in *JAKMIP1*^-/-^ SH-SY5Y cells. A)** Visualization of Sanger sequencing results from *GENEWIZ*, showing the genomic DNA sequence of WT, EV control and both *JAKMIP1*^-/-^ SH-SY5Y cell lines (named KO-A and KO-B) and the subsequent translated amino acid sequence. A 2-bp deletion is identified (mismatch in DNA alignment shown in red) within the gRNA sequence (in purple) in exon 5 of the *JAKMIP1* gene following clonal isolation and expansion. This disrupts the reading frame of the gene, leading to a premature TGA (i.e., UGA in mRNA) stop codon (shown by an asterisk in the amino acid sequence) instead of an amino acid at position 295 in the KO-A and KO-B cell lines. Stop codons are also outlined with red rectangles for clarity. **B)** Effects of the 2-bp deletion shown in part A) on the downstream amino acid sequence of JAKMIP1. Additional UGA stop codons are found encoded towards the end of exon 5 (amino acid^313^ and amino acid^316^) and in exon 6 (amino acid^343^). Created with *Benchling* (Benchling [Biology Software]. (2022). Retrieved from https://benchling.com.). EV – empty vector; gRNA – guide RNA; KO-A – *JAKMIP1*-knockout A; KO-B – *JAKMIP1*-knockout B; WT – wild-type.

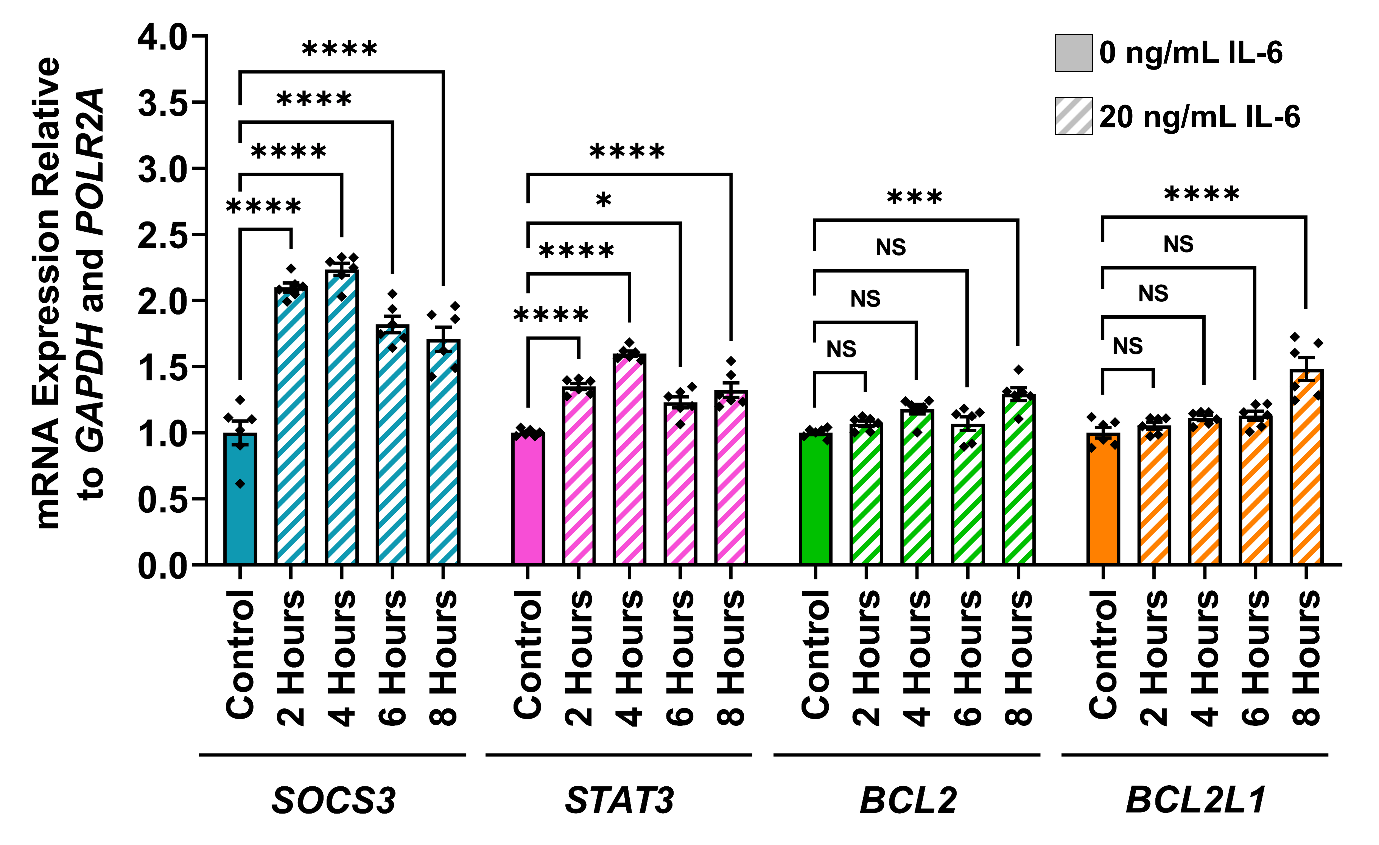

**Supplementary Figure 4 | Identifying STAT3-responsive genes induced by IL-6 treatment in SH-SY5Y cells.** Relative mRNA expression of typical STAT3-target genes (*SOCS3*, *STAT3*, *BCL2* and *BCL2L1*) in WT SH-SY5Y cells following treatment with 20 ng/mL IL-6 for various timepoints indicated (2-8 hours), measured by qRT-PCR. Cells treated with complete medium (i.e., culture medium was refreshed) for 8 hours were used as a control. All values are normalized to the non-IL-6-treated control. Values presented as mean ± SEM of N = 2 independent experiments, 3 replicates each. One-way ANOVA with Tukey’s HSD test for multiple comparisons. Statistical significance displayed against the non-IL-6-treated control; NS – not significant (P > 0.05); ***P < 0.001; ****P < 0.0001). ANOVA – analysis of variance; mRNA – messenger RNA; qRT-PCR – quantitative reverse transcription PCR; SEM – standard error of the mean; WT – wild-type.

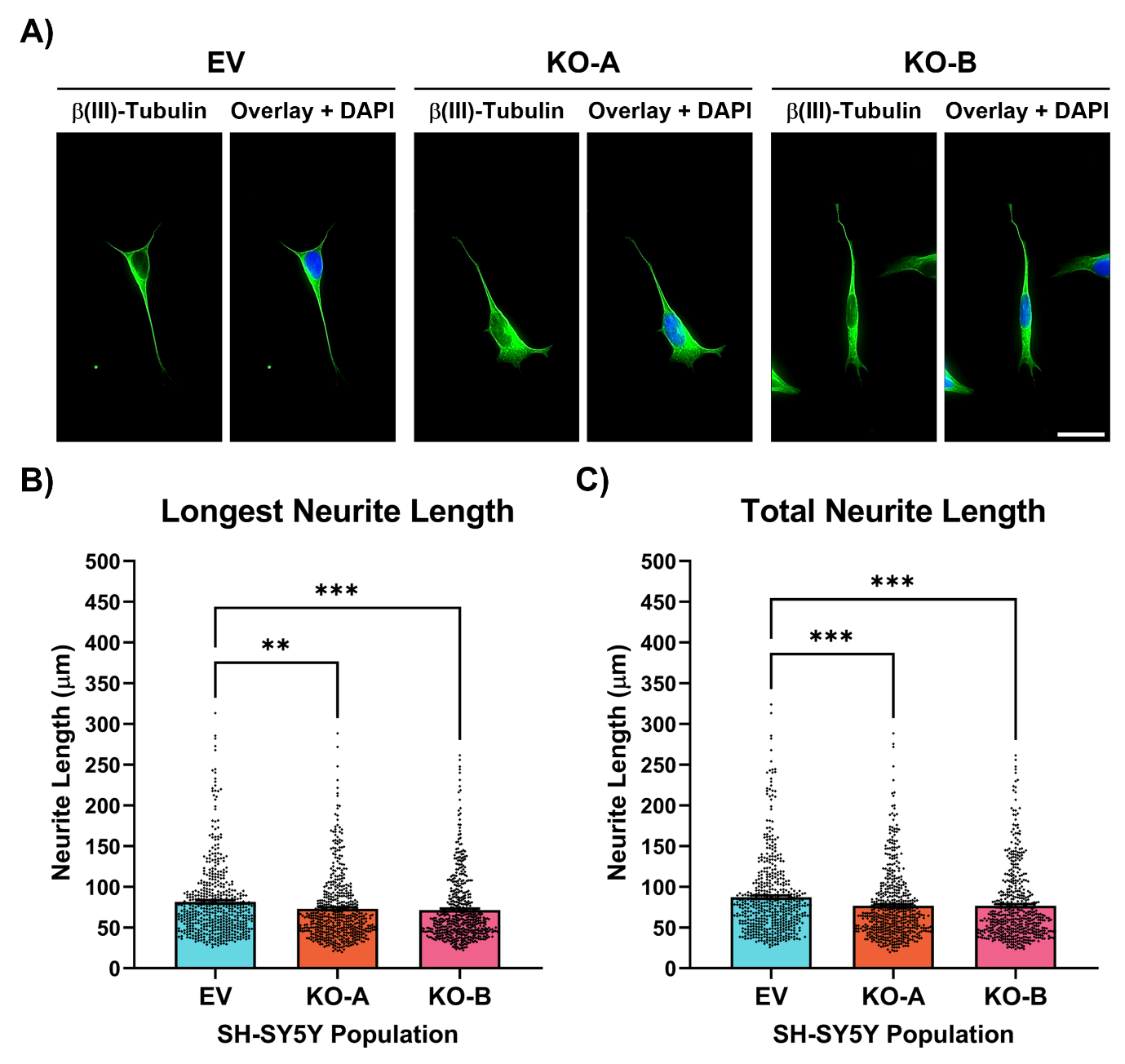

**Supplementary Figure 5 | JAKMIP1-/- SH-SY5Y cells display subtle deficits in neurite extension and neuritogenesis. A)** EV, KO-A and KO-B SH-SY5Y cells were differentiated on PDL-coated glass coverslips prior to fixation and immunostaining for β(III)-Tubulin (Alexa Fluor® 488; seen in green). Images representative of N = 3 independent differentiations, 2 separate coverslips per differentiation experiment and 515-522 SH-SY5Y cells traced per genotype. Scale bar = 50 μm. B) Measurement of the length of the longest neurite (LNL) produced by individual SH-SY5Y cells in part A). Kruskal-Wallis with Dunn’s multiple comparisons test for multiple comparisons. Statistical significance displayed against the EV control; NS – not significant (P > 0.05); ***P < 0.001; ****P < 0.0001. C) Measurement of the sum length of all neurites (TNL) produced by individual SH-SY5Y cells in part A). Kruskal-Wallis with Dunn’s multiple comparisons test for multiple comparisons. Statistical significance displayed as in B). EV – empty vector; KO-A – *JAKMIP1*-knockout A; KO-B – *JAKMIP1*-knockout B; PDL – poly-D-Lysine.

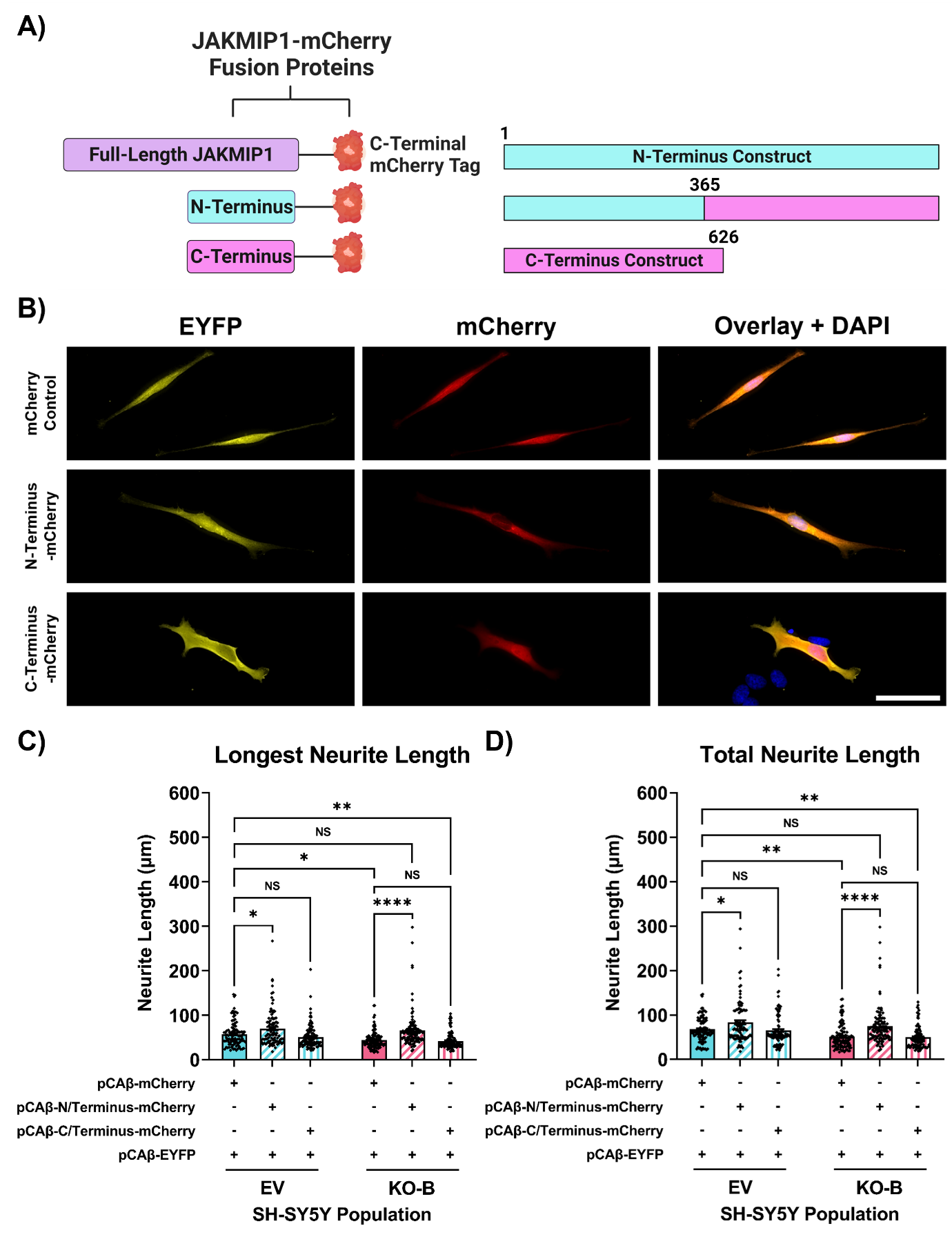

**Supplementary Figure 6 | The JAKMIP1 N-terminus is responsible for the neurite outgrowth-promoting effects of JAKMIP1. A)** Left: schematic representation of JAKMIP1-mCherry fusion proteins, including the full-length and truncated N- and C-termini domains. Right: Outline of amino acid composition of truncated JAKMIP1 constructs. The N-terminus is composed of amino acid^1-365^, and the C-terminus is composed of amino acid^365-626^. Created with BioRender.com. **B)** Representative fluorescence micrographs of differentiated EV and KO-B SH-SY5Y cells transfected with plasmid vectors encoding truncated JAKMIP1-mCherry fusion constructs (in conjunction with a separate vector encoding EYFP for neurite tracing) by Nucleofection™. Transfected SH-SY5Y cells were differentiated for 48 hours before fixation. Images representative of N = 3 independent differentiations, 2 separate coverslips per differentiation experiment and 97-110 SH-SY5Y cells traced per genotype/transfection combination. Scale bar = 50 μm. **C)** Measurement of the length of the longest neurite (LNL) produced by individual transfected SH-SY5Y cells in B). Two-way ANOVA with Tukey’s HSD test for multiple comparisons. Statistical significance displayed against the mCherry-OE EV control; NS – not significant (P > 0.05); *P < 0.05; **P < 0.01; ***P < 0.001; ****P < 0.0001. **D)** Measurement of the sum length of all neurites (TNL) produced by individual SH-SY5Y cells in B). Two-way ANOVA with Tukey’s HSD test for multiple comparisons. Statistical significance displayed as in C). ANOVA – analysis of variance; EV – empty vector; EYFP – enhanced yellow fluorescent protein; KO-B – *JAKMIP1*-knockout B; OE – overexpression; PDL – poly-D-Lysine.

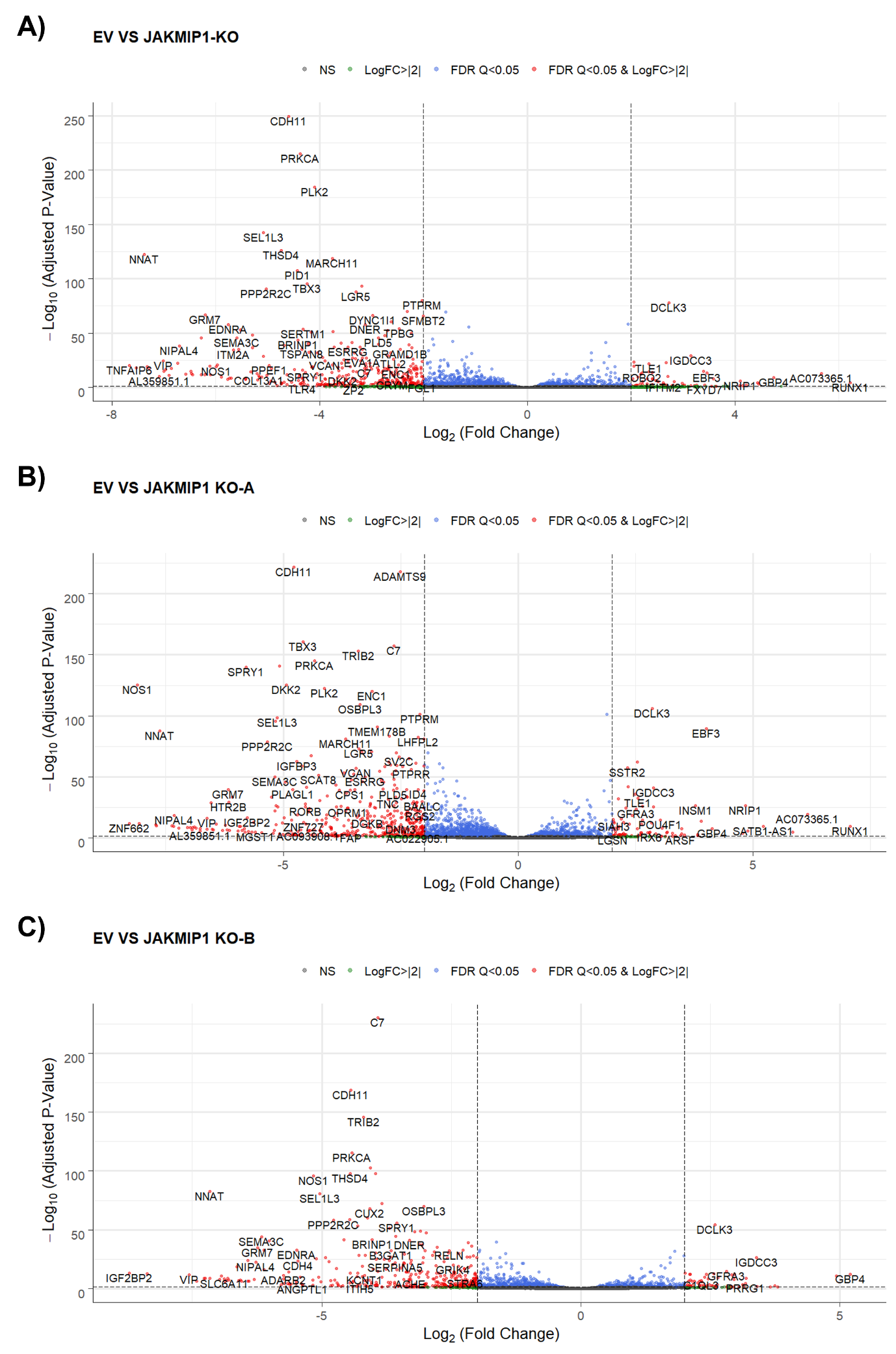

**Supplementary Figure 7 | Global gene expression changes in differentiated *JAKMIP1*^-/-^ SH-SY5Y cells.** Volcano plots for DEGs identified in **A)** both KO-A and KO-B combined; or **B)** KO-A only; or **C)** KO-B only compared to the EV control. Grey dots = non-significant DEGs (adjusted P-value > 0.05); green dots = significant DEGs (adjusted P-value < 0.05) with an absolute Log_2_FC ≥ 2; blue dots = significant DEGs (adjusted P-value < 0.05) with Q-value following Benjamini-Hochberg FDR correction ≤ 0.05; red dots = significant DEGs with both an absolute Log_2_FC ≥ 2 and Q-value following Benjamini-Hochberg FDR correction ≤ 0.05. DEG – differentially expressed gene; EV – empty vector control; FDR – false discovery rate; KO-A – *JAKMIP1*-knockout A; KO-B – *JAKMIP1*-knockout B; Log_2_FC – base 2 logarithm of differential gene expression fold change (FC).

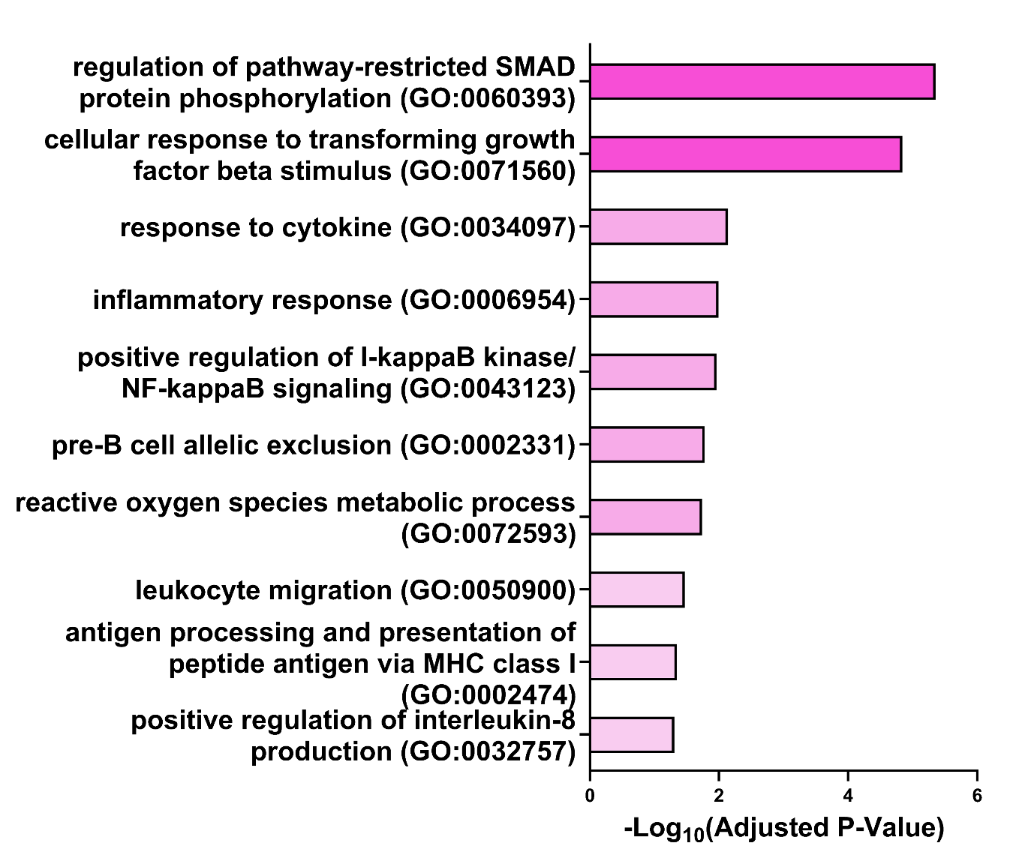

**Supplementary Figure 8 | Enrichment of cytokine signaling, inflammation and immune function-related GO Biological Process 2021 terms in downregulated DEGs.** Additional results of GO analysis performed on the separated downregulated DEGs relating to cytokine signaling, inflammation and immune cell function. 10 terms of interest are reported with the adjusted P-value following Benjamini-Hochberg FDR correction for multiple testing. Terms are sorted by significance of adjusted P-value. DEG – differentially expressed gene; FDR – false discovery rate.

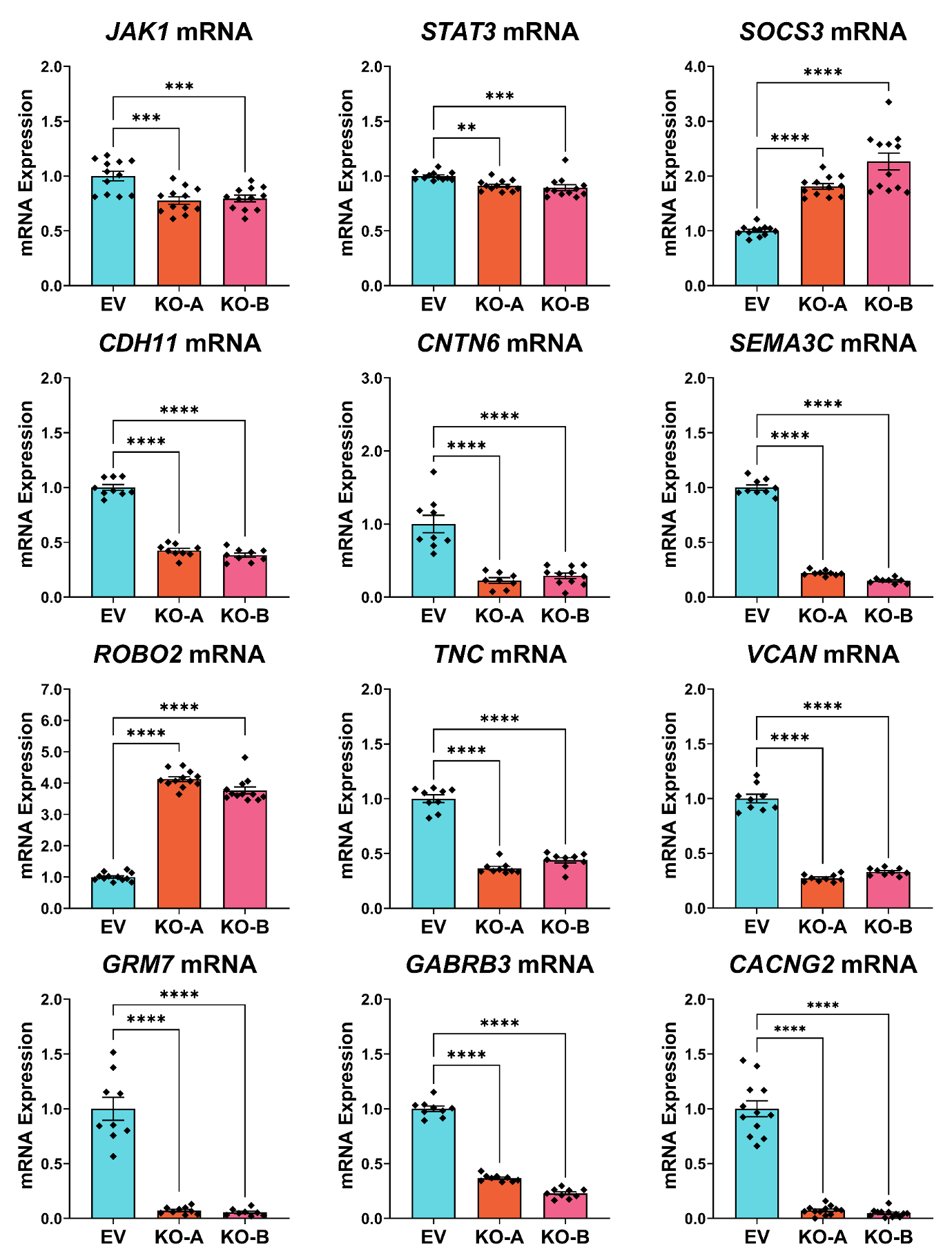

**Supplementary Figure 9 | Validation of differential gene expression analysis by qRT-PCR.** Relative mRNA expression of various identified DEGs from the RNA-seq in differentiated EV, KO-A and KO-B SH-SY5Y cells measured by qRT-PCR. DEGs fall into several broad categories, namely IL-6/JAK1/STAT signaling components (*JAK1*, *STAT3* and *SOCS3*); CAMs (*CDH11* and *CNTN6*); axon guidance ligands and receptors (*SEMA3C* and *ROBO2*); ECM components (*TNC* and *VCAN*); and synaptic receptors and ion channels (*GRM7*, *GABRB3* and *CACNG2*). *GAPDH* and *POLR2A* were used as housekeeping genes for relative mRNA expression quantification. All values are normalized to the EV control. Values presented as mean ± SEM of N = 3 independent experiments, 3-4 replicates each. Statistical significance displayed against the EV control; **P < 0.01; ***P < 0.001; ****P < 0.0001. One-way ANOVA with Tukey’s HSD test for multiple comparisons. DEG – differentially expressed gene; EV – empty vector control; KO-A – *JAKMIP1*-knockout A; KO-B – *JAKMIP1*-knockout B; mRNA – messenger RNA; RNA-seq – RNA sequencing; qRT-PCR – quantitative reverse transcription polymerase chain reaction; SEM – standard error of the mean.

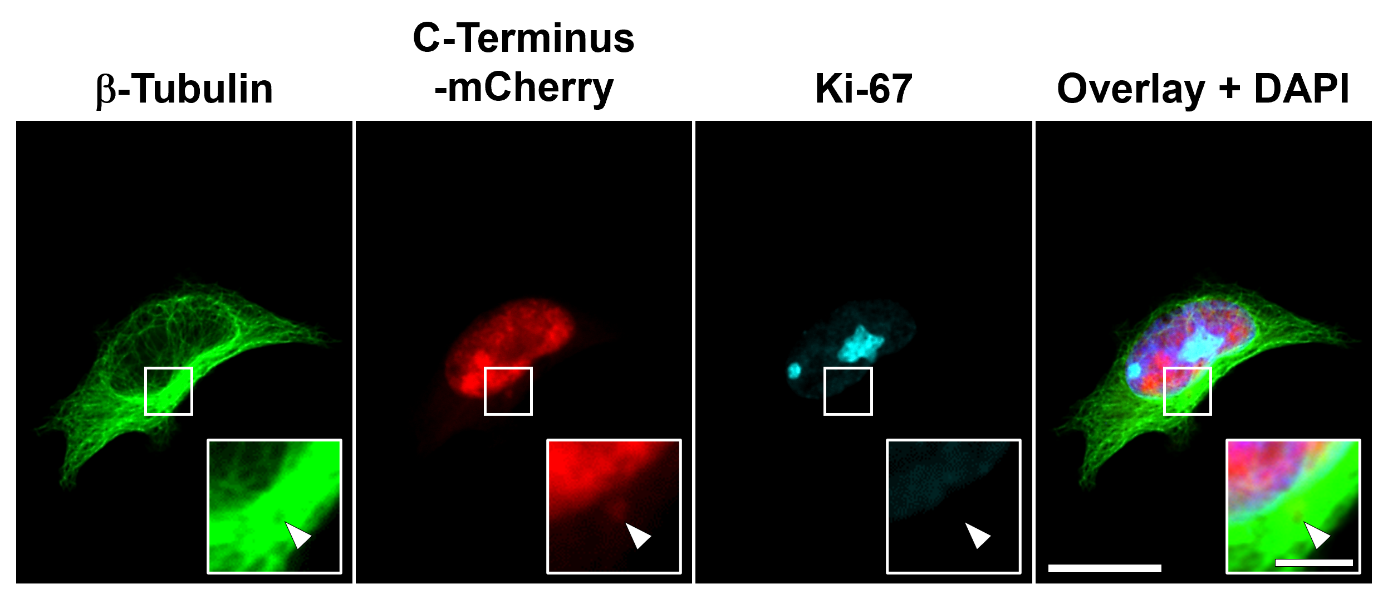

**Supplementary Figure 10 | The JAKMIP1 C-terminus can display centrosomal localization.** HEK-293 cells transfected with the mCherry-tagged JAKMIP1 C-terminus construct. Immunostaining for β-Tubulin and Ki-67. Scale bar = 20 μm. Insets show enlarged view of regions of interest outlined with white squares. Arrowhead indicates centrosomal localization of the JAKMIP1 C-terminus construct observed in a subset of transfected cells. Inset scale bar = 5 μm.

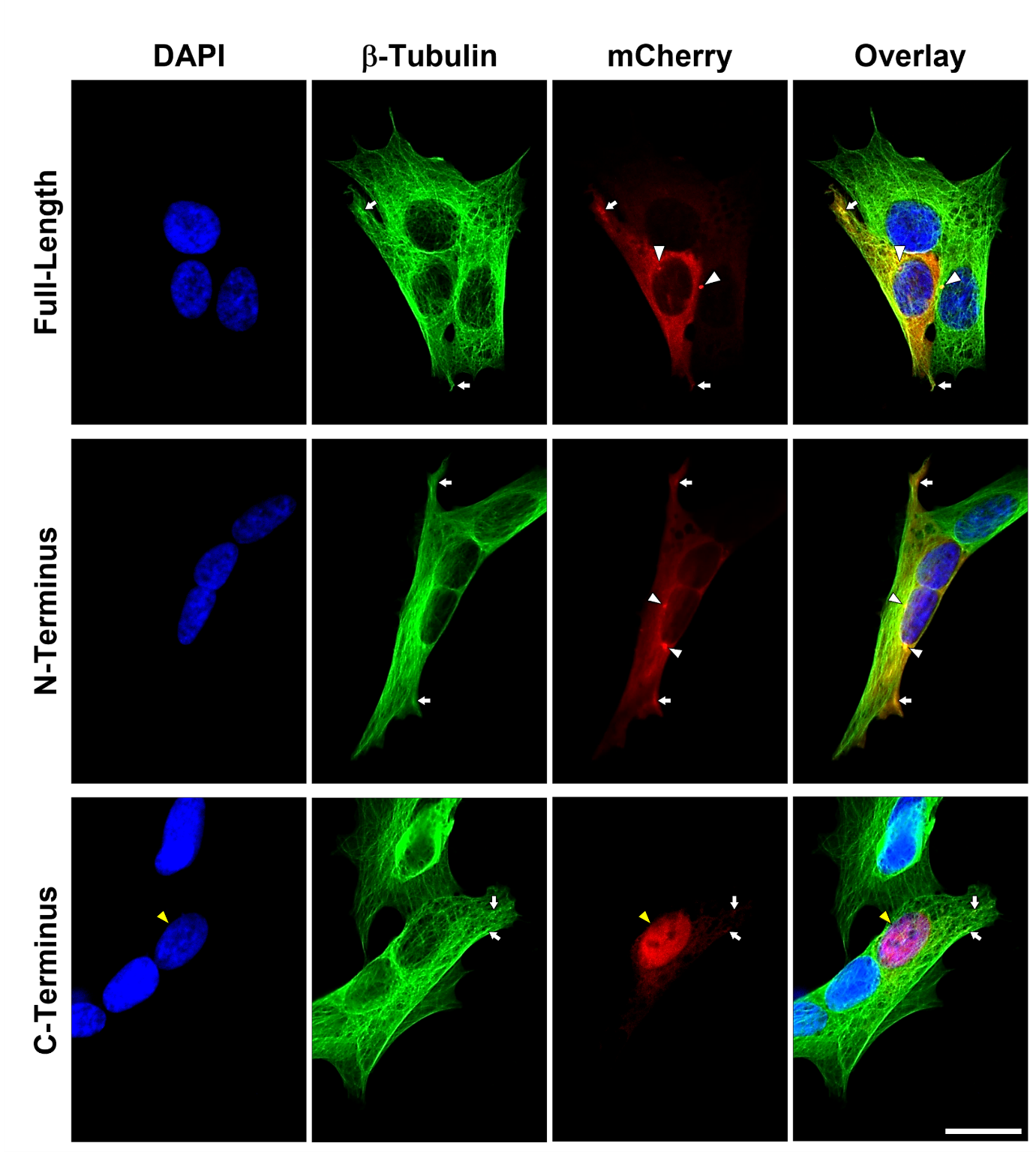

**Supplementary Figure 11 | The JAKMIP1 C-terminus displays predominantly nuclear localization.** Representative fluorescence micrographs of SH-SY5Y cells transfected with full-length or truncated JAKMIP1 constructs tagged with mCherry (using jetOPTIMUS®). Images representative of N = 3 independently fixed plates of cells, 1 separate coverslip stained per plate. Transfected cells were fixed 48 hours post transfection. Immunostaining for β-Tubulin (Alexa Fluor® 488; seen in green). White arrowheads indicate juxtanuclear puncta visible in both the full-length and N-terminus constructs; white arrows indicate co-localization with microtubules; and yellow arrowheads indicate nuclear localization of the C-terminus construct. Scale bar = 20 μm. PDL – poly-D-Lysine; WT – wild-type.

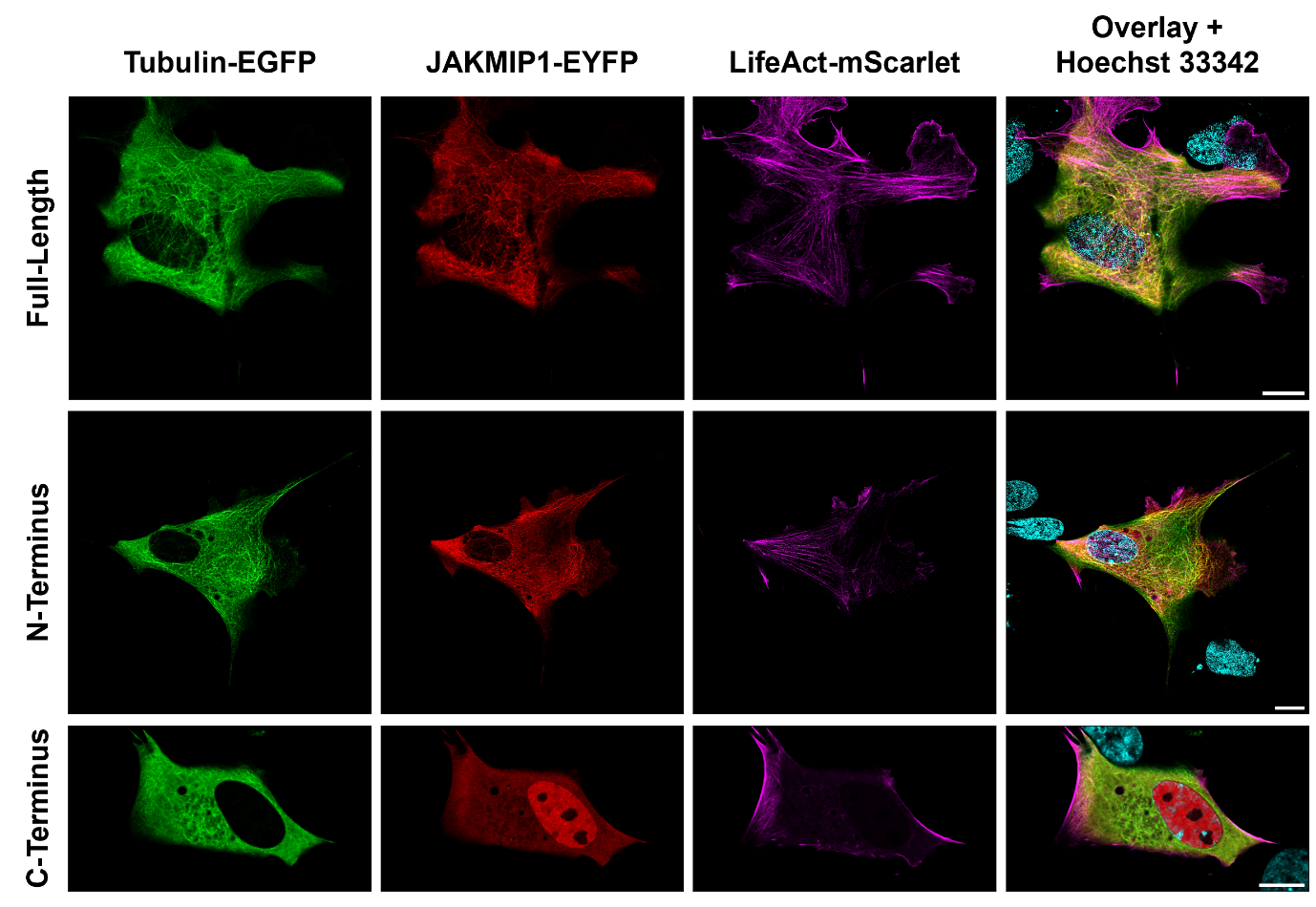

**Supplementary Figure 12 | Live-cell imaging confirms nuclear localization of the JAKMIP1 C-terminus.** HEK-293 cells transfected with plasmid vectors encoding EGFP-tagged Tubulin (green), EYFP-tagged full-length or truncated JAKMIP1 constructs (red) and mScarlet-tagged LifeAct (magenta) with Lipofectamine™ LTX. Nuclei were counterstained with 50 ng/mL Hoechst 33342 (cyan) for at least 30 minutes prior to imaging. Images captured with a Leica STELLARIS 8 confocal microscope by Paul McCormick. Scale bars = 10 μm.

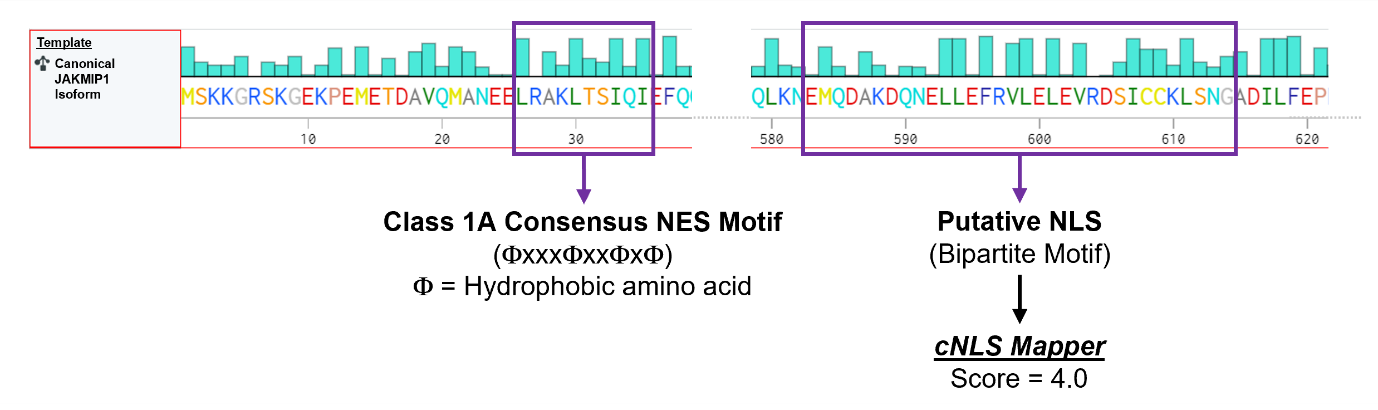

**Supplementary Figure 13 | Putative nuclear localization and export sequences in JAKMIP1.** Amino acid sequence of JAKMIP1. Cyan bars above each amino acid represent the hydrophobicity of the residue (longer bar = more hydrophobic). *cNLS Mapper* (available at: https://nls-mapper.iab.keio.ac.jp/cgi-bin/NLS_Mapper_form.cgi) identifies a putative bipartite NLS motif at the C-terminus of JAKMIP1 (located at amino acid^583-614^) with a score of 4.0. A class 1A consensus NES motif at the N-terminus of JAKMIP1 (located at amino acid^26-35^). Created with *Benchling* (Benchling [Biology Software]. (2022). Retrieved from https://benchling.com.). Φ – hydrophobic amino acid; NES – nuclear export sequence; NLS – nuclear localization sequence.

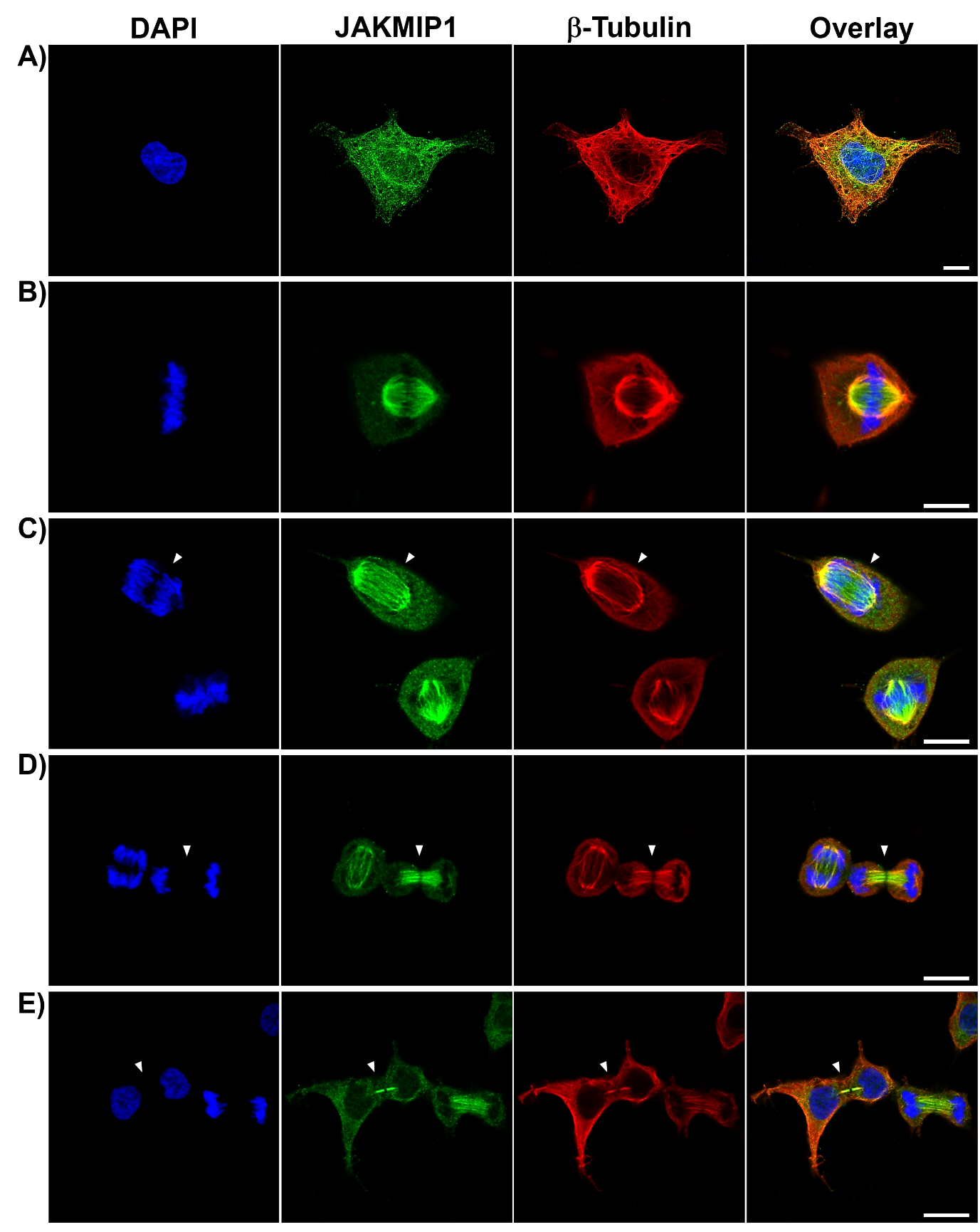

**Supplementary Figure 14 | JAKMIP1 is localized to spindle microtubules during mitosis.** HEK-293 cells were cultured on PDL-coated coverslips prior to fixation with PFA and immunostaining for JAKMIP1 (Alexa Fluor® 488; seen in green) and β-Tubulin (Alexa Fluor® 568; seen in red). Scale bars = 10 μm. **A)** Non-dividing HEK-293 cell in interphase. **B)** HEK-293 cell in metaphase. **C)** HEK-293 cell in early anaphase (indicated by arrowhead). **D)** HEK-293 cells in late anaphase (indicated by arrowhead. **E)** HEK-293 cells in late telophase prior to completion of cytokinesis (indicated by arrowhead). PDL – poly-D-Lysine; PFA – paraformaldehyde.

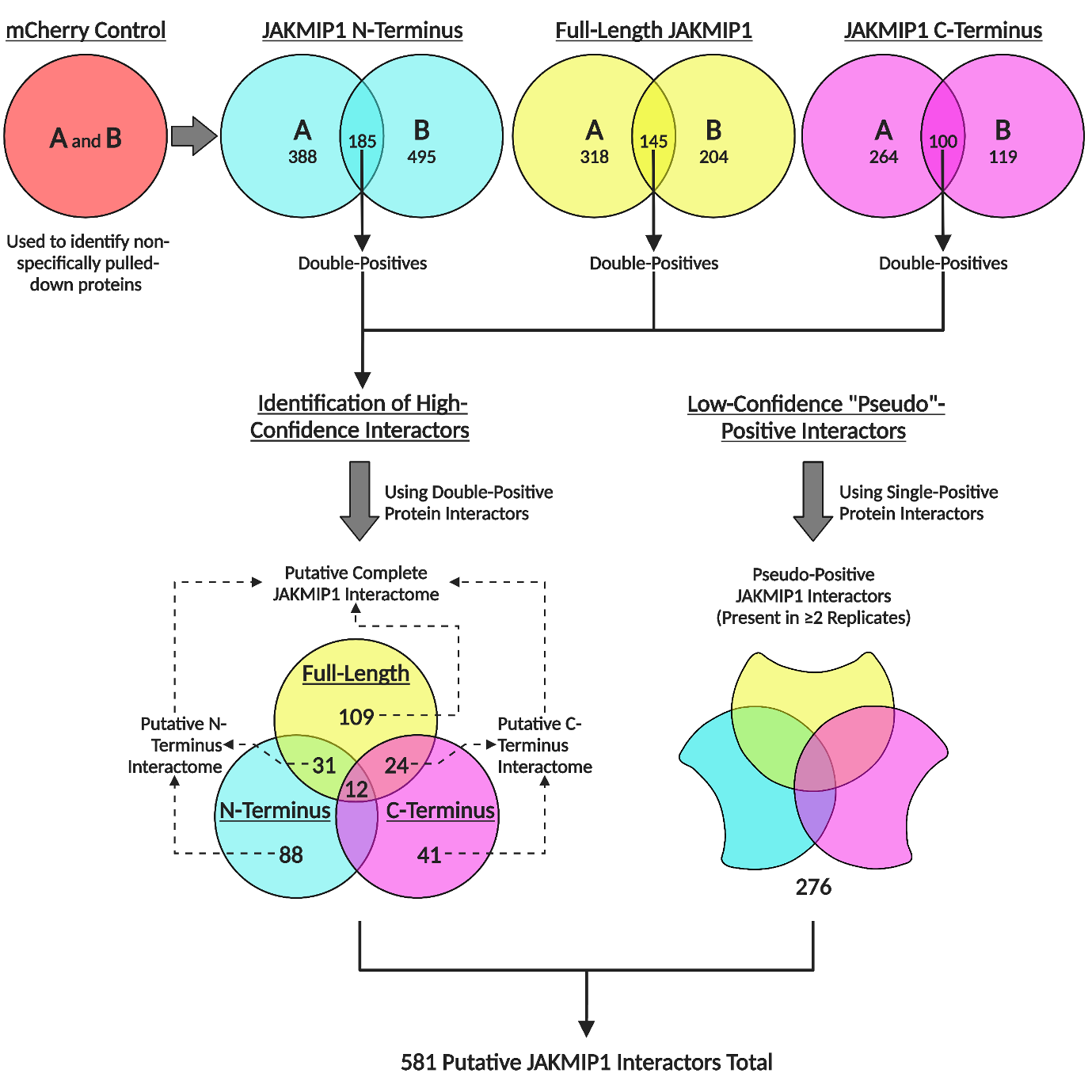

**Supplementary Figure 15 | Identification of putative JAKMIP1-interacting proteins from IP-MS data.** Overview of IP-MS analysis results, illustrating how the final set of 581 putative JAKMIP1-interacting proteins were identified. Created with BioRender.com.

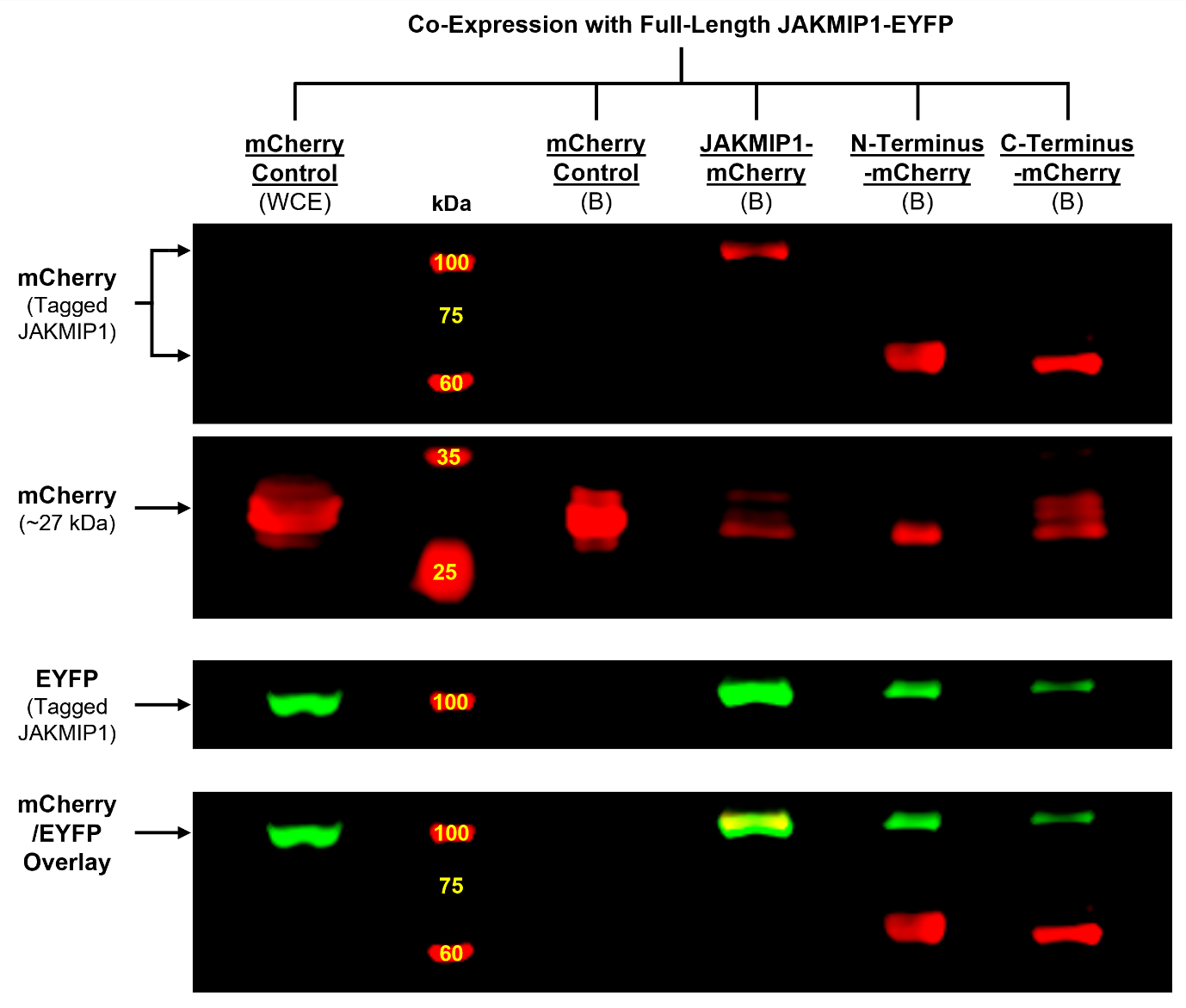

**Supplementary Figure 16 | JAKMIP1 can dimerize at both its N- and C-termini.** HEK-293 cells were transfected (using Lipofectamine™ LTX) with plasmid vectors encoding mCherry-tagged JAKMIP1 constructs in conjunction with an EYFP-tagged JAKMIP1 construct, allowed to express fusion proteins for 48 hours and then lysed for co-IP. JAKMIP1-EYFP/mCherry-OE SH-SY5Y cells were used as a control. Immunoprecipitation of mCherry-containing protein complexes was carried out using RFP-Trap® Agarose beads. Images representative of N = 2 independent samples generated from separate co-IP experiments; each sample subjected to Western blotting once. Lanes indicative of bound (“B”) fractions from cells expressing JAKMIP1-EYFP in conjunction with mCherry or JAKMIP1-mCherry constructs as indicated above each lane. The whole-cell extract (“WCE”) of JAKMIP1-EYFP/mCherry-OE HEK-293 cells was included as a positive control for protein-of-interest detection. Each fraction indicated was loaded into wells of polyacrylamide gels for SDS-PAGE. PVDF membranes were sequentially probed for EYFP and mCherry. JAKMIP1-EYFP is co-precipitated by the RFP-Trap® Agarose beads in HEK-293 cells expressing all the JAKMIP1-mCherry fusion constructs but not mCherry only. An overlay of the mCherry/EYFP signal is provided to demonstrate the similarity in molecular weight of the JAKMIP1-mCherry and JAKMIP1-EYFP fusion proteins. co-IP – co-immunoprecipitation; EYFP – enhanced yellow fluorescent protein; OE – overexpression; PVDF – polyvinylidene difluoride; SDS-PAGE – sodium dodecyl sulfate polyacrylamide gel electrophoresis.

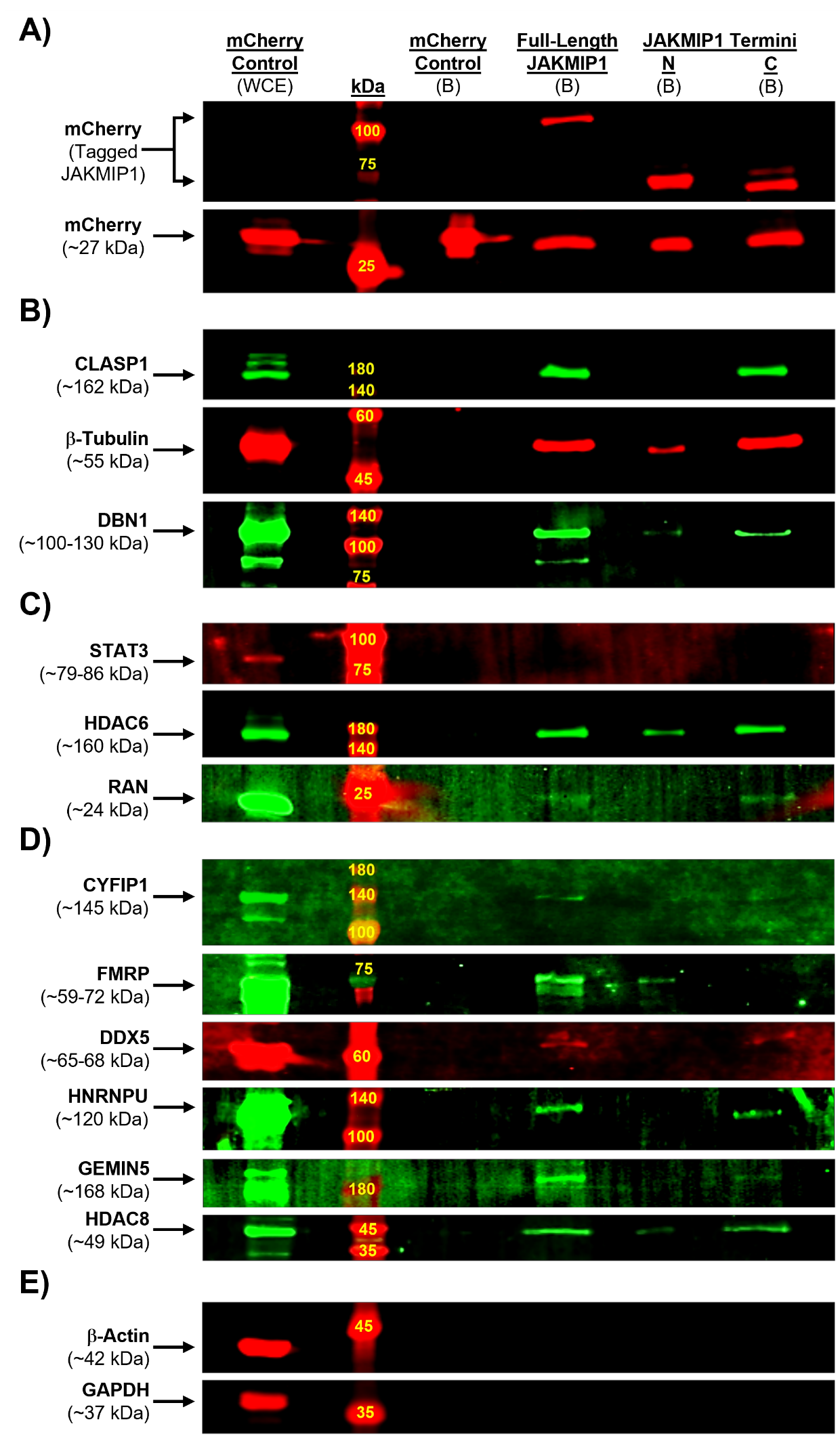

**Supplementary Figure 17 | Confirmation of putative JAKMIP1 interactors by co-IP.** Known and novel JAKMIP1-interacting proteins from IP-MS data were validated by co-IP. WT SH-SY5Y cells were transfected (using Lipofectamine™ LTX) with plasmid vectors encoding mCherry-tagged JAKMIP1 constructs and allowed to express fusion proteins for 48 hours prior to lysis and co-IP. mCherry-OE SH-SY5Y cells were used as a control. N = 2-3 independent samples generated from separate co-IP experiments; each sample subjected to Western blotting 1-2 times. Lanes indicative of bound (“B”) fractions from cells expressing each construct or mCherry only. The whole-cell extract (“WCE”) of mCherry-OE cells was included as a positive control for protein-of-interest detection. Proteins of interest targeted by primary antibodies are indicated to the left of the images. **A)** All JAKMIP1 constructs can be detected by mCherry at the appropriate molecular weight. **B)** All constructs interact with b-Tubulin and DBN1 but only the full-length and C-terminus pulls down CLASP1. **C)** No detectable interaction between the JAKMIP1 constructs and STAT3, but all constructs are capable of interacting with HDAC6, and RAN co-precipitates with the full-length or C-terminus constructs. **D)** CYFIP1 interacts with the full-length JAKMIP1 construct only, whereas FMRP interacts with the full-length and N-terminus constructs; and DDX5, HNRNPU and GEMIN5 interact with the full-length and C-terminus constructs. HDAC8 co-precipitates with all three constructs. **E)** No detectable contamination of the bound fractions by b-Actin or GAPDH, known non-JAKMIP1-interacting proteins. Co-IP – co-immunoprecipitation; IP-MS – immunoprecipitation mass spectrometry; OE – overexpression; SDS-PAGE – sodium dodecyl sulfate polyacrylamide gel electrophoresis; WT – wild-type.

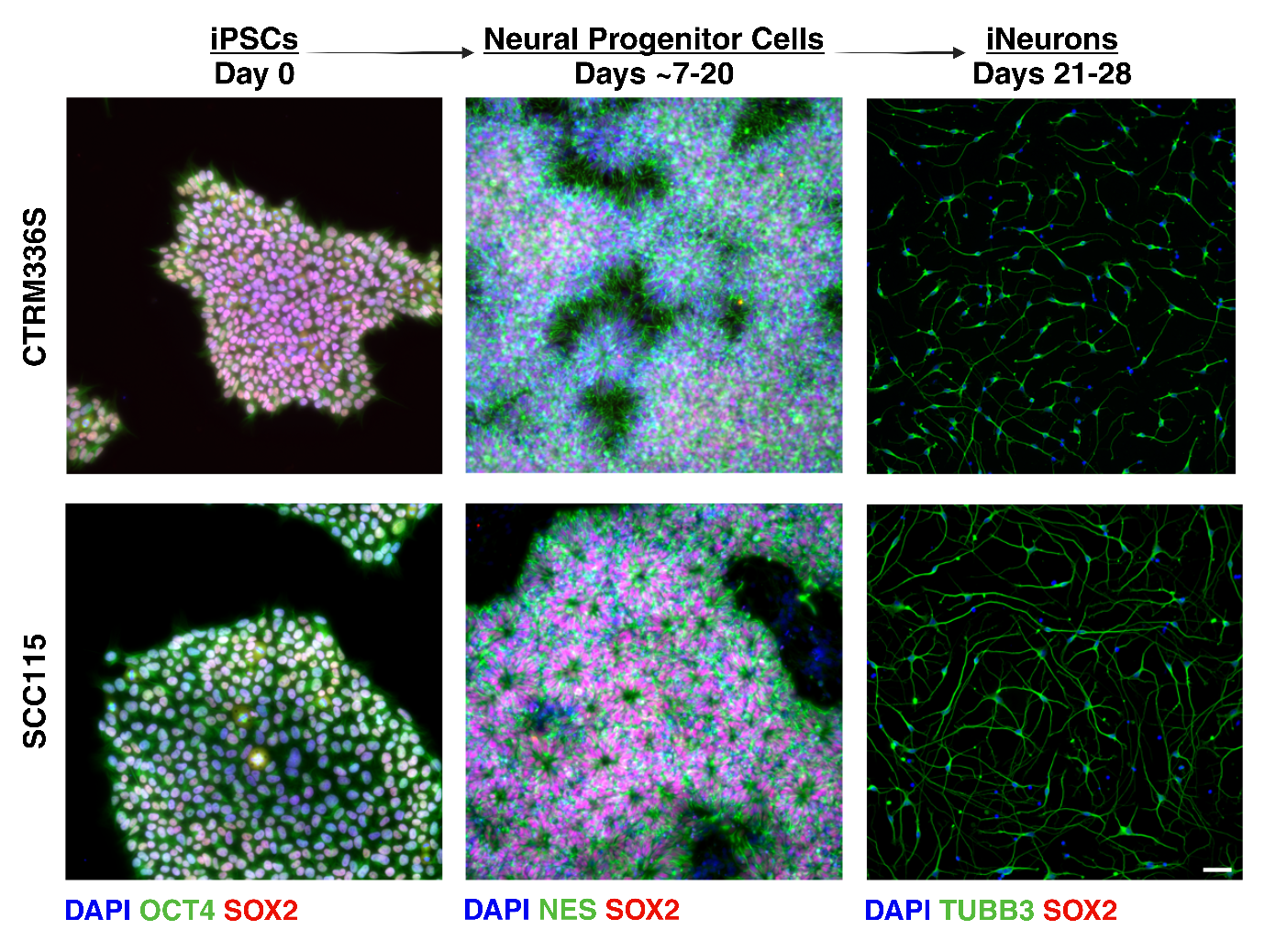

**Supplementary Figure 18 | Confirmation of successful neuralization of CTRM336S and SCC115 hiPSCs into iNeurons.** Representative fluorescence micrographs of CTRM336S and SCC115 cells fixed and stained for various markers for the neuralization process: prior to neuralization as hiPSCs (OCT4 (green) and SOX2 (red)), neural progenitor cells (NES (green) and SOX2 (red)) and iNeurons (TUBB3). hiPSC – human induced pluripotent stem cell; iNeuron – hiPSC-derived neuron. Scale bar = 100 μm.

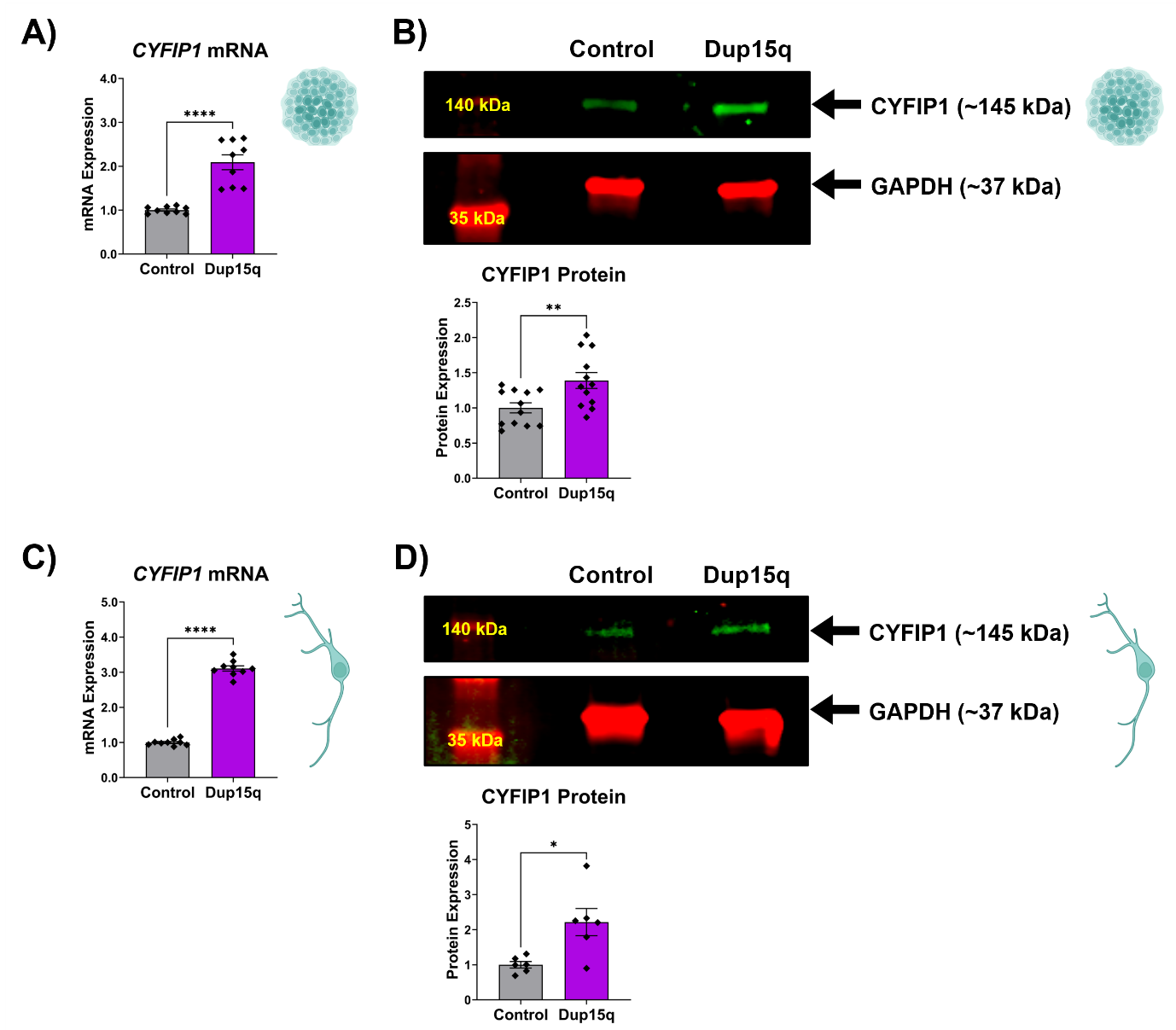

**Supplementary Figure 19 | *CYFIP1* expression is raised in Dup15q hiPSCs and iNeurons.** Relative mRNA and protein expression of *CYFIP1* in control and Dup15q **A-B)** hiPSCs and **C-D)** iNeurons. **A)** *CYFIP1* mRNA expression in control and Dup15q hiPSCs measured by qRT-PCR. Values presented as mean ± SEM of N = 3 independent experiments, 2-3 replicates each. One-way ANOVA with Tukey’s HSD test for multiple comparisons. Statistical significance displayed against the control; *P < 0.05; **P < 0.01; ****P < 0.0001. **B)** CYFIP1 protein expression in control and Dup15q hiPSCs measured by Western blotting. Values presented as mean ± SEM of N = 4 independent experiments, 3 replicates each. One-way ANOVA with Tukey’s HSD test for multiple comparisons. Statistical significance displayed as in A). **C)** *CYFIP1* mRNA expression in control and Dup15q iNeurons measured by qRT-PCR. Values presented as mean ± SEM of N = 3 independent experiments, 2-3 replicates each. One-way ANOVA with Tukey’s HSD test for multiple comparisons. Statistical significance displayed as in A). **D)** CYFIP1 protein expression in control and Dup15q iNeurons measured by Western blotting. Values presented as mean ± SEM of N = 3 independent experiments, 2 replicates each. One-way ANOVA with Tukey’s HSD test for multiple comparisons. Statistical significance displayed as in A). ANOVA – analysis of variance; Dup15q – chromosome 15q-duplication syndrome; mRNA – messenger RNA; qRT-PCR – quantitative reverse transcription PCR; SEM – standard error of the mean.

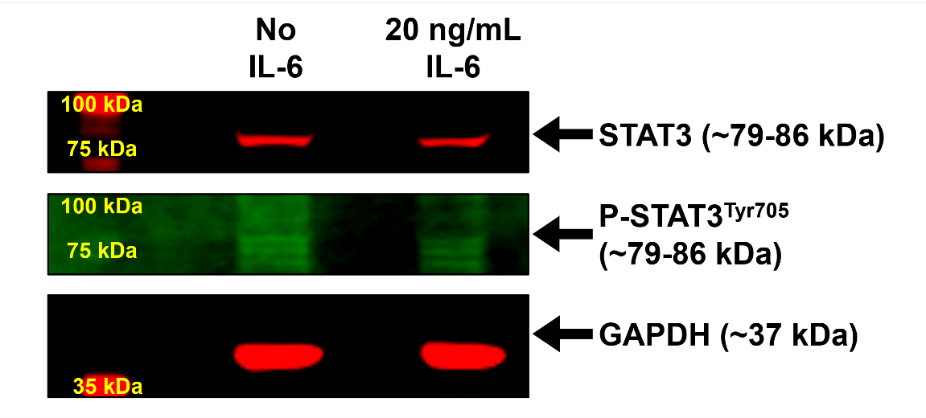

**Supplementary Figure 20 | IL-6 does not induce STAT3 (Tyr^705^) phosphorylation in iNeurons.** CTRM336S iNeurons were treated with 20 ng/mL IL-6 or B-27™ medium for 30 minutes prior to lysis for protein extraction. Western blotting was performed to measure STAT3 (Tyr^705^) phosphorylation relative to the GAPDH loading control. Note that no clear band for P-STAT3^Tyr705^ is visible (the brightness has been raised to the point of revealing background autofluorescence). Tyr^705^ – tyrosine residue at amino acid position 705.

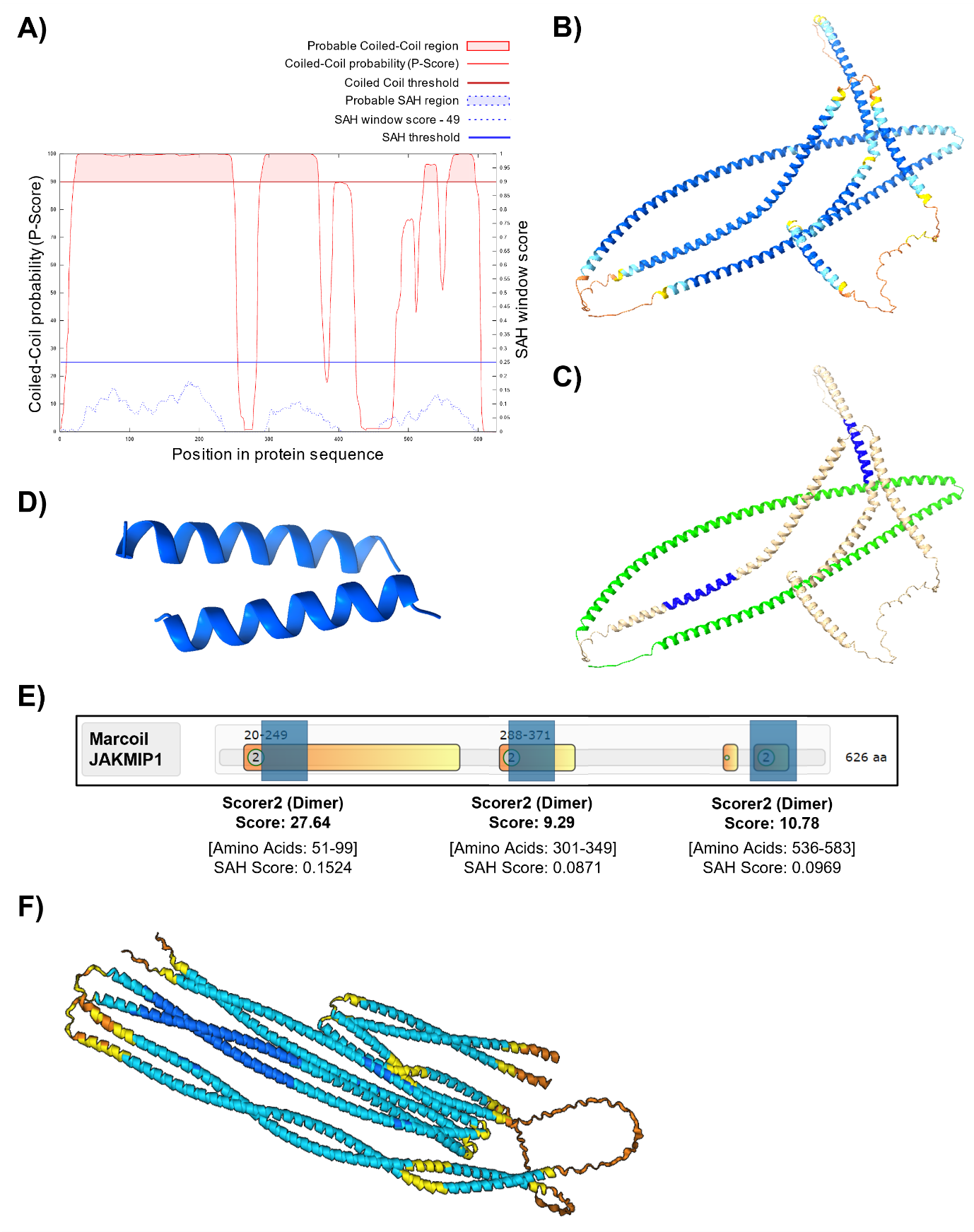

**Supplementary Figure 21 | JAKMIP1 is composed of highly helical structures forming coiled coils.** **A)** *Marcoil*-based prediction of coiled coils in the canonical JAKMIP1 isoform performed through *Waggawagga* (available at: https://waggawagga.motorprotein.de/). Results from *Marcoil* suggest JAKMIP1 is likely to have three main groups of coiled coils (red shaded areas in upper region of the graph) located at its N-terminus (amino acids^20-249^), RNA-binding domain (amino acids^288-371^) and C-terminus (amino acids^523-541^ and amino acids^556-595^). **B)** Predicted three-dimensional model structure of JAKMIP1 obtained from *AlphaFold* (available at: https://alphafold.ebi.ac.uk/), which are largely composed of long helices. Confidence of the model is reflected by the color of the amino acid residues (blue = very high confidence; cyan = confident; yellow = low confidence; orange = very low confidence). **C)** The N-terminal coiled coil domain and leucine zipper motif of the JAKMIP1 structure from B), highlighted in green and blue respectively. **D)** Predicted model of the leucine zipper region highlighted in C) by *AlphaFold 3* (available at: https://www.alphafoldserver.com/). **E)** *Marcoil*-based prediction of JAKMIP1 dimerization at three predicted coiled coil regions from part A). High *Scorer2* scores indicates that JAKMIP1 may form dimers mediated by the coiled coils highlighted in orange-yellow. For reference, MYH2, a cytoskeletal protein with known coiled-coils and is established to dimerize, has *Scorer2* scores between 6.83-18.55. SAH scores for the regions highlighted in blue are also provided. **F)** Predicted model of a JAKMIP1-JAKMIP1 dimer by *AlphaFold 3*, with colors of amino acid residues meaning the same as in B). The predicted structure demonstrates a well-fitting JAKMIP1 dimer with an even more coiled structure than singular JAKMIP1, reinforcing dimerization and the presence of functional coiled coils. SAH – single alpha helix.

Supplementary Tables (File S2)

Supplementary tables are contained in File S2, which can be accessed online in the *figshare* repository at the following DOI: 10.6084/m9.figshare.30402832.
